# The sandarazols are cryptic and structurally unique plasmid encoded toxins from a rare myxobacterium

**DOI:** 10.1101/2020.10.06.323741

**Authors:** Fabian Panter, Chantal D. Bader, Rolf Müller

## Abstract

Soil dwelling bacteria such as myxobacteria defend themselves by using secondary metabolites to inhibit growth of competing microorganisms. In this work we describe a new plasmid found in *Sandaracinus* sp. MSr10575 named pSa001 spanning 209.7 kbp that harbors a cryptic secondary metabolite biosynthesis gene cluster (BGC). Activation of this BGC by homologous recombination mediated exchange of the native promoter sequence against a vanillate inducible system led to production and subsequent isolation and structure elucidation of novel secondary metabolites, the sandarazols A-G. The sandarazol structure contains intriguing features such as an α-chlorinated ketone, an epoxyketone and a (2R)-2-amino-3- (N,N-dimethylamino)-propionic acid building block. In depth investigation of the underlying biosynthetic machinery led to a concise biosynthetic model for the new compound family, including several uncommon biosynthesis steps. The chlorinated congener sandarazol C shows an IC_50_ value of 0.5 µM against HCT 116 cells and a MIC of 14 µM against *Mycobacterium smegmatis*, which points at the sandarazol’s potential function as defensive secondary metabolites or toxins. The sandarazols’ BGC location on pSa001 is one of the very few example of large multimodular BGCs on a replicative plasmid, whose existence points at the mechanism of horizontal gene transfer events of entire multimodular BGCs to exchange chemical warfare capabilities between bacterial species.

## Introduction

Bacterial antibiotics resistance genes cause antimicrobial resistance (AMR) in pathogens thus leading to enormous challenges in the treatment of infectious diseases in the clinics. ^1^ AMR genes are often encoded on plasmids, horizontally transferrable elements employed by bacteria to survive antibiotics treatment and effectively spread AMR. ^2^ In addition to that, AMR genes co-localized with antibiotics biosynthesis gene clusters (BGCs) are important for self-resistance during antibiotics production in bacteria capable of antibiotics biosynthesis. Similarly to the plasmid mediated exchange of antibiotic resistance, bacteria are able to gain an advantage over competing bacteria by exchanging secondary metabolite BGCs in horizontal gene transfer events. ^3^ The mechanism for plasmid mediated exchange of secondary metabolite BGCs is known for plasmid encoded toxins, with the anthrax causing *Bacillus anthracis* as the most prominent example. ^4^ The *Bacillus* plasmid is shared between different bacilli and thereby transmits the anthrax toxin virulence factor. Exchange mechanisms of such plasmids encoding toxins enable the respective bacteria to acquire the ability for chemical warfare. As bacteria are competing with other microbes for nutrients in their ecological niches, the encoded chemical warfare molecules optimized by evolution are attractive targets in the search for novel bioactive natural products. For actinobacteria it has already been shown that defensive secondary metabolites can be encoded as multimodular BGCs on autonomously replicating plasmids. A BGC responsible for production of the highly cytotoxic mycolactones for example is also encoded on a large plasmid. ^5^ This cytotoxin is produced by *Mycobacterium ulcerans* and released during the infection process of human skin, while the bacterium feeds on skin cells. Streptomycetes can also host plasmid borne antibiotics BGCs as exemplified by the esmeraldins, a series of phenazine antibiotics. ^6^ With respect to the total output of natural products and natural product diversity, bacteria have emerged as major players as they are prolific producers of biologically active secondary metabolites. ^7,8^ Besides the well-described bacterial phyla such as actinobacteria, firmicutes and cyanobacteria that are responsible for the majority of identified biologically active natural products to date, Gram-negative proteobacteria such as myxobacteria have also shown great promise for the discovery of novel bioactive secondary metabolites. ^9–11^ Especially novel bacterial genera among the myxobacteria, such as the recently identified genus *Sandaracinus* studied herein, hold promise for finding interesting novel natural products chemistry. ^12^ In recent years, availability of cheap and reliable DNA sequencing technologies along with the development of BGC prediction tools such as antiSMASH depict the theoretical genetically encoded bacterial secondary metabolome as to date largely underexploited. ^13,14^ However, many of the BGCs that can be identified *in-silico* are ‘cryptic’ under laboratory cultivation conditions, meaning the production of the corresponding secondary metabolite is completely repressed or remains below the detection limit of modern analytical tools such as liquid chromatography (LC) coupled mass spectrometry (MS) instrumentation. Activating such cryptic clusters by genetic tools holds promise for discovering new chemistry, but currently remains a non-automatable, labor-intensive process. ^15^ Therefore, a strong focus on rational prioritization of BGCs to be activated by heterologous expression or gene cluster overexpression using heterologous promoters, remains crucial to focus discovery efforts on BGCs encoding for bioactive natural products. ^16^ Methods employed for BGC prioritization so far include expected target-guided mining for potential self-resistance genes and exploring biosynthetic gene cluster complexity for modular BGC architectures as well as the presence of certain tailoring enzymes such as halogenases or epoxidases. ^17^ Still, presence or absence of a BGC on a plasmid has not yet been used as a means for BGC prioritization. Based on the reported examples mentioned above, we reasoned that compounds originating from BGCs encoded on autonomously replicating plasmids seem to have a high likelihood to show biological activity. They thus represent a prime target in the search for novel bioactive natural products for anti-infective research and oncology and should be prioritized in activation efforts of cryptic BGCs.

As part of our ongoing efforts to isolate taxonomically diverse myxobacteria we isolated a novel myxobacterial strain belonging to the *Sandaracinus* clade. Surprisingly, the strain’s genome comprises a circular 209.7kbp plasmid called pSa001 that features a large polyketide synthase (PKS) non-ribosomal peptide synthetase (NRPS) hybrid BGC in its sequence. In this work, we show that activation of this BGC by genetic engineering of this stain leads to production of several derivatives of the sandarazols. These compounds turned out to be strong and chemically novel toxins biosynthesized by a series of intriguing biosynthetic steps.

## Results and Discussion

### Biosynthetic Gene Cluster Identification and Activation

Myxobacteria are Gram-negative *δ*-proteobacteria with a rich and diverse secondary metabolism. ^10,11^ A survey of known and unknown natural products in 2300 myxobacteria showed structural novelty to be clearly correlated to phylogenetic distance and thus heavily reliant on in depth investigation of novel myxobacterial genera. ^12^ We thus chose to dive into the secondary metabolome of a myxobacterial strain called MSr10575 that was recently isolated in-house. The strain belongs to the *Sandaracinus* genus, a rare myxobacterial genus little studied for secondary metabolism with the type strain *Sandaracinus amylolyticus* NOSO-4T. ^18^ This strain was the only member in the Sorangineae clade until MSr10575 was isolated and provided the indiacens A and B, two prenylated indole secondary metabolites. ^19^ We chose to sequence the genome of the second member of the *Sandaracinus* family (strain MSr10575) using single molecule real time sequencing technology and annotated the BGCs in its sequence with on our in-house antiSMASH server to obtain an overview about the strains’ secondary metabolite production potential. ^13^ Sequencing coverage analysis and genome assembly revealed that the genome of MSr10575 consists not only of a bacterial chromosome of 10.75 Mbp but also of an autonomously replicating plasmid of 209.7 kbp, which we named pSa001 (see SI). As sequencing coverage per base on the pSa001 plasmid is twice as large as observed for the bacterial chromosome, we assume a median value of two pSa001 copies per MSr10575 cell (see SI). The overall GC content of the plasmid is 70% and the codon usage bias favors GC rich codons over AT rich codons for the same amino acid at a rate of 3.6 to 1. As these values do not differ significantly from the corresponding parameters for the MSr10575 chromosome, the plasmid seems well adapted to its host strain (see SI). The presence of this plasmid is outstanding as autonomously replicating plasmids in myxobacteria are very rare. The only other example of a characterized myxobacterial plasmid is pMF1 from *Myxococcus fulvus*. ^20^ In comparison to pMF1, pSa001 is not only significantly larger, but also encodes a large type 1 *in-trans* acyl transferase (*trans*-AT) PKS NRPS hybrid BGC, which was named sandarazol (*szo*) BGC (see SI). Besides the *szo* BGC, the plasmid contains 5 ORFs encoding putative transposases and one ORF encoding a putative integrase indicating its ability to integrate into (or transfer parts of its sequence into) foreign bacterial genomes after conjugation. It might thus serve as a BGC shuttle vector for horizontal gene transfer of the BGC. Cultivation of wild type MSr10575 followed by extraction and liquid chromatography – tandem mass spectrometry (LC-MS^2^) analysis and GNPS based spectral networking of the bacterial metabolome did not reveal a family of secondary metabolites matching the BGC architecture on the plasmid in expected secondary metabolite size and fragmentation pattern (see SI). ^21^ We therefore assumed the plasmid borne *szo* BGC to be ‘cryptic’ as seen in many other cases such as the pyxidicyclines or taromycin. ^22,23^ All the coding regions in the *szo* cluster spanning from *szoA* to *szoO* are encoded on the same DNA strand and the intergenic regions seemed too small to contain important elements other than ribosome binding sites. We therefore reasoned that the *szo* cluster is encoded as a single transcriptional unit even though it spans 44.5 kbp. Activating the sandarazol BGC with a single promoter exchange in line with the experiments described in promoter exchange BGC activation in myxobacteria seemed therefore a reasonable strategy. ^22,24^ As we set out to create a plasmid for *szo* cluster overexpression by single crossover promoter exchange, we chose a plasmid with a pBelobac replication origin, as this would subsequently allow to extract the entire 209.7 kbp pSa001 and transfer it as replicative plasmid into *E. coli* for further investigations into the plasmids replication mechanism. ^25^ To investigate the secondary metabolite products of the BGC, we chose BGC activation in the native host MSr10575 first, as MSr10575 was assumed to be able to produce the corresponding small molecule product. We activated the *szo* cluster on the plasmid by promoter exchange against the vanillate promoter and repressor cassette, because this tool achieved overexpression of BGCs in several other myxobacteria including *Myxococcus xanthus* and *Pyxidicoccus fallax* (see SI). ^22,26^ Compared to other promoters, the vanillate promoter and repressor system shows tight control of BGC expression as well as strong secondary metabolite production upon BGC induction in myxobacteria.^27^

To exchange the promoter of the *szo* cluster, the first 2000 bp of the *szo* BGC were PCR-amplified, fused to a vanillate promoter and repressor cassette and ligated into the commercial pBeloBac11. This plasmid featuring a BeloBac origin and a kanamycin resistance in its backbone was named pBeloBac-TransAT (see SI). Promoter exchange of the *szo* clusters’ native promoter against the vanillate cassette is achieved by homologous recombination of pBelo-Bac-TransAT with the pSa001 plasmid after electroporation (Figure 1). MSr10575 clones harboring the recombined fused plasmid called pBeloBacSa001 were selected on 25 µg/mL kanamycin and genotypically verified by PCR (see SI). To obtain a comprehensive overview about the metabolic differences between MSr10575 wild type and MSr10575::pBeloBacSa001, analytical scale fermentation cultures were prepared in triplicates, extracted and subjected to UHPLC-qTOF analysis. The triplicate analyses were bucketed and all MSr10575 wild type derived liquid chro-matography – mass spectrometry (LC-MS) features were subtracted from the detected LC-MS features using principal component analysis (PCA).*(24)* These features were then used for selective acquisition of LC-MS^2^ data covering exclusively the mutant derived LC-MS features that are subsequently used for spectral networking in GNPS.^21^ We thus obtained spectral networks comprising all LC-MS features linked to activation of the sandarazol BGC (see Figure 1, SI).

**Figure 1.**
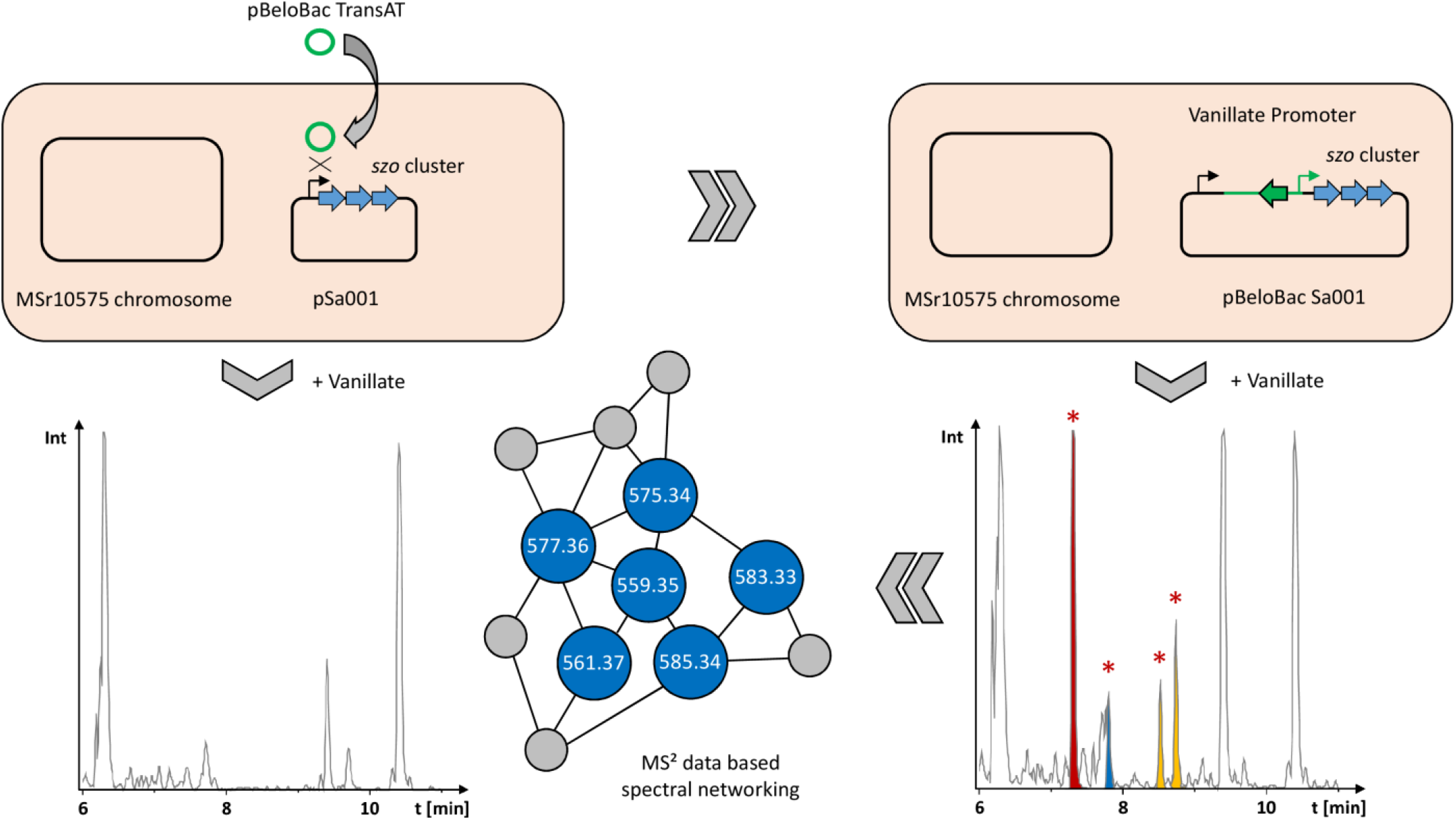
Schematic overview on the activation of the plasmid borne *szo* BGC in wild type *Sandaracinus* sp. MSr10575 by homologous recombination, the corresponding changes in the extracts’ LC-MS chromatograms and LC-MS^2^ based spectral networking for identification of the produced sandarazols.

### Isolation and structure elucidation

The novel secondary metabolites observed in this PCA-based analysis of LC-MS features were named the sandarazols. Besides sandarazol A (575.344 Da [M+H]^+^; C_30_H_47_N_4_O_7_, Δ = 0.64 ppm), we were able to identify and characterize seven structural variants by their MS signals appearing, approximately half of which showed the characteristic isotope pattern for chlorine containing compounds such as sandarazol C (583.3256 Da [M+H]^+^; C_29_H_48_ClN_4_O_6_, Δ = 0.07 ppm). ^28^ The sandarazols A, B, C and F were isolated from vanillate induced large scale cultures of MSr10575::pBeloBac Sa001 using a sequence of different techniques such as liquid/liquid extraction, CPC chromatography and HPLC under N_2_ atmosphere (see SI, Figure 2). It is worth noting that not even trace amounts of the sandarazols can be detected in the MSr10575 wild type extracts indicating the *szo* cluster to be fully cryptic in the wild type strain. The structure formulae of sandarazol E and G were assigned by comparison of their MS^2^ spectra to the other derivatives (see SI). As the compounds are sensitive to oxygen and/or strongly acidic or basic pH values, compound isolation was performed in ammonium formate buffered eluents and under constant N_2_ stream wherever possible (see Figure 2). ^1^H, ^13^C and HSQC-dept spectra of sandarazol A reveal two α-protons based on their characteristic chemical shift at *δ*^1^H = 4.78 and 4.45 ppm (see Figure 3). COSY and HMBC correlations of the first methine group at *δ*^1^H = 4.78 ppm to one methylene group, two methyl groups with a downfield *δ*^13^C chemical shift of 45.2 ppm and a quaternary carbon with a characteristic amide shift, show that this α-proton is part of (2R)-2-Amino-3-(N,N-dimethylamino)-propionic acid (D-DimeDap). The second α-proton at *δ*^1^H = 4.45 ppm exhibits COSY and HMBC correlations to one methine group, two methyl groups and another amide function, wherefore the second amino acid of the molecule was determined as valine. HMBC correlations of the valine α-proton to the DimeDap carboxylic acid function suggests their connection via the valine N-terminus. Further HMBC correlations of the DimeDap α-proton to another amide function, reveal N-terminal elongation by the polyketide part of the molecule. Characteristic chemical shifts of two protons at *δ*^1^H = 3.47 ppm and 3.46 ppm with correlations to two aliphatic double bond protons at one side and one epoxide on the other side, suggest sandarazol A to contain an epoxyketone, as well as unsaturation in β-γ position relative to the amide bond. Typical *J*_H-H_ coupling values of 15.4 Hz for the aliphatic double bonds indicate *E* configuration. ^29^ A bis-methylated terminal double bond is found at the ketone side of the epoxyketone using the 1D and 2D NMR spectra. The C-terminal end of valine show correlations to another methine group with characteristic chemical shifts close to an α-proton shift, suggesting further elongation of the molecule by another amino acid. This amino acid is determined to be leucine coupled to a propionate unit by the recorded 1D and 2D NMR spectra. The high field chemical shifts of an additional methyl group at *δ*^1^H = 1.60 and *δ*^13^C = 5.7 ppm in this part of the molecule, reveal heterocyclization of the propionate-leucine motif as a five membered ring via the amide N, which also serves as connection between the valine and leucine building blocks (see Figure 2,3).

**Figure 2.**
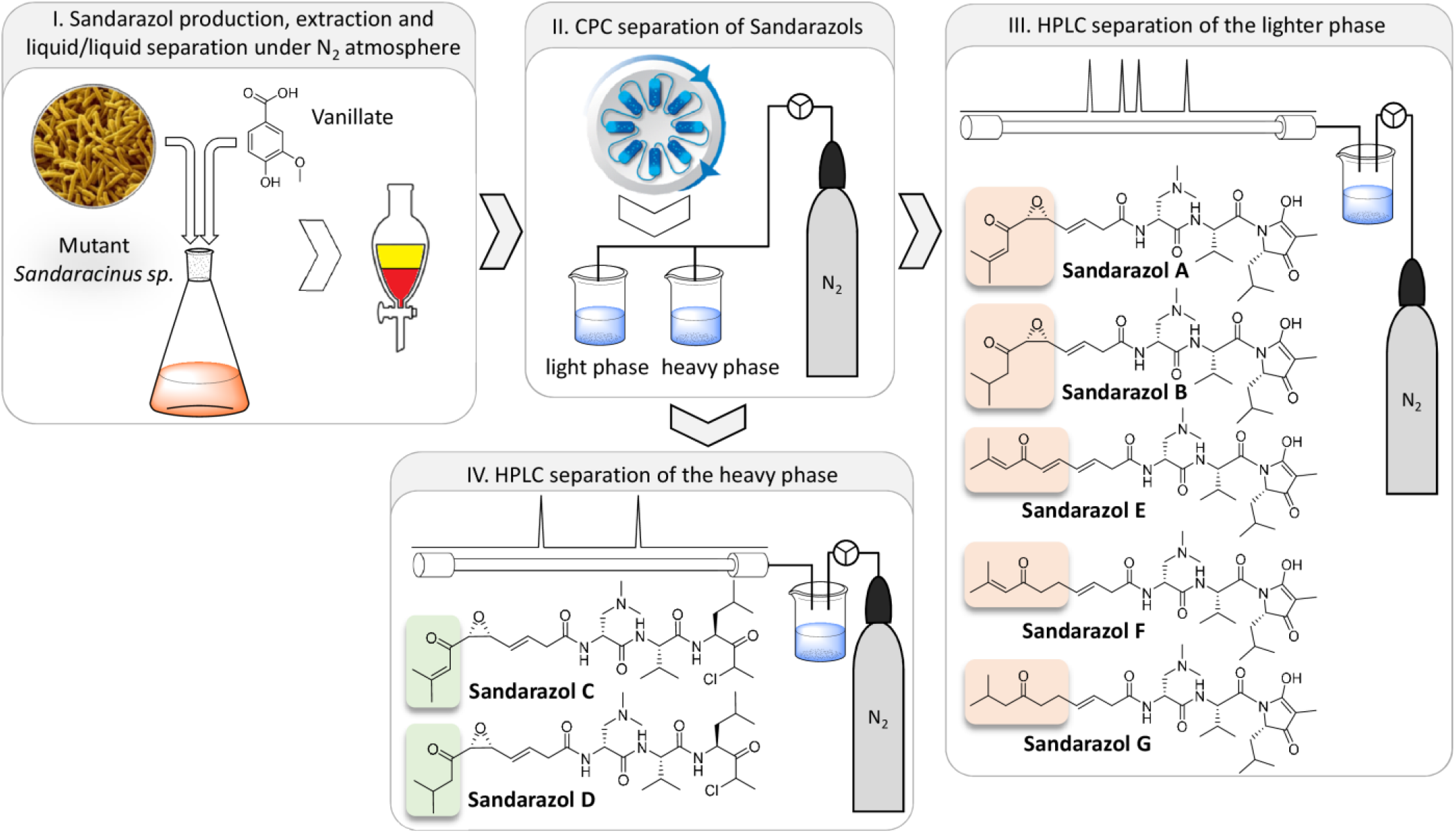
Workflow and isolation scheme for preparation of the sandarazols from *Sandaracinus* sp. MSr10575::pBeloBacSa001 including structure formulae for all structurally elucidated sandarazol derivatives

**Figure 3.**
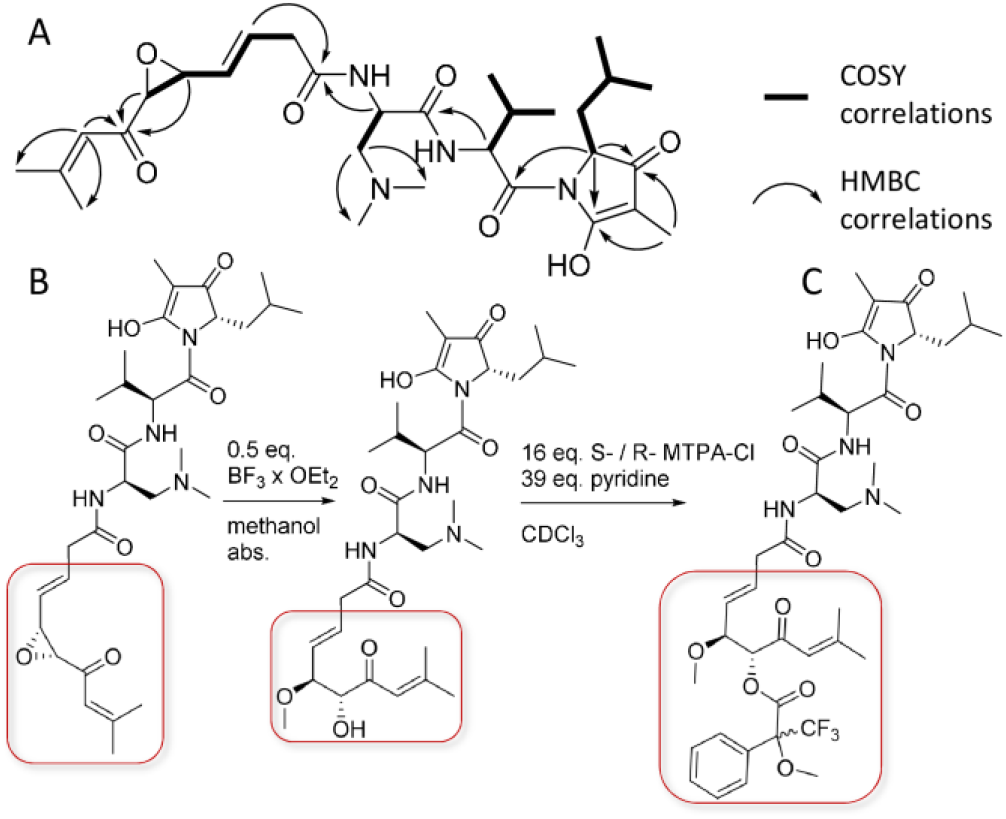
A) NMR correlations important for structure elucidation B) Reaction sequence to open the epoxide and form a methanol adduct C) Mosher’s esterification of sandarazol A used to elucidate the epoxide’s configuration.

In contrast to sandarazol A, HSQC spectra of sandarazol B reveal a missing double bond in the terminal moiety of the polyketide part of the molecule (Figure 2). Sandarazol F differs in the polyketide part of the molecule, as HSQC spectra reveal the two epoxide bearing methines to be replaced by two aliphatic double bond protons. In sandarazol C, however, the polyketide part equates the one of sandarazol A, but 1D and HSQC spectra show a missing elongated leucine derived heterocycle. The methyl group at *δ*^13^C = 6.0 in sandarazol A, is shifted to *δ*^13^C = 20.7 ppm in sandarazol C. In line with the *hr*MS spectra of sandarazol C, chlorination of the molecule is observed at the α-position of the elongated leucine, confirmed by the characteristic chemical shifts and COSY as well as HMBC correlations of the surrounding methyl and methine groups.

All sandarazol derivatives contain 5 stereocenters, two of which are connected by the epoxide. The chlorinated derivatives sandarazol C and D show an additional stereocenter at the chlorinated carbon atom, which racemizes fast at room temperature (see SI). Its original stereo configuration can therefore not be determined as racemization occurs already under fermentation conditions. The stereo centers contained in the L-valine (L-Val), L-leucine (L-Leu) and (2R)-2-amino-3-(N,N-dimethyl amino)-propionic acid (D-DimeDap) building blocks were confirmed by Marfey’s analysis using commercially available standards (see SI). ^30^

The stereocenters in the polyketide part of the molecule are hidden in the orientation of the epoxide. Direct configurative assignment was not possible, as established derivatization methods like the Mosher’s method rely on reaction of functional groups like alcohols. ^31^ To tackle this issue, the epoxide was transformed into an alcohol by Lewis acid catalyzed addition of methanol to the epoxide (Figure 3). The resulting secondary alcohol inherits its stereochemistry from the epoxide due to retention of the epoxide’s stereochemistry in the S_N_2 based ring opening reaction (Figure 3). Subsequent Mosher’s esterification of the secondary alcohol revealed the alcohol to be in R-configuration and thus the epoxide in sandarazols to be R, R-configured. With full structure elucidation, the sandarazols’ absolute configuration as well as the *szo* biosynthetic gene cluster at hand we were able to devise a biosynthetic model for the sandarazols.

### Biosynthesis of the Sandarazols

The sandarazol biosynthesis pathway is based on a type 1 *trans*-AT PKS-NRPS hybrid BGC spanning 44.5 kbp and 15 open reading frames (ORFs) (*szoA* to *szoO*). The megasynthase genes are encoded on the ORFs *szoE* to *szoH*. The in*trans* acting AT and the very unusual in-*trans* acting DH domains are encoded on *szoC* and *szoD*. Tailoring enzymes such as the epoxidase (SzoP), the halogenase (SzoI), the β-branching cassette (SzoK to SzoO) and the biosynthetic machinery to supply the (2L)-2,3-Diamino-propionic acid precursor (SzoB and SzoJ) are encoded on the remaining tailoring enzymes in the BGC (see Figure 4). Sandarazol biosynthesis starts with a *trans*-AT PKS starter module on SzoE that loads an acetate unit onto the first acyl carrier protein (ACP). The peculiarity of this first module is the presence of a β-branching ACP, which acts at the site where the β-branching cassette SzoK to SzoO adds a β-methyl branch to the first PKS extension (Figure 4 B). ^32^ In contrast to standard β-branching cassettes like PyxK to PyxO in the pyxipyrrolone BGC consisting of an ACP, a ketosynthase (KS), a hy-droxymethylglutaryl–CoA synthase (HMG) and two enoyl-CoA dehydratases (ECH), the latter ECH domain is replaced with SzoO, a thioesterase (TE). ^33^ This enzyme is able to replace the decarboxylation function of the second ECH as described in the curacin biosynthesis. ^34^ While most branched chain end moieties in polyketide synthase are biosynthesized by incorporation of an isovaleryl- or an isoamyl-starter unit as in fulvuthiacene biosynthesis for example, bongkrekic acid biosynthesis contains a similar branched chain starter unit biosynthesis based on a β-branching cassette. ^35,36^ As the *szo* cluster neither encodes enoyl reductase (ER) domains nor a standalone short chain reductase protein, the reduction step leading to the production of the saturated tail group seen in the sandarazol variants B, D and G or the α,β saturation next to the ketone in sandarazol F and G remains elusive (structures see Figure 2,4).

**Figure 4.**
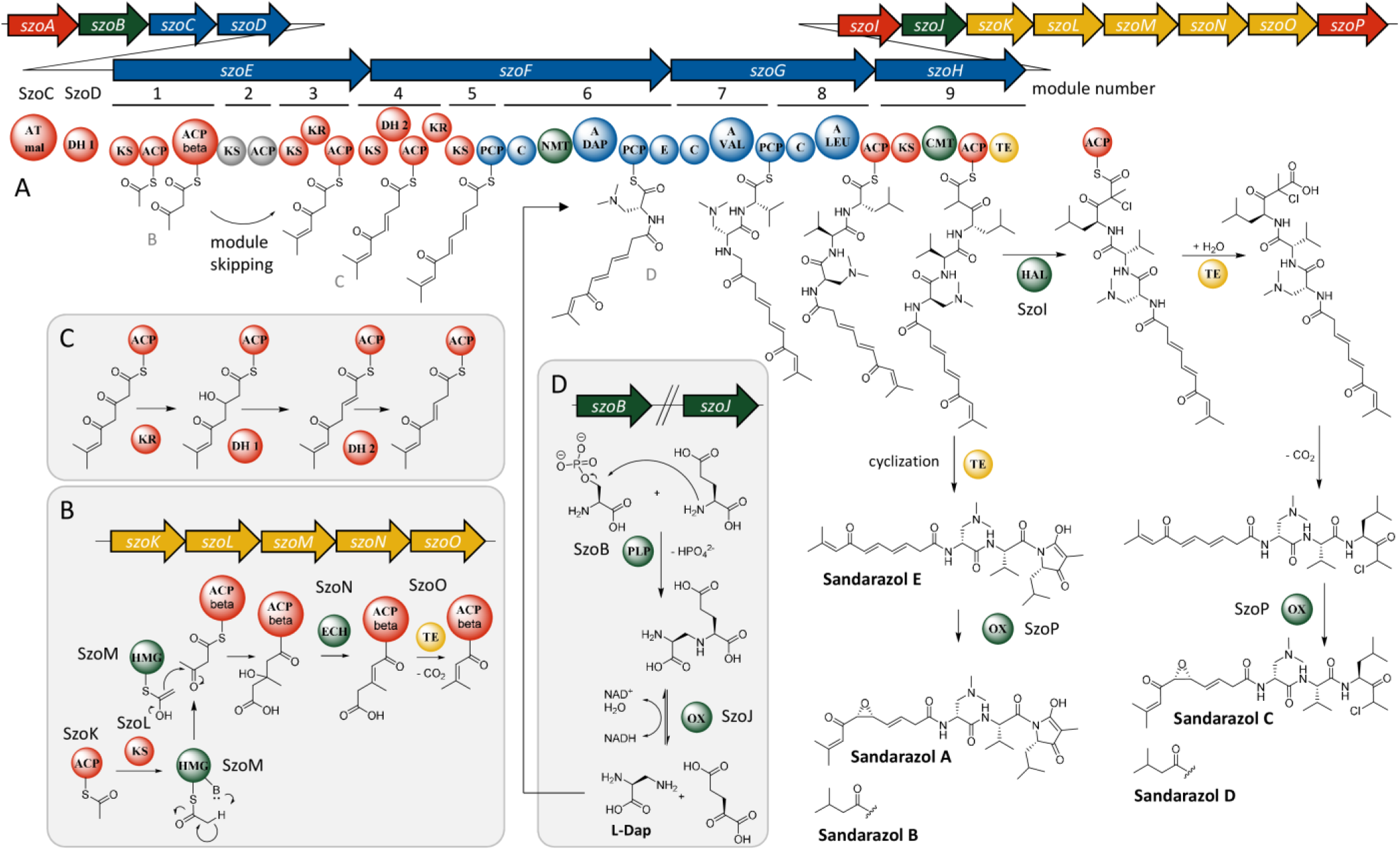
A) Scheme of the sandarazol BGC and sandarazol biosynthesis by its megasynthase including the corresponding tailoring reactions B) β-branching reaction cascade to form the branched chain tail group C) Reaction cascade that leads to the formation of two isomerized double bonds D) Reaction sequence that supplies the amino acid L-Dap to module 6 of the assembly line

Module 2 is most likely skipped as we see no polyketide extension in this module even though both the KS and the ACP in this module seem functional based on sequence alignment analysis. The following modules 3 to 5 incorporate three units of malonyl-CoA by successive decarboxylative claisen condensation to attach 6 carbon atoms to the growing polyketide chain. In this process, we observe consecutive formation of two double bonds through β-ketoreduction by a ketoreductase (KR) and subsequent dehydration most likely performed by the in-*trans-*acting dehydratase SzoD in modules 4 and 5. In-*trans-*acting dehydratases have already been described in *trans*-AT PKS pathways such as the CurE and CorN proteins from curacin and corallopy-ronin respectively. ^37,38^ The KR domain located in module 3 is most likely responsible for β-ketoreduction in module 4, as module 4 does not contain a KR domain and no β-ketoreduction is observed in module 3. Both double bonds formed in modules 4 and 5 are isomerized from an α,β-double bond to a β,γ-double bond by the double bond shifting DH_2_ domain encoded on *szoF*. ^*39*^ Similarly to the double bond shifting domains in corallopyronins, the double bond shifting DH_2_ on SzoF contains the first DH domain consensus motif but lacks the second DxxxQ consensus motif, a characteristic that is conserved in double bond shifting DH domains (see SI). ^38,40^ The subsequent NRPS module on *szoF* contains an adenylation (A) domain with a specificity code close to serine specificity according to NRPS predictor 2. This domain is assumed to accept (2S)-2,3-diamino-propionic acid (L-Dap) that is then incorporated into the nascent chain (Figure 4). ^41^ Biosynthesis of L-Dap is presumed in analogy to the described pathway in *Staphylococcus aureus*. ^42^ The proteins carrying out the corresponding biosynthetic steps in MSr10575 are the SbnA homolog SzoB and the SbnB homolog SzoJ. SbnA is a pyridoxal phosphate (PLP) dependent aminotransferase type enzyme that connects the amino group of glutamate to replace the phosphate moiety of an O-Phosphoserine molecule from the bacterium’s primary metabolism (see Figure 4 D). ^43^ SbnB is an NAD dependent oxidase that releases α-ketoglutarate from this intermediate and forms L-Dap. A SAM dependent N-methyl transferase domain in module 6 subsequently transfers two methyl groups to the free amino group of the attached L-Dap building block to form L-DimeDap, which is subsequently epimerized to D-DimeDap by the following epimerase domain (Figure 4). In contrast to most N-methyl transferase domains built into NRPS systems that transfer a methyl group onto the nitrogen atom that will later form the peptide bond, this methylation reaction transfers two methyl groups to the nitrogen atom in 3 position of L-Dap. ^44^ The following two modules incorporate L-Val and L-Leu according to NRPS textbook logic. ^45^ Incorporation of all three amino acids could be proven by feeding of stable isotope labelled precursors (see SI). Module 9 is a PKS module, that incorporates another malonyl-CoA unit by decarboxylative Claisen condensation and adds an α-methyl branch via a SAM-dependent C-methyl transferase domain to the molecule (Figure 4).

We did not find any non-chlorinated, non-cyclized sandarazols in the culture broth and cyclisation of an amide nitrogen with a carboxylic acid function to form the sandarazol A heterocycle is very unlikely to occur spontaneously. We therefore suspect that chlorination of sandarazols by the halogenase SzoJ occurs on the assembly line. This reaction is likely to govern, whether sandarazol is released from the megasynthase as a chlorinated open chain molecule prone to decarboxylation like seen in sandarazol C, or as a non-chlorinated cyclized product like sandarazol A (Figure 4). Halogenation by SzoJ occurs via the accepted FAD-depended halogenation mechanism similar to synthetic α-chlorination of β-keto acids. ^46,47^ This chlorinated ACP-bound intermediate is released from the assembly line by the TE and quickly loses its terminal carboxylic acid moiety by decarboxylation, a reaction occurring spontaneously in α-chlorinated β-keto acids. If chlorination does not occur, the molecule is subsequently cleaved off the assembly line by a TE domain with a much slower rate, thus cyclizing the product to form a 5 membered ring system as seen for example in sandarazol A and B (see Figure 4). To finalize sandarazol biosynthesis, epoxidation needs to occur next to the ketone moiety in the polyketide part of the molecule. Catalysis of such a reaction is likely performed by FAD-dependent monooxygenase SzoP that putatively performs a reaction similar to the one performed by the FAD-dependent epoxidizing styrene monooxygenase. ^48^ The reductive part of this two component epoxidation system is the reductase protein SzoA that recycles the monooxygenase for the next reaction as it was shown for the styrene monooxygenase. ^49^ We assume that sandarazols are finally exported into the surrounding medium by an ABC exporter of broad range specificity encoded somewhere else in the MSr10575 genome, as no specific exporter system is encoded within or in close proximity to the sandarazol BGC or elsewhere the pSa001 plasmid.

### Biological activity of the Sandarazols

Due to the low stability of the sandarazols, antimicrobial and anti-proliferative activities were only determined for sandarazol A and C as representatives for one chlorinated and one cyclized member of the sandarazol compound family respectively. The non-chlorinated sandarazol A displays only very limited biological activity, while the chlorinated sandarazol C shows prominent cytotoxicity as well as antibiotic activity against *Mycobacterium smegmatis* and a variety of other indicator bacteria for gram-positive pathogens (see Table 1).

**Table 1.** Antimicrobial and cytotoxic activities of sandarazol A and C as minimum inhibitory concentrations (MIC) and inhibitory concentrations at 50% inhibition (IC_50_)

| Microbial strain | MIC sandarazol A | MIC sandarazol C |
| --- | --- | --- |
| *C. albicans* | >110 μM | 110 μM |
| *P. anomala* | >110 μM | 110 μM |
| *C. freundii* | >110 μM | >110 μM |
| *A. baumannii* | >110 μM | >110 μM |
| *S. aureus* | 110 μM | 55 μM |
| *B. subtilis* | >110 μM | 55 μM |
| *E. coli* | >110 μM | >110 μM |
| *P. aeruginosa* | >110 μM | 55 μM |
| *M. smegmatis* | >110 μM | 14 μM |
| Cell line | IC<sub>50</sub> sandarazol A | IC<sub>50</sub> sandarazol C |
| Human colon cancer HCT116 | >64 μM | 0.5 μM |

We observe that the biological activity of the sandarazols mainly stems from the chlorinated sandarazols such as sandarazol C, which shows both the best antimicrobial and anti-proliferative activity. Its high cytotoxicity of 0.5 µM against HCT116 cells might make sandarazol C a promising candidate for further medicinal chemistry approaches. Such approaches directed towards improvements of the sandarazols’ stability against oxygen would probably increase the observable IC_50_ values, as the values shown here cannot be determined under anaerobic conditions that stabilize the sandarazols. The antimicrobial and anti-proliferative activities of sandarazol C indicate, that those secondary metabolites could very well be encoded on the respective plasmid Sa001 to enhance the bacterium’s capability to defend itself against competitors like other bacteria or eukaryotic micro-organisms, while being able to transfer this capability to other bacteria by conjugation.

### Conclusion and Outlook

In this work, we identified pSa001 as the second myxobacterial autonomously replicating plasmid known to date. It was found in the newly isolated *Sandaracinus* sp. MSr10575, the second myxobacterial isolate belonging to the *Sandaracinus* clade. The 209.7 kbp pSa001 plasmid itself encodes the *szo* biosynthetic gene cluster as well as five transposase- and one integrase-type ORFs. Therefore, the plasmid’s content including the *szo* BGC could be transferred between bacterial species, not only by conjugative transfer of the plasmid as an autonomously replicating unit, but also by transposition or integration into the acceptor strain’s genome. We chose to investigate and characterize the pSa001 borne BGC that turned out to be cryptic under laboratory conditions. Previous examples of such plasmid borne secondary metabolite pathways have shown to be responsible for esmeraldin and mycolactone production, both of which show significant biological activity. ^5,6^ The biosynthetic origin of these two compounds led to the theory that the BGC encoded on pSa001 also encodes the biosynthetic machinery for toxin production. After activation of the BGC located on the plasmid pSa001 by promoter exchange, we observed the production of the sandarazols, a series of type1-*trans-*AT PKS-NRPS hybrid secondary metabolites. The sandarazols show promising anti-bacterial and anti-proliferative activities. Following in-detail investigation of the *szo* BGC, we developed a concise biosynthetic model, explaining the sandarazol biosynthesis on its megasynthase protein. Furthermore, we shed light on a variety of uncommon biosynthetic steps such as the first description of incorporation of D-DimeDap into natural products, polyketide β-branching, isomerization of two consecutive double bonds by a single shifting DH-domain and α-chlorination governed release of the sandarazols from the assembly line, all of which warrant further investigations of biosynthetic details. The discovery of the sandarazols does not only highlights the capability of myxobacteria to biosynthesize diverse and biologically active secondary metabolites, it also emphasizes the promise of activating biosynthetic gene clusters by inducible heterologous promoters. Vanillate-inducible promoter systems have proven once more to be able to unlock a larger fraction of the cryptic secondary metabolome of myxobacteria ion promoter exchange experiments. ^9^ The combination of the sandarazols’ biological activity and their intriguing structure and biosynthesis with the biological activity of natural products encoded on other bacterial plasmids highlight plasmid encoded BGCs as valuable targets in the mining of BGCs encoding for novel biologically active natural products in the future. As for myxobacteria, encoding toxin BGCs on plasmids that are transferrable between themselves may provide myxobacteria a competitive edge in their ecological niche. This transfer of toxin production machinery would make particular sense, as myxobacteria do not live individually but rather as part of a swarm that shows wolf pack like predatory behavior. ^50^ So distribution of the genetic means for toxin production among peers should be advantageous for these microorganisms in their collective predation and survival strategies.

## Supporting information

NMR data

pSa001 plasmid sequence

## Author Contributions

All authors have given approval to the final version of the manuscript.

## Acknowledgement

The authors want to thank S. Schmidt for antimicrobial profiling of the sandarazols and A. Amann for the cytotoxicity assays. Furthermore, the authors want to thank N. Zaburannyi for assembly of the MSr10575 genome.

