## Supporting information for "The sandarazols are cryptic and structurally unique plasmid encoded toxins from a rare myxobacterium"

**9 REFERENCES ..........** ERROR! BOOKMARK NOT DEFINED.

### 1 Myxobacterial growth conditions

#### 1.1 Myxobacterial culture media

*Table S 1. Recipe for 2SWT medium*

| 2SWT – Medium | | | |
|---|---|---|---|
| Amount | Ingredient | Concentration | Supplier |
| 3 g/L | Tryptone | - | BD |
| 1 g/L | Soytone | - | BD |
| 3.5 g/L | Soluble Starch | - | Roth |
| 4 g/L | Maltose Monohydrate | - | |
| 2 g/L | Glucose | - | Roth |
| 10 g/L | Starch (soluble) | - | Roth |
| 0.5 g/L | $CaCl_2$ | - | Sigma Aldrich |
| 1 g/L | $MgSO_4$ x $7H_2O$ | - | Grüssing |
| 10 ml/L | TRIS x HCl ph8 | 1M | Sigma Aldrich |
| 100 µL/L | Sterile Vit. B12 solution (added after autoclaving) | 1 mg/ml | Roth |
| 200 µL/L | Sterile FeEDTA solution (added after autoclaving) | 8 mg/ml | Sigma Aldrich |
| Dissolved in milli-Q. Water, pH adjusted to 7.2 with 1N KOH | | | |

*Table S 2. Recipe for S15 medium*

| S15 – Medium | | | |
|---|---|---|---|
| Amount | Ingredient | Concentration | Supplier |
| 3 g/L | Tryptone | - | BD |
| 1 g/L | Soytone | - | BD |
| 3.5 g/L | Soluble Starch | - | Roth |
| 4 g/L | Maltose Monohydrate | - | |
| 2 g/L | Glucose | - | Roth |
| 10 g/L | Starch (soluble) | - | Roth |
| 0.5 g/L | $CaCl_2$ | - | Sigma Aldrich |
| 1 g/L | $MgSO_4$ x $7H_2O$ | - | Grüssing |
| 10 ml/L | TRIS x HCl ph8 | 1M | Sigma Aldrich |
| 100 µL/L | Sterile Vit. B12 solution (added after autoclaving) | 1 mg/ml | Roth |
| 200 µL/L | Sterile FeEDTA solution (added after autoclaving) | 8 mg/ml | Sigma Aldrich |
| Dissolved in milli-Q. Water, pH adjusted to 7.2 with 1N KOH | | | |

The myxobacterial strain MSr10575 was kept in agar culture both for storage over short amounts of time and for cloning. The agar media used are S15 agar and S15 soft agar, which is prepared by adding 14 g/L agarose and 8 g/L agarose (BD) to S15 medium preparations before autoclaving.

*Table S 3. Recipe for 2SWYT medium*

| 2SWYT – Medium | | | |
|---|---|---|---|
| Amount | Ingredient | Concentration | Supplier |
| 3 g/L | Tryptone | - | BD |
| 1 g/L | Soytone | - | BD |
| 3.5 g/L | Soluble Starch | - | Roth |
| 4 g/L | Maltose Monohydrate | - | |
| 10 g/L | Baker's yeast (alive) | - | |
| 2 g/L | Glucose | - | Roth |
| 10 g/L | Starch (soluble) | - | Roth |
| 0.5 g/L | $CaCl_2$ | - | Sigma Aldrich |
| 1 g/L | $MgSO_4$ x $7H_2O$ | - | Grüssing |
| 10 ml/L | TRIS x HCl ph8 | 1M | Sigma Aldrich |
| 100 µL/L | Sterile Vit. B12 solution (added after autoclaving) | 1 mg/ml | Roth |
| 200 µL/L | Sterile FeEDTA solution (added after autoclaving) | 8 mg/ml | Sigma Aldrich |
| Dissolved in milli-Q Water, pH adjusted to 7.2 with 1N KOH | | | |

#### 1.2 Myxobacterial fermentation conditions for LC-MS analysis

Cultures for UHPLC-*hr*MS analysis are grown in 300 ml shake flasks containing 50 ml of 2SWT medium for *Sandaracinus sp.* MSr10575 inoculated with 1 ml of pre culture. Media for mutant MSr10575 strains were supplemented with 50 mg/L kanamycin (Roth) and 1mM of aqueous sterile filtrated K-vanillate solution if the corresponding strain's vanillate promotor is to be induced. After inoculation the medium is supplemented with 2% of sterile XAD-16 adsorber resin (Sigma Aldrich) suspension in water to bind secondary metabolites in the culture medium and limit sandarazol auto toxicity. Small scale cultures were grown for 10-12 days. After fermentation the culture is pelleted in a 50 ml falcon at 6000 rcf for 10 minutes using an Eppendorf falcon table centrifuge and stored at -20°C until further use.

### 2 Analytical methods used in this work

#### 2.1 Metabolite extraction procedure for analytical scale extractions

The frozen cell pellet is transferred into a 100 ml Erlenmeyer flask and a magnetic stirrer is added. 50 ml of acetone (fluka analytical grade, redistilled in house) are added onto the pellet and the mixture is stirred for 60 min on a magnetic stirrer. The acetone extract is left to settle in order to sediment cell debris and XAD resin for a second extraction step. The supernatant is filtered with a 125 micron folded filter keeping cell pellet and XAD-16 resin in the Erlenmeyer flask for a second extraction step. The residual pellet and XAD-16 resin is extracted again with 30 ml of distilled acetone for 60 min on a magnetic stirrer and filtered through the same folded filter. The combined extracts are transferred into a 100 ml round bottom flask. The acetone is evaporated using a rotary evaporator at 260 mbar and 40 °C water bath temperature. The residual water is evaporated at 20 mbar until the residue in the flask is completely dry. The residue is taken up in 550 µl of methanol (Chromasolv HPLC grade, Sigma Aldrich) and transferred into an 1.5 ml Eppendorf tube. This tube is centrifuged with a Hitachi table centrifuge at 15000 rpm for 2 minutes to remove residual insolubilities such as salts, cell debris and XAD fragments. The residual extract is diluted 1:10 for UHPLC-*hr*MS analysis.

#### 2.2 Standardized UHPLC MS conditions

UPLC-*hr*MS analysis is performed on a Dionex (Germering, Germany) Ultimate 3000 RSLC system using a Waters (Eschborn, Germany) BEH C18 column (50 x 2.1 mm, 1.7 µm) equipped with a Waters VanGuard BEH C18 1.7 µm guard column. Separation of 1 µl sample is achieved by a linear gradient from (A) $H_2O$ + 0.1 % FA to (B) ACN + 0.1 % FA at a flow rate of 600 µL/min and a column temperature of 45 °C. Gradient conditions are as follows: 0 – 0.5 min, 5% B; 0.5 – 18.5 min, 5 – 95% B; 18.5 – 20.5 min, 95% B; 20.5 – 21 min, 95 – 5% B; 21-22.5 min, 5% B. UV spectra are recorded by a DAD in the range from 200 to 600 nm. The LC flow is split to 75 µL/min before entering the Bruker Daltonics maXis 4G *hr*ToF mass spectrometer (Bremen, Germany) equipped with an Apollo II ESI source. Mass spectra are acquired in centroid mode ranging from 150 – 2500 m/z at a 2 Hz full scan rate. Mass spectrometry source parameters are set to 500V as end plate offset; 4000V as capillary voltage; nebulizer gas pressure 1 bar; dry gas flow of 5 l/min and a dry temperature of 200°C. Ion transfer and quadrupole settings are set to funnel RF 350 Vpp.; multipole RF 400 Vpp as transfer settings and ion energy of 5eV as well as a low mass cut of 300 m/z. Collision cell is set to 5.0 eV and pre-pulse storage time is set to 5 µs. Spectra acquisition rate is set to 2Hz. Calibration is done automatically before every LC-MS run by injection of sodium formate and calibration on the respective clusters formed in the ESI source. All MS analyses are acquired in the presence of the lock masses ($C_{12}H_{19}F_{12}N_3O_6P_3$, $C_{18}H_{19}O_6N_3P_3F_2$ and $C_{24}H_{19}F_{36}N_3O_6P_3$) which generate the $[M+H]^+$ ions of 622.0289; 922.0098 and 1221.9906.

#### 2.3 Methodology for statistics based metabolome filtering

In order to detect all metabolites appearing after genetic manipulation of MSr10575 we compare wild type MSr10575 to the sandarazol cluster activation mutant in an unbiased principal component analysis (PCA) based statistical analysis adapted from Panter et al..[1] For this purpose, LC-MS chromatograms of 3 independent cultivations are measured as 2 technical replicates each, giving a total number of 6 LC-*hr*MS chromatograms per strain. To obtain all molecular features in the 6 LC-*hr*MS chromatograms of the bacterial extracts of the induced mutant strains and the 6 LC-*hr*MS chromatograms of the corresponding wild type strain extracts, the T-ReX-3D molecular feature finder implemented in Bruker Metaboscape 4.01 is used. Compound detection parameters intensity threshold is set to 10000, mz threshold to 0.005 Da and minimum compound length to 4 spectra. PCA t-test tables are created with the built in PCA t-test routine and filtered according to 6 appearances in the sandarazol cluster activation mutant extract chromatograms and 0 appearances in the MSr10575 wild type extract chromatograms. The t-test table from the bucketing as well as the processed scheduled precursor list used to acquire SPL-MS/MS data will be supplied upon request.

#### 2.4 Acquisition parameters for acquiring high-resolution tandem MS data

LC and MS conditions for SPL guided MS/MS data acquisitions are kept constant according to section standardized UHPLC-MS conditions. MS/MS data acquisition parameters are set to exclusively fragment scheduled precursor list entries. SPL tolerance parameters for precursor ion selection are set to 0.2 minutes and 0.05 m/z in the SPL MS/MS method. The method picks up to 2 precursors per cycle, applies smart exclusion after 5 spectra and performs CID and MS/MS spectra acquisition time ramping. CID Energy is ramped from 35 eV for 500 m/z to 45 eV for 1000 m/z and 60 eV for 2000 m/z. MS full scan acquisition rate is set to 2Hz and MS/MS spectra acquisition rates are ramped from 1 to 4 Hz for precursor ion intensities of 10 kcts. to 1000 kcts..

#### 2.5 Spectral networking parameters for GNPS clustering

All supporting GNPS clustering data presented here is created based on exported .mzML files from the UHPLC-*hr*$MS^2$ chromatograms using the parameters specified in the experimental section of the main text. The MS/MS chromatograms are exported containing all MS/MS data as an .mzML data file and uploaded to the GNPS server at University of California San Diego via FileZilla FTP upload to ftp:// ccms-ftp01.ucsd.edu and all acquired SPL MS/MS spectra are used for spectral network creation. [2] A molecular network is created using the online workflow at GNPS. The data is filtered by removing all MS/MS peaks within +/- 17 Da of the precursor m/z. MS/MS spectra are window filtered by choosing only the top 6 peaks in the +/- 50 Da window throughout the spectrum. The data is then clustered with a parent mass tolerance of 0.05 Da and a MS/MS fragment ion tolerance of 0.1 Da to create consensus spectra. Further, consensus spectra that contained less than 1 spectra are discarded. A network is then created where edges are filtered to have a cosine score above 0.65 and more than 4 matched peaks. Further edges between two nodes are kept in the network if and only if each of the nodes appeared in each other's respective top 10 most similar nodes. [2] The dataset is downloaded from the server and subsequently visualized using Cytoscape 3.7.2.

#### 2.6 Identification of minor sandarazols and sandarazol fragments by GNPS analysis

In this GNPS based spectral networking approach the largest set of clustered MS/MS spectra represents the sandarazol compound family. As one would expect, this GNPS cluster contains the sandarazols described in this work as well as a number of additional minor derivatives of the sandarazols or sandarazol fragments that we could not isolate or structurally elucidate due to their low production titers. Neutral formulas given here are calculated using the built-in tool in Bruker Compass Data analysis.

*Table S 4. List of all MS signals tied to the sandarazol MS/MS cluster created using the GNPS software*

| measured parent mass $[M+H]^+$ | putative neutral formula | compound name |
|---|---|---|
| 575.341 Da | $C_{30}H_{46}N_4O_7$ | Sandarazol A |
| 577.365 Da | $C_{30}H_{48}N_4O_7$ | Sandarazol B |
| 583.325 Da | $C_{29}H_{47}N_4O_6Cl$ | Sandarazol C |
| 585.343 Da | $C_{29}H_{49}N_4O_6Cl$ | Sandarazol D |
| 559.349 Da | $C_{30}H_{46}N_4O_6$ | Sandarazol E |
| 561.365 Da | $C_{30}H_{48}N_4O_6$ | Sandarazol F |
| 563.380 Da | $C_{30}H_{50}N_4O_6$ | Sandarazol G |
| 589.359 Da | $C_{29}H_{53}N_4O_6Cl$ | N.A. |
| 551.357 Da | $C_{28}H_{46}N_4O_7$ | N.A. |
| 549.365 Da | $C_{28}H_{44}N_4O_7$ | N.A. |
| 579.376 Da | $C_{30}H_{50}N_4O_7$ | N.A. |
| 593.354 Da | $C_{30}H_{48}N_4O_8$ | Sandarazol A hydrolysed (diol) |
| 571.362 Da | $C_{29}H_{51}N_4O_5Cl$ | N.A. |

| | | |
|---|---|---|
| 569.346 Da | $C_{29}H_{49}N_4O_5Cl$ | N.A. |
| 545.370 Da | $C_{30}H_{48}N_4O_5$ | N.A. |
| 543.354 Da | $C_{30}H_{46}N_4O_5$ | N.A. |
| 603.375 Da | $C_{32}H_{50}N_4O_7$ | N.A. |
| 770.448 Da | $C_{44}H_{59}N_5O_7$ | N.A. |
| 768.433 Da | $C_{44}H_{57}N_5O_7$ | N.A. |
| 772.466 Da | $C_{44}H_{61}N_5O_7$ | N.A. |
| 802.395 Da | $C_{44}H_{56}N_5O_7Cl$ | N.A. |
| 804.412 Da | $C_{44}H_{58}N_5O_7Cl$ | N.A. |
| 806.426 Da | $C_{44}H_{60}N_5O_7Cl$ | N.A. |

#### 2.7 MS² spectra analysis to assign the sandarazol structures not elucidated by NMR

A couple of sandarazol derivatives that were not structurally elucidated by NMR were assigned by analyzing their MS/MS spectra. The MS/MS spectra were acquired according to the parameters described in the MS/MS parameter description.

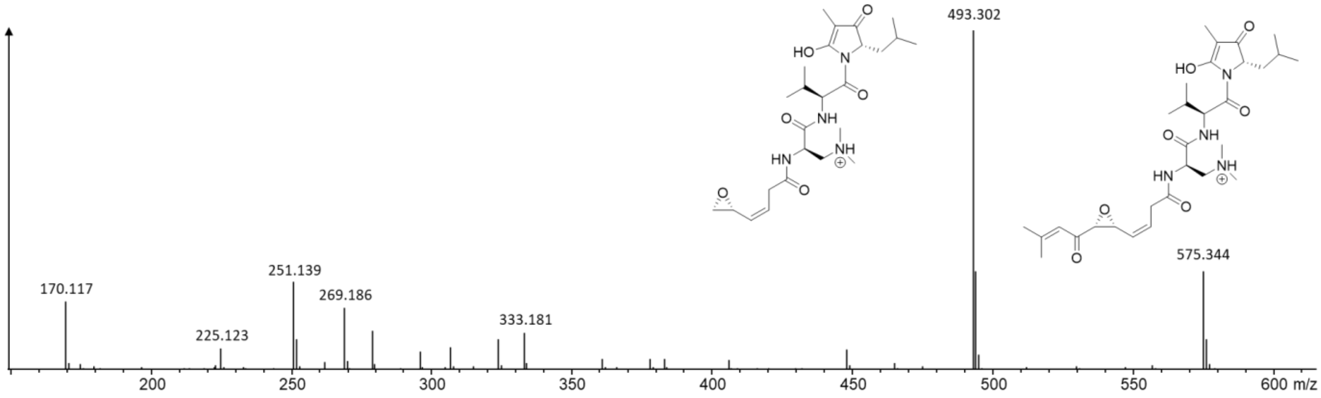

*Figure S 1. Annotated MS² spectrum of sandarazol A*

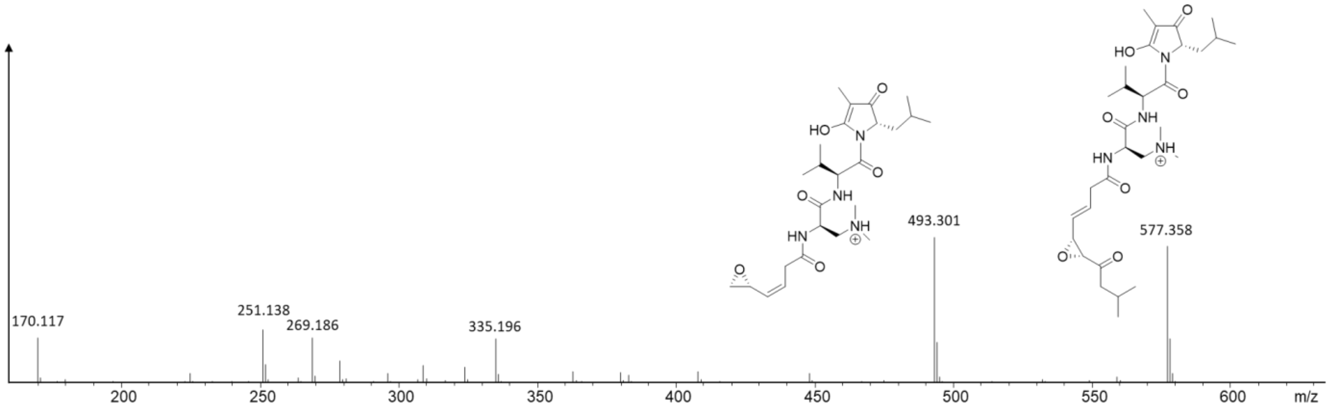

*Figure S 2. Annotated MS² spectrum of sandarazol B*

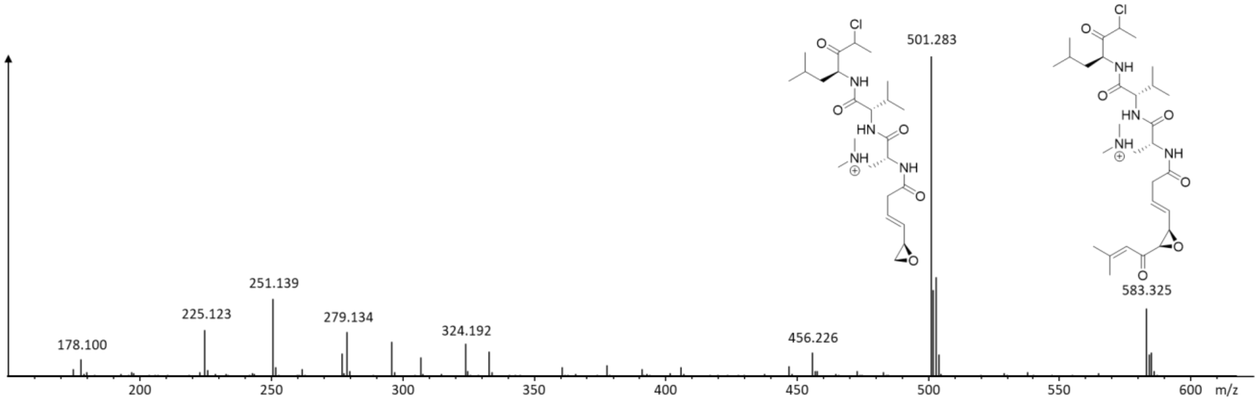

*Figure S 3. Annotated MS² spectrum of sandarazol C*

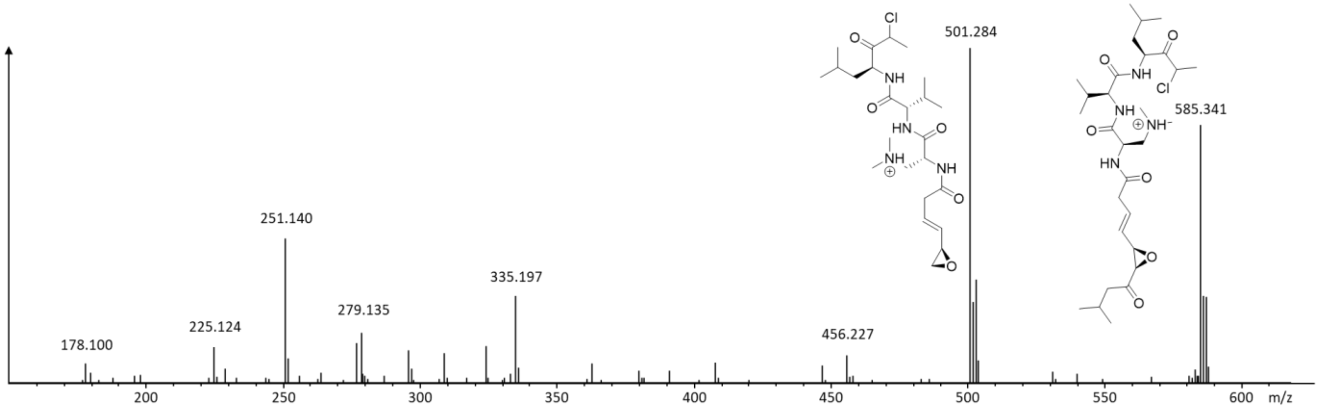

*Figure S 4. Annotated MS² spectrum of sandarazol D*

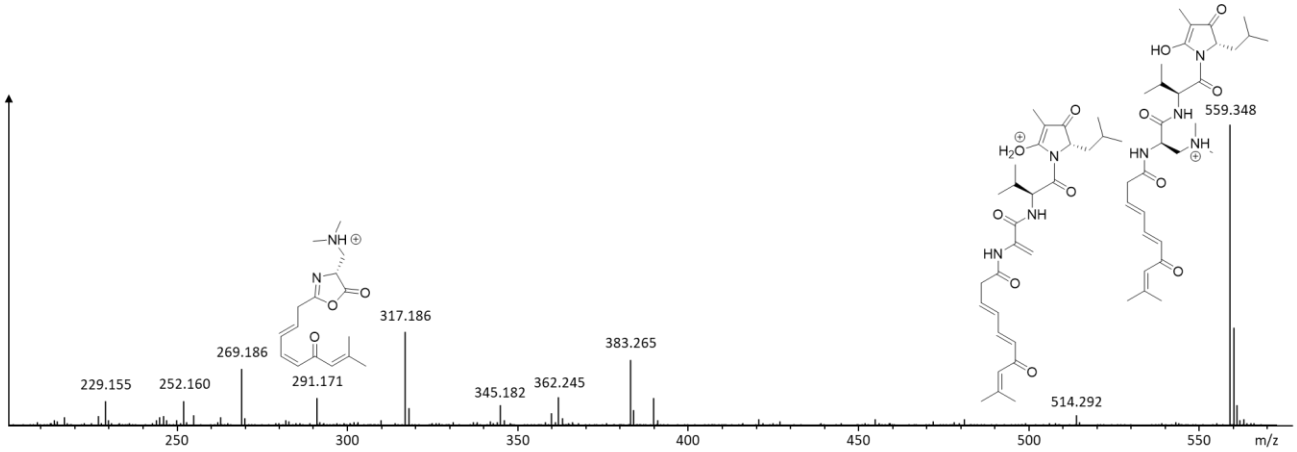

*Figure S 5. Annotated MS² spectrum of sandarazol E*

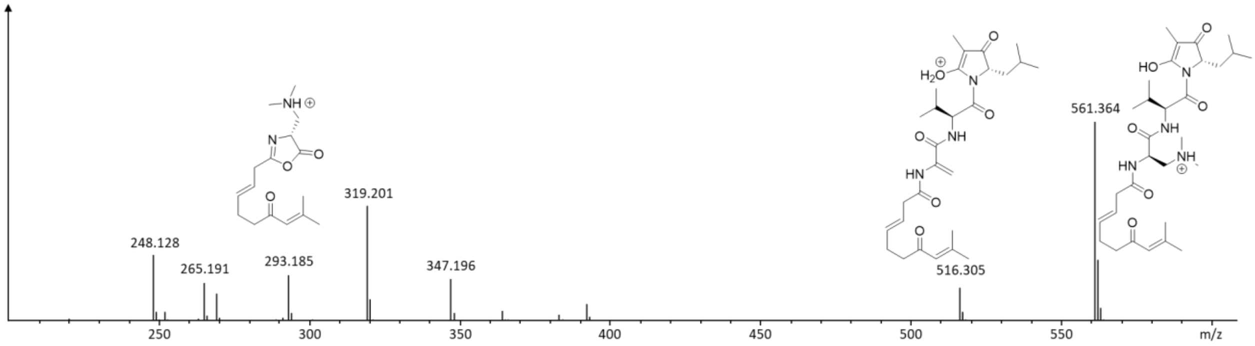

*Figure S 6. Annotated MS² spectrum of sandarazol F*

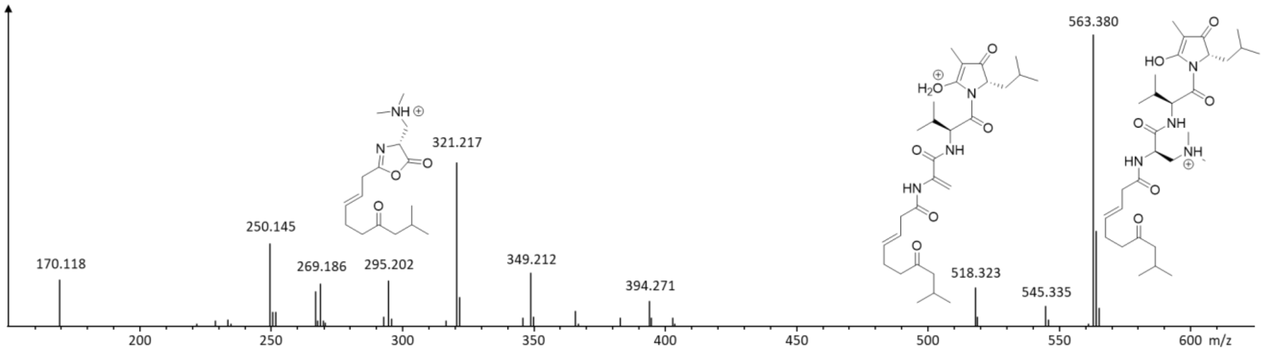

*Figure S 7. Annotated MS² spectrum of sandarazol G*

Sandarazol A, B, C and F were structurally elucidated using multidimensional NMR analyses. The structural formulae of sandarazol D, E and G can be inferred on the basis of the above described NMR spectra. First of all, we can clearly see in the MS² spectra of sandarazol A, B and C, if the structure contains an epoxyketone, as the bond between the epoxide and the ketone is preferentially fragmented. This means that the difference between sandarazol A and B in the structural formulae is the same difference as the difference between sandarazol C and D as both sandarazols show an identical MS² fragment with a terminal epoxide. In sandarazols E, F and G that are all lacking one oxygen atom, we can clearly see that this terminal epoxide fragment does no longer exist. It is therefore safe to assume that the missing oxygen atom is indeed the epoxide oxygen atom in the sandarazols' epoxyketone moiety. In the three MS2 spectra we can see a fragment that is either 291.171, 293.185 or 295.202 Da representing the PKS part of the molecule that still contains the one double bond present in all sandarazols. As we known the structural formula of sandarazol F that has only one additional double bond (for which there would be 2 regioisomers), we can assume that sandarazol E has two additional double bonds and is therefore the direct precursor of sandarazol A, while sandarazol G has no additional double bonds and has therefore only one possible structural formula. As all sandarazols are produced by the same assembly line that is stereospecific, we assume the stereo centers in all sandarazol derivatives to have the same configuration. It is worth mentioning that the stereocenter at the position of the chlorine in sandarazol C and D is rapidly racemizing and we are therefore unable to determine its configuration.

#### 2.8 Stable isotope labelling experiments

In order to take a deeper look at the sandarazol biosynthesis we decided to feed L-Serine to an induced analytical culture of MSr10575 :: pBeloBacSa001 to see whether we observe incorporation into the sandarazols. This would equally mean that the uncommon amino acid (2R)-2-Amino-3-(N,N-dimethyl amino)-propanoic acid (D-DimeDap) provides from serine.

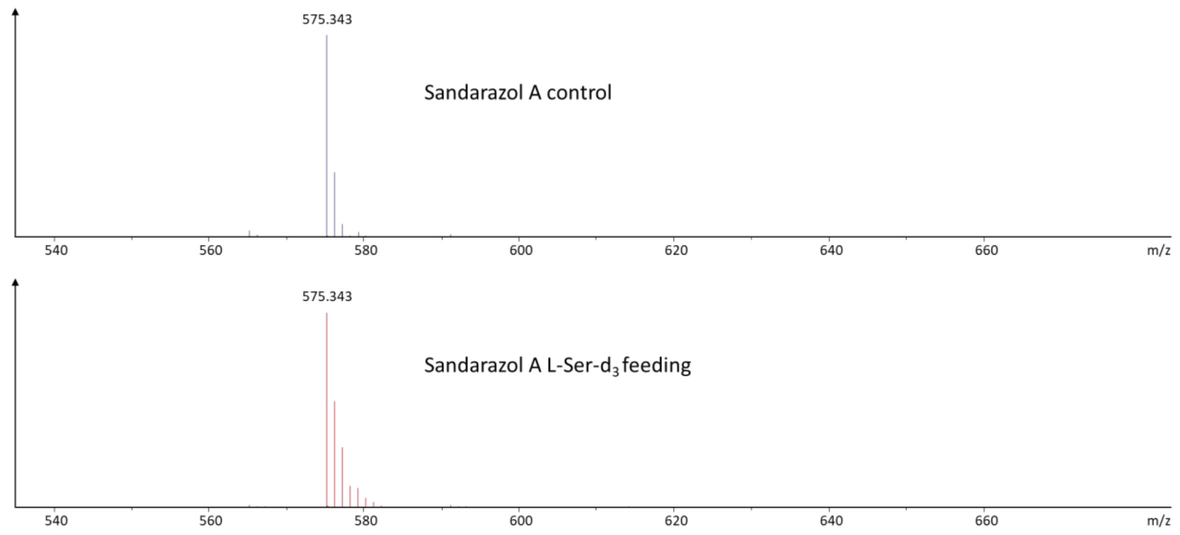

*Figure S 8. Influence of feeding of L-serine-$d_3$ on the isotope pattern of sandarazol A*

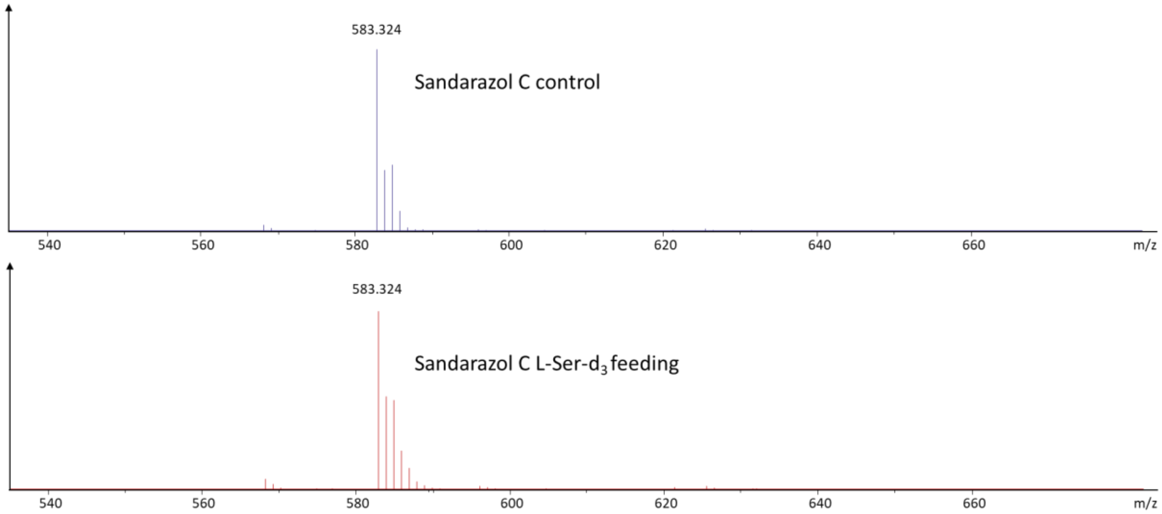

*Figure S 9. Influence of feeding of L-serine-d$_3$ on the isotope pattern of sandarazol C*

We see a clear shift of sandarazols towards higher masses both in sandarazol A and sandarazol C. This means that there is additional evidence that L-DimeDap is indeed synthesized from L-serine as proposed in the sandarazol biosynthesis.

Furthermore, we also fed stable isotope labelled L-Valine and L-Leucine in order to prove incorporation of these amino acids by the sandarazol assembly line. To check the integration of L-Valine, L-Valine-d$_8$ was fed to an induced culture of MSr10575 :: pBeloBacSa001.

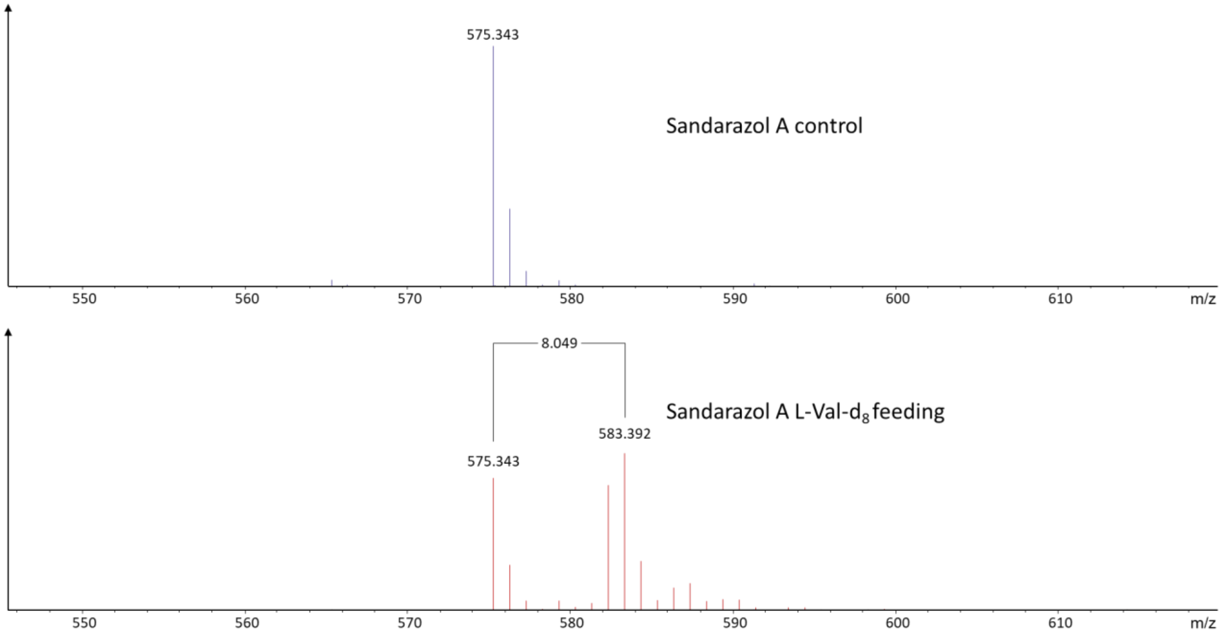

*Figure S 10. Influence of feeding of L-valine-d$_8$ on the isotope pattern of sandarazol A*

We can see a clear +8 da shift upon feeding of L-Val-$d_8$, which is in line with incorporation of Valine into the sandarazols. The small amount of +7 shift is caused by D-H exchange at the alpha position of the Valine, which is often observed in feeding experiments. To check the integration of L-Leucine, L-Leucine-$d_3$ was fed to an induced culture of MSr10575 :: pBeloBacSa001.

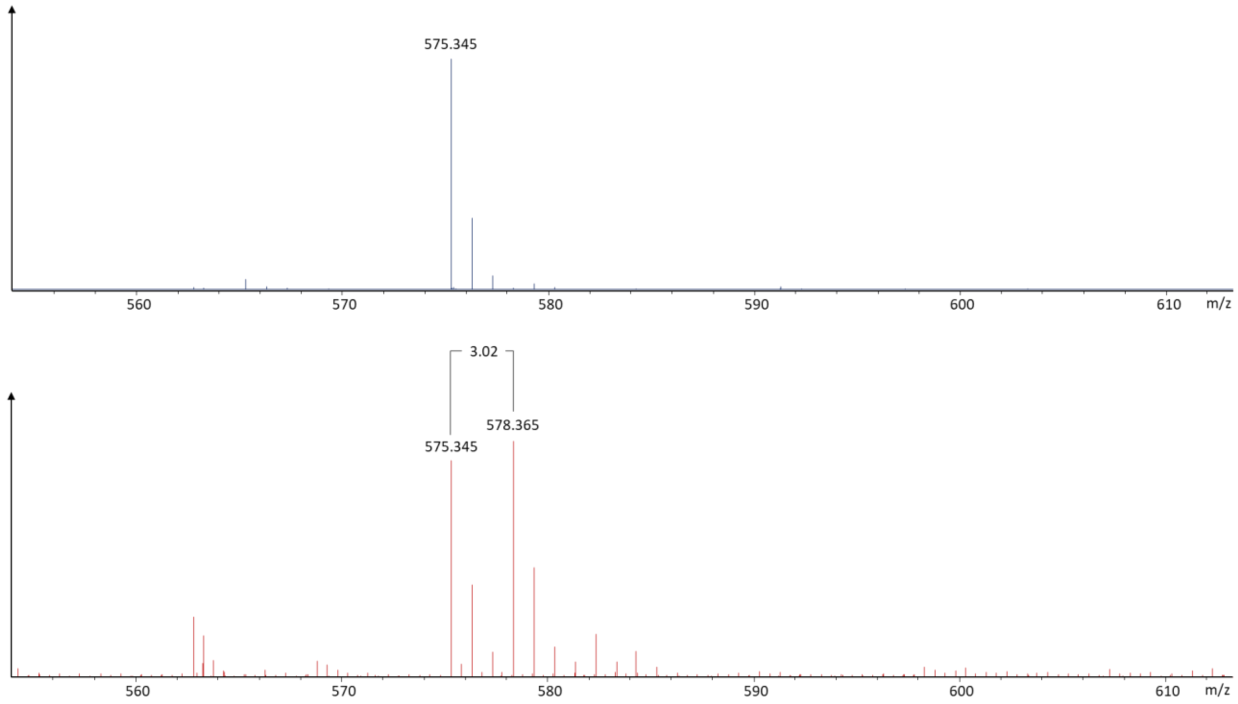

*Figure S 11. Influence of feeding of L-leucine-$d_3$ on the isotope pattern of sandarazol A*

We observe a mass shift of +3 Daltons upon feeding of L-Leucine-$d_3$ which indicates incorporation of the labelled leucine into the sandarazols during biosynthesis.

### 3 Genome sequencing of *Sandaracinus sp.* MSr10575

#### 3.1 Preparation of genomic DNA for PacBio sequencing of MSr10575

To isolate total DNA for sequencing purposes such as PacBio sequencing, phenol-chloroform gDNA extraction is used.

1) Spin down 50ml of fresh myxobacterial culture 6000 rcf 10 min
2) Discard the supernatant
3) Wash the cells once with SET Buffer, centrifuge at 6000 rcf 10 min
4) Resuspend cell pellet in 5ml SET Buffer
5) Add 100 µL of lysozyme (50 mg/ml in $ddH_2O$) stock solution as well as 50µL RNAse A (10 mg/ml in $ddH_2O$) stock solution
6) Add 300 µL Proteinase K solution (10 mg/ml 50 mM Tris 1mM $CaCl_2$) invert several times and add 600µL 10% SDS solution
7) Incubate at 55°C for 2 h, invert every 15 min
8) Add even Volume (6ml) of Phenol/Chloroform/Isoamylalcohol (25:24:1) and swing the tube for 60 min at 5 rpm
9) Centrifuge the mixture at 6000 rcf for 5 min at room temperature
10) Transfer the upper phase into a new tube using a cut end 1ml tip
11) Add even Volume (6ml) of Phenol/Chloroform/Isoamylalcohol (25:24:1) and swing the tube for 60 min at 5 rpm
12) Centrifuge the mixture at 6000 rcf for 5 min at room temperature
13) Transfer the upper phase into a new tube using a cut end 1ml tip
14) Add even Volume (6ml) of Chloroform/Isoamylalcohol (24:1) and swing the tube for 60 min at 5 rpm
15) Centrifuge the mixture at 6000 rcf for 5 min at room temperature
16) Transfer the upper phase into a new tube using a cut end 1ml tip
17) Add 1/10 of the total volume of 3M NaOAc solution pH 5.5 and mix well by inverting several times
18) Add 2.5 Volumes of ice cold ethanol (100% technical purity, -20°C) and invert the tube several times, DNA precipitation should be visible as a cotton like fog in the tube
19) Spool the DNA on a sealed Pasteur pipette

20) Rinse the DNA with 70% Ethanol (cold , -20°C)
21) Air dry the DNA for at least 15 minutes (Dry DNA will become completely translucid)
22) Resuspend dried DNA in 0.5 ml of $ddH_2O$ and keep the Eppendorf tube at room temperature for 24 Hours

#### 3.2 Results of PacBio sequencing of MSr10575 wild type

The wild type strain *Sandaracinus sp.* MSr10575 was sequenced using a PacBio RS II device at the DSMZ using a single SMRT cell. The raw sequence reads were assembled in the SMRT portal software as recommended by the manufacturer. The MSr10575 genome sequence consists of a closed circular bacterial chromosome spanning 10.754 Mbp and a closed circular extrachromosomal mega plasmid spanning 209.732 kbp. Sequence coverage across the circular MSr10575 bacterial chromosome is approximately 65 fold. Coverage of the bacterial mega plasmid containing the *szo* biosynthetic gene cluster is approximately 130 fold indicating a plasmid copy number per cell of about 2 as bases on the mega plasmid are about twice as likely to be sequenced than bases on the genome. The presence of this extrachromosomal mega plasmid can be easily seen even before assembly by the distribution of sequence coverage depth. Assembly of the genomic DNA confirms this finding as it produces two closed unitigs with the above mentioned characteristics.

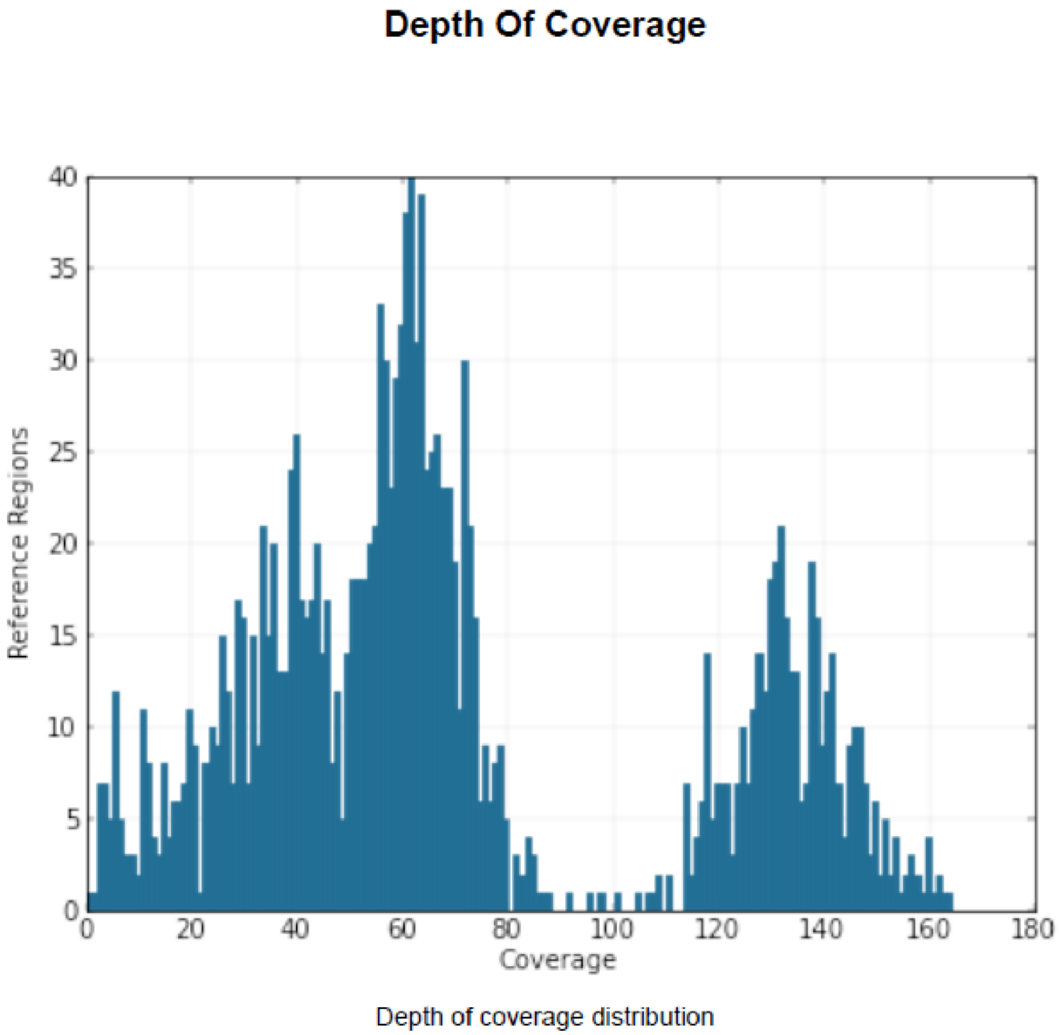

*Figure S 12. Sequence coverage depth distribution per base revealing the presence of an extrachromosomal element with a copy number of approximately 2*

#### 3.3 Codon usage analysis for the MSr10575 chromosome, the pSa001 plasmid and the szo gene cluster

To compare the DNA of MSr10575s Chromosome to its native plasmid pSa001 as well as the corresponding *szo* BGC we performed in depth codon usage analysis.

*Table S 5. Codon usage analysis for the MSr10575 chromosomal DNA*

| MSr10575 chromosomal DNA | | |
|---|---|---|
| Total GC | | 71% |
| Amino Acid (AA) | Freq | % |
| A: | 446,706 | 13.60% |
| C: | 43,918 | 1.30% |
| D: | 210,780 | 6.40% |
| E: | 216,177 | 6.60% |
| F: | 94,839 | 2.90% |
| G: | 289,370 | 8.80% |
| H: | 71,343 | 2.20% |
| I: | 130,988 | 4.00% |
| K: | 50,475 | 1.50% |
| L: | 319,459 | 9.70% |
| M: | 57,556 | 1.70% |
| N: | 47,978 | 1.50% |
| P: | 190,035 | 5.80% |
| Q: | 77,333 | 2.40% |
| R: | 302,988 | 9.20% |
| S: | 175,636 | 5.30% |
| T: | 175,830 | 5.30% |
| V: | 276,434 | 8.40% |
| W: | 45,569 | 1.40% |
| Y: | 57,756 | 1.80% |
| *: | 8,749 | 0.30% |
| AA_Group | Freq | % |
| Acidic: | 426957 | 13.00% |
| Basic: | 424806 | 12.90% |
| Charged: | 851763 | 25.90% |
| Polar_Uncharged: | 624020 | 19.00% |

| | | |
|---|---|---|
| Hydrophobic: | 1850956 | 56.30% |
| GC-rich: | 1229099 | 37.40% |
| AT-rich: | 382036 | 11.60% |

| Codon | AA | Codon usage per AA | Freq |
|---|---|---|---|
| GCA | A | 3.90% | 17,528 |
| GCC | A | 19.90% | 88,883 |
| GCG | A | 74.10% | 331,147 |
| GCT | A | 2.00% | 9,148 |
| TGC | C | 89.30% | 39,218 |
| TGT | C | 10.70% | 4,700 |
| GAC | D | 73.40% | 154,779 |
| GAT | D | 26.60% | 56,001 |
| GAA | E | 8.40% | 18,206 |
| GAG | E | 91.60% | 197,971 |
| TTC | F | 98.90% | 93,795 |
| TTT | F | 1.10% | 1,044 |
| GGA | G | 8.00% | 23,240 |
| GGC | G | 63.00% | 182,206 |
| GGG | G | 19.60% | 56,606 |
| GGT | G | 9.40% | 27,318 |
| CAC | H | 88.00% | 62,797 |
| CAT | H | 12.00% | 8,546 |
| ATA | I | 0.10% | 151 |
| ATC | I | 98.50% | 129,043 |
| ATT | I | 1.40% | 1,794 |
| AAA | K | 2.60% | 1,294 |
| AAG | K | 97.40% | 49,181 |
| CTA | L | 0.30% | 803 |
| CTC | L | 64.60% | 206,223 |
| CTG | L | 31.70% | 101,292 |
| CTT | L | 1.10% | 3,430 |
| TTA | L | 0.00% | 53 |

| | | | |
|---|---|---|---|
| TTG | L | 2.40% | 7,658 |
| ATG | M | 100% | 57,556 |
| AAC | N | 92.40% | 44,336 |
| AAT | N | 7.60% | 3,642 |
| CCA | P | 1.90% | 3,587 |
| CCC | P | 36.00% | 68,493 |
| CCG | P | 58.30% | 110,801 |
| CCT | P | 3.80% | 7,154 |
| CAA | Q | 4.20% | 3,212 |
| CAG | Q | 95.80% | 74,121 |
| AGA | R | 0.50% | 1,384 |
| AGG | R | 1.40% | 4,327 |
| CGA | R | 6.20% | 18,776 |
| CGC | R | 69.30% | 209,886 |
| CGG | R | 14.40% | 43,521 |
| CGT | R | 8.30% | 25,094 |
| AGC | S | 37.50% | 65,925 |
| AGT | S | 1.20% | 2,074 |
| TCA | S | 0.90% | 1,546 |
| TCC | S | 8.60% | 15,157 |
| TCG | S | 50.60% | 88,949 |
| TCT | S | 1.10% | 1,985 |
| ACA | T | 1.20% | 2,110 |
| ACC | T | 41.50% | 73,025 |
| ACG | T | 55.60% | 97,746 |
| ACT | T | 1.70% | 2,949 |
| GTA | V | 0.40% | 1,155 |
| GTC | V | 46.60% | 128,748 |
| GTG | V | 51.80% | 143,167 |
| GTT | V | 1.20% | 3,364 |
| TGG | W | 100% | 45,569 |
| TAC | Y | 85.10% | 49,169 |

| | | | |
|---|---|---|---|
| TAT | Y | 14.90% | 8,587 |
| TAA | * | 1.40% | 126 |
| TAG | * | 10.60% | 928 |
| TGA | * | 88.00% | 7,695 |

*Table S 6. Codon usage analysis for the MSr10575 borne pSa001 plasmid*

| pSa001 plasmid DNA | | |
|---|---|---|
| Total GC | | 70% |
| Amino Acid (AA) | Freq | % |
| A: | 5,874 | 13.90% |
| C: | 896 | 2.10% |
| D: | 2,814 | 6.70% |
| E: | 2,623 | 6.20% |
| F: | 1,205 | 2.90% |
| G: | 3,729 | 8.80% |
| H: | 982 | 2.30% |
| I: | 1,524 | 3.60% |
| K: | 430 | 1.00% |
| L: | 3,884 | 9.20% |
| M: | 599 | 1.40% |
| N: | 658 | 1.60% |
| P: | 2,564 | 6.10% |
| Q: | 859 | 2.00% |
| R: | 3,955 | 9.40% |
| S: | 2,547 | 6.00% |
| T: | 2,251 | 5.30% |
| V: | 3,367 | 8.00% |
| W: | 588 | 1.40% |
| Y: | 715 | 1.70% |
| *: | 112 | 0.30% |

| AA_Group | Freq | % |
|---|---|---|
| Acidic: | 5437 | 12.90% |
| Basic: | 5367 | 12.70% |
| Charged: | 10804 | 25.60% |
| Polar_Uncharged: | 8514 | 20.20% |
| Hydrophobic: | 23334 | 55.30% |
| GC-rich: | 16122 | 38.20% |
| AT-rich: | 4532 | 10.70% |

| Codon | AA | Codon usage per AA | Freq |
|---|---|---|---|
| GCA | A | 9.00% | 528 |
| GCC | A | 22.10% | 1,297 |
| GCG | A | 62.80% | 3,691 |
| GCT | A | 6.10% | 358 |
| TGC | C | 86.20% | 772 |
| TGT | C | 13.80% | 124 |
| GAC | D | 70.00% | 1,969 |
| GAT | D | 30.00% | 845 |
| GAA | E | 15.90% | 416 |
| GAG | E | 84.10% | 2,207 |
| TTC | F | 92.40% | 1,114 |
| TTT | F | 7.60% | 91 |
| GGA | G | 13.20% | 492 |
| GGC | G | 55.50% | 2,068 |
| GGG | G | 18.30% | 681 |
| GGT | G | 13.10% | 488 |
| CAC | H | 81.80% | 803 |
| CAT | H | 18.20% | 179 |
| ATA | I | 1.60% | 24 |
| ATC | I | 89.80% | 1,369 |
| ATT | I | 8.60% | 131 |

| | | | |
|---|---|---|---|
| AAA | K | 10.50% | 45 |
| AAG | K | 89.50% | 385 |
| CTA | L | 1.30% | 49 |
| CTC | L | 61.60% | 2,393 |
| CTG | L | 27.00% | 1,049 |
| CTT | L | 4.60% | 179 |
| TTA | L | 0.30% | 10 |
| TTG | L | 5.30% | 204 |
| ATG | M | 100% | 599 |
| AAC | N | 82.70% | 544 |
| AAT | N | 17.30% | 114 |
| CCA | P | 4.30% | 110 |
| CCC | P | 30.50% | 782 |
| CCG | P | 56.70% | 1,454 |
| CCT | P | 8.50% | 218 |
| CAA | Q | 13.20% | 113 |
| CAG | Q | 86.80% | 746 |
| AGA | R | 1.10% | 45 |
| AGG | R | 2.60% | 101 |
| CGA | R | 10.00% | 395 |
| CGC | R | 56.70% | 2,242 |
| CGG | R | 17.40% | 689 |
| CGT | R | 12.20% | 483 |
| AGC | S | 30.40% | 775 |
| AGT | S | 4.60% | 116 |
| TCA | S | 2.80% | 71 |
| TCC | S | 15.30% | 390 |
| TCG | S | 43.70% | 1,114 |
| TCT | S | 3.20% | 81 |
| ACA | T | 4.00% | 91 |
| ACC | T | 35.20% | 793 |
| ACG | T | 55.20% | 1,243 |

| | | | |
|---|---|---|---|
| ACT | T | 5.50% | 124 |
| GTA | V | 3.10% | 105 |
| GTC | V | 46.90% | 1,579 |
| GTG | V | 45.10% | 1,517 |
| GTT | V | 4.90% | 166 |
| TGG | W | 100% | 588 |
| TAC | Y | 80.40% | 575 |
| TAT | Y | 19.60% | 140 |
| TAA | * | 3.60% | 4 |
| TAG | * | 8.00% | 9 |
| TGA | * | 88.40% | 99 |

*Table S 7. Codon usage analysis for the* szo *BGC*

| *szo* Cluster on pSa001 | | |
|---|---|---|
| Total GC | | 71% |
| Amino Acid (AA) | Freq | % |
| A: | 2,511 | 14.80% |
| C: | 212 | 1.30% |
| D: | 1,127 | 6.70% |
| E: | 1,061 | 6.30% |
| F: | 473 | 2.80% |
| G: | 1,541 | 9.10% |
| H: | 449 | 2.70% |
| I: | 647 | 3.80% |
| K: | 151 | 0.90% |
| L: | 1,742 | 10.30% |
| M: | 202 | 1.20% |
| N: | 219 | 1.30% |
| P: | 964 | 5.70% |
| Q: | 316 | 1.90% |

| | | |
|---|---|---|
| R: | 1,663 | 9.80% |
| S: | 960 | 5.70% |
| T: | 820 | 4.80% |
| V: | 1,370 | 8.10% |
| W: | 212 | 1.30% |
| Y: | 276 | 1.60% |
| *: | 23 | 0.10% |

| AA_Group | Freq | % |
|---|---|---|
| Acidic: | 2188 | 12.90% |
| Basic: | 2263 | 13.40% |
| Charged: | 4451 | 26.30% |
| Polar_Uncharged: | 3015 | 17.80% |
| Hydrophobic: | 9662 | 57.00% |
| GC-rich: | 6679 | 39.40% |
| AT-rich: | 1766 | 10.40% |

| Codon | AA | Codon usage per AA | Freq |
|---|---|---|---|
| GCA | A | 8.20% | 205 |
| GCC | A | 21.60% | 543 |
| GCG | A | 65.20% | 1,636 |
| GCT | A | 5.10% | 127 |
| TGC | C | 85.80% | 182 |
| TGT | C | 14.20% | 30 |
| GAC | D | 71.40% | 805 |
| GAT | D | 28.60% | 322 |
| GAA | E | 13.50% | 143 |
| GAG | E | 86.50% | 918 |
| TTC | F | 94.90% | 449 |
| TTT | F | 5.10% | 24 |
| GGA | G | 11.90% | 183 |
| GGC | G | 55.90% | 862 |

| | | | |
|---|---|---|---|
| GGG | G | 18.40% | 283 |
| GGT | G | 13.80% | 213 |
| CAC | H | 84.20% | 378 |
| CAT | H | 15.80% | 71 |
| ATA | I | 1.40% | 9 |
| ATC | I | 92.10% | 596 |
| ATT | I | 6.50% | 42 |
| AAA | K | 9.30% | 14 |
| AAG | K | 90.70% | 137 |
| CTA | L | 1.10% | 19 |
| CTC | L | 67.60% | 1,178 |
| CTG | L | 24.50% | 426 |
| CTT | L | 3.30% | 58 |
| TTA | L | 0.20% | 3 |
| TTG | L | 3.30% | 58 |
| ATG | M | 100% | 202 |
| AAC | N | 82.60% | 181 |
| AAT | N | 17.40% | 38 |
| CCA | P | 2.70% | 26 |
| CCC | P | 30.40% | 293 |
| CCG | P | 60.10% | 579 |
| CCT | P | 6.80% | 66 |
| CAA | Q | 9.80% | 31 |
| CAG | Q | 90.20% | 285 |
| AGA | R | 0.70% | 11 |
| AGG | R | 1.40% | 23 |
| CGA | R | 9.40% | 157 |
| CGC | R | 60.60% | 1,008 |
| CGG | R | 15.80% | 262 |
| CGT | R | 12.10% | 202 |
| AGC | S | 28.10% | 270 |
| AGT | S | 2.70% | 26 |

| | | | |
|---|---|---|---|
| TCA | S | 0.90% | 9 |
| TCC | S | 16.00% | 154 |
| TCG | S | 49.40% | 474 |
| TCT | S | 2.80% | 27 |
| ACA | T | 2.30% | 19 |
| ACC | T | 36.30% | 298 |
| ACG | T | 56.70% | 465 |
| ACT | T | 4.60% | 38 |
| GTA | V | 2.30% | 31 |
| GTC | V | 50.60% | 693 |
| GTG | V | 43.50% | 596 |
| GTT | V | 3.60% | 50 |
| TGG | W | 100% | 212 |
| TAC | Y | 80.80% | 223 |
| TAT | Y | 19.20% | 53 |
| TAA | * | 0.00% | 0 |
| TAG | * | 0.00% | 0 |
| TGA | * | 100% | 23 |

#### 3.4 Results of Illumina sequencing of MSr10575 mutants

In order to confirm successful homologous recombination in MSr10575 we prepared MSr10575 :: pBeloBac Sa001 genomic DNA and sent it to GMAK (Braunschweig, Germany) for Illumina sequencing. Results confirm that on the one hand, the extrachromosomal element in MSr10575 remains a plasmid and is not recombined into the MSr10575 bacterial chromosome and on the other hand that the homologous recombination to create pBeloBac Sa001 did not create any errors such as SNP's or short insertion/deletions.

### 4 *In-silico* analysis of the sandarazol biosynthetic gene cluster

The sandarazol biosynthetic gene cluster is a type 1 Trans-AT Polyketide synthase (PKS) non-ribosomal peptide synthethase (NRPS) hybrid biosynthetic gene cluster located on a natural plasmid spanning 209,732 bp in *Sandaracinus sp.* MSr10575 wild type. The biosynthetic gene cluster with the corresponding annotations can be found under GenBank tag Sandaracinus strain MSr10575 pSa001 plasmid under GenBank accession number MW053453. Based on this GenBank file, the BGC will be available at the MiBiG database.

#### 4.1 Overview over the pSa001 plasmid

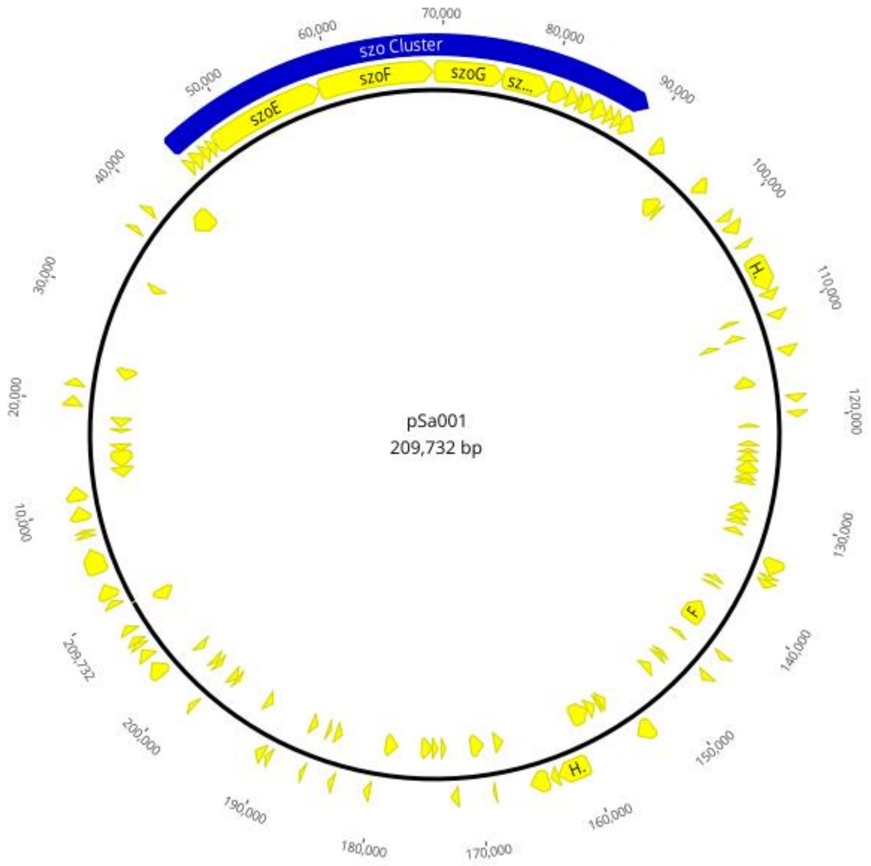

*Figure S 13. Schematic view of the pSa001 plasmid in MSr10575*

*Table S 8. Tabulated view of all detected CDS on the pSa001 plasmid*

| Predicted protein product | CDS start | CDS end | Length | reading direction |
|---|---|---|---|---|
| Hypothetical protein CDS | 424 | 1,092 | 669 | forward |
| Hypothetical protein CDS | 1,239 | 2,678 | 1,440 | forward |
| Hypothetical protein CDS | 3,780 | 6,122 | 2,343 | forward |
| Hypothetical protein CDS | 7,308 | 7,649 | 342 | forward |
| Hypothetical protein CDS | 7,748 | 8,023 | 276 | forward |
| Hypothetical protein CDS | 8,914 | 10,161 | 1,248 | forward |
| Hypothetical protein CDS | 10,784 | 12,055 | 1,272 | forward |
| Hypothetical protein CDS | 12,353 | 13,375 | 1,023 | reverse |
| Serine/threonine protein kinase CDS | 13,372 | 15,174 | 1,803 | reverse |
| Hypothetical protein CDS | 15,171 | 15,899 | 729 | reverse |
| Hypothetical protein CDS | 17,118 | 17,507 | 390 | reverse |
| Hypothetical protein CDS | 17,721 | 18,662 | 942 | reverse |
| Serine/threonine protein kinase CDS | 19,526 | 20,536 | 1,011 | forward |
| Hypothetical protein CDS | 21,401 | 22,156 | 756 | forward |
| Transposase family protein CDS | 22,701 | 23,870 | 1,170 | reverse |
| Hypothetical protein CDS | 32,461 | 33,102 | 642 | reverse |
| Hypothetical protein CDS | 36,823 | 37,218 | 396 | forward |
| FG-GAP repeat-HVR domain containing protein CDS | 38,898 | 39,500 | 603 | forward |
| Hypothetical protein CDS | 40,535 | 42,781 | 2,247 | reverse |
| szoA CDS | 44,462 | 45,190 | 729 | forward |
| szoB CDS | 45,218 | 46,258 | 1,041 | forward |
| szoC CDS | 46,255 | 47,103 | 849 | forward |
| szoD CDS | 47,100 | 47,843 | 744 | forward |
| szoE CDS | 47,836 | 58,599 | 10,764 | forward |
| szoF CDS | 58,596 | 69,254 | 10,659 | forward |
| szoG CDS | 69,290 | 75,655 | 6,366 | forward |

| | | | | |
|---|---|---|---|---|
| szoH CDS | 75,652 | 80,040 | 4,389 | forward |
| szoI CDS | 80,103 | 81,854 | 1,752 | forward |
| szoJ CDS | 81,863 | 82,915 | 1,053 | forward |
| szoK CDS | 82,912 | 83,196 | 285 | forward |
| szoL CDS | 83,193 | 84,515 | 1,323 | forward |
| szoM CDS | 84,512 | 85,747 | 1,236 | forward |
| szoN CDS | 85,744 | 86,487 | 744 | forward |
| szoO CDS | 86,591 | 87,379 | 789 | forward |
| szoP CDS | 87,428 | 88,675 | 1,248 | forward |
| chitinase CDS | 90,944 | 92,113 | 1,170 | forward |
| Hemolysin-type calcium binding region CDS | 93,669 | 95,156 | 1,488 | reverse |
| Hypothetical protein CDS | 95,305 | 95,604 | 300 | reverse |
| Hypothetical protein CDS | 96,300 | 97,517 | 1,218 | forward |
| Hypothetical protein CDS | 100,082 | 100,789 | 708 | forward |
| Oxidoreductase, NmrA-like CDS | 101,215 | 102,189 | 975 | forward |
| Hypothetical protein CDS | 103,311 | 103,757 | 447 | forward |
| Hypothetical protein CDS | 104,927 | 108,178 | 3,252 | forward |
| Hypothetical protein CDS | 108,191 | 109,087 | 897 | forward |
| Hypothetical protein CDS | 109,621 | 109,992 | 372 | reverse |
| transposase CDS | 110,176 | 111,045 | 870 | forward |
| Activator of HSP90 ATPase 1 family protein CDS | 111,170 | 111,658 | 489 | reverse |
| arsR CDS | 111,637 | 112,071 | 435 | reverse |
| Hypothetical protein CDS | 113,516 | 114,496 | 981 | forward |
| Hypothetical protein CDS | 115,666 | 116,799 | 1,134 | reverse |
| Cobyrinic acid ac-diamidesynthase CDS | 118,113 | 118,772 | 660 | forward |
| Hypothetical protein CDS | 119,528 | 120,130 | 603 | forward |
| Hypothetical protein CDS | 120,567 | 121,022 | 456 | reverse |
| Hypothetical protein CDS | 122,528 | 122,926 | 399 | reverse |
| Hypothetical protein CDS | 123,247 | 123,663 | 417 | reverse |
| Hypothetical protein CDS | 123,660 | 124,586 | 927 | reverse |

| | | | | |
|---|---|---|---|---|
| Peptidase, M23/M37 family CDS | 124,583 | 125,872 | 1,290 | reverse |
| Hypothetical protein CDS | 125,869 | 126,546 | 678 | reverse |
| Hypothetical protein CDS | 126,566 | 127,024 | 459 | reverse |
| Hypothetical protein CDS | 129,053 | 129,844 | 792 | reverse |
| Hypothetical protein CDS | 129,857 | 130,693 | 837 | reverse |
| Hypothetical protein CDS | 130,745 | 131,302 | 558 | reverse |
| Hypothetical protein CDS | 131,693 | 132,355 | 663 | reverse |
| Hypothetical protein CDS | 133,584 | 135,074 | 1,491 | forward |
| Hypothetical protein CDS | 135,169 | 135,609 | 441 | forward |
| Hypothetical protein CDS | 135,650 | 136,342 | 693 | forward |
| Hypothetical protein CDS | 137,252 | 137,734 | 483 | reverse |
| Hypothetical protein CDS | 138,098 | 138,316 | 219 | reverse |
| FG-GAP repeat/HVR domain protein CDS | 140,368 | 142,821 | 2,454 | reverse |
| Hypothetical protein CDS | 143,491 | 144,018 | 528 | forward |
| Hypothetical protein CDS | 144,309 | 144,659 | 351 | reverse |
| Hypothetical protein CDS | 145,832 | 146,539 | 708 | forward |
| Hypothetical protein CDS | 147,166 | 147,534 | 369 | reverse |
| Hypothetical protein CDS | 147,544 | 147,918 | 375 | reverse |
| Sigma54 specific transcriptional regulator, Fis family CDS | 148,954 | 149,781 | 828 | reverse |
| Integrase CDS | 152,839 | 154,389 | 1,551 | forward |
| transcriptional regulator CDS | 155,087 | 155,770 | 684 | reverse |
| Glyoxalase /bleomycin resistance protein/ dioxygenase CDS | 155,775 | 156,173 | 399 | reverse |
| Hypothetical protein CDS | 156,401 | 157,324 | 924 | reverse |
| Serine/threonine protein kinase CDS | 157,324 | 159,282 | 1,959 | reverse |
| Hypothetical protein CDS | 159,642 | 162,587 | 2,946 | forward |
| Serine/threonine specific protein phosphatase (Putative) CDS | 162,596 | 163,369 | 774 | forward |
| Hypothetical protein CDS | 163,605 | 165,287 | 1,683 | forward |

| | | | | |
|---|---|---|---|---|
| Hypothetical protein CDS | 166,983 | 167,897 | 915 | reverse |
| Hypothetical protein CDS | 168,518 | 168,739 | 222 | forward |
| Hypothetical protein CDS | 169,022 | 170,371 | 1,350 | reverse |
| ThiJ/PfpI domain protein CDS | 172,051 | 172,707 | 657 | forward |
| Hypothetical protein CDS | 173,007 | 173,519 | 513 | reverse |
| lipoprotein CDS | 173,852 | 174,481 | 630 | reverse |
| Hypothetical protein CDS | 174,478 | 175,581 | 1,104 | reverse |
| Hypothetical protein CDS | 178,139 | 179,374 | 1,236 | reverse |
| Hypothetical protein CDS | 180,285 | 180,782 | 498 | forward |
| Transposase CDS | 183,745 | 184,122 | 378 | forward |
| Transposase IS66 CDS | 184,134 | 184,844 | 711 | reverse |
| Hypothetical protein CDS | 185,405 | 185,839 | 435 | reverse |
| ferredoxin/ferredoxin--NADP reductase CDS | 186,640 | 186,900 | 261 | forward |
| Resolvase domain protein CDS | 187,006 | 187,677 | 672 | reverse |
| Hypothetical protein CDS | 189,926 | 190,441 | 516 | forward |
| Hypothetical protein CDS | 190,677 | 191,483 | 807 | forward |
| Hypothetical protein CDS | 192,307 | 192,924 | 618 | reverse |
| Hypothetical protein CDS | 196,545 | 196,949 | 405 | reverse |
| Hypothetical protein CDS | 196,943 | 197,599 | 657 | reverse |
| Hypothetical protein CDS | 198,400 | 198,837 | 438 | forward |
| Hypothetical protein CDS | 199,053 | 199,622 | 570 | reverse |
| Hypothetical protein CDS | 199,716 | 200,141 | 426 | reverse |
| Hypothetical protein CDS | 201,803 | 202,342 | 540 | reverse |
| Hypothetical protein CDS | 202,505 | 204,013 | 1,509 | forward |
| Hypothetical protein CDS | 204,523 | 205,509 | 987 | forward |
| Transcriptional regulator, MarR family CDS | 205,861 | 206,316 | 456 | forward |
| Hypothetical protein CDS | 206,313 | 206,825 | 513 | forward |
| pdxH CDS | 207,468 | 208,112 | 645 | forward |
| Transposase CDS | 208,247 | 209,497 | 1,251 | reverse |

#### 4.2 *In-silico* blast analysis of the sandarazol biosynthetic gene cluster

Every coding sequence in the sandarazol biosynthesis gene cluster (*szo* cluster) was extracted translated and searched with the blastp algorithm against the RefSeq non-redundant protein sequence database at NCBI. [3]

*Table S 9. Tabulated blastP results for the CDS regions present in the szo biosynthetic gene cluster*

| CDS Name | Length [AA] | Closest homologue [Organism of origin] | Identity [%] and alignment length [AA] | Proposed function | Accession Nr. |
|---|---|---|---|---|---|
| *szoA* | 242 | Cytochrome C reductase subunit [Cystobacter fuscus] | 32.2; 192 | monooxygenase reductase | WP_002625886 |
| *szoB* | 346 | 2,3 diaminopropionate biosynthesis protein SbnA [Acidobacterium sp.] | 52.5; 318 | diaminopropionate biosynthesis | PYS24491 |
| *szoC* | 282 | ACP-acyltransferase [Ralstonia solanacerum] | 50.0; 280 | malonyl-CoA transfer | WP_064046451 |
| *szoD* | 247 | Enoyl-CoA hydratase [Bacillus megatherium] | 45.4; 247 | *in-trans* dehydration | WP_097812162 |
| *szoE* | 3587 | Acyl transferase domain containing Protein [Dendosporobacter quercicolus] | 37.8; 1875 | Polyketide biosynthesis | SDL84501 |
| *szoF* | 3552 | Polyketide synthase PksM [Streptomyces leeuwenhoekii] | 33.5; 1882 | Polyketide biosynthesis | CQR60407 |
| *szoG* | 2121 | Amino acid adenylation domain containing Protein [Brevibacillus sp.] | 36.3; 2108 | Amino acid incorporation | WP_116333041 |
| *szoH* | 1462 | Type 1 Polyketide synthase [Stigmatella aurantiaca] | 37.7; 1462 | Polyketide biosynthesis | WP_075005930 |
| *szoI* | 583 | Halogenase [Burkholderia Pseudomallei] | 45.1; 531 | Halogenase | WP_119566518 |
| *szoJ* | 350 | 2,3 diaminopropionate biosynthesis protein SbnB [Acidobacterium sp.] | 45.8; 345 | diaminopropionate biosynthesis | PYS24498 |
| *szoK* | 94 | Acyl carrier protein [Duganella sp.] | 43.8; 88 | β-branching | WP_056154626 |
| *szoL* | 440 | Beta-ketoacyl synthase [Salinispora arenicola] | 40.2; 425 | β-branching | WP_018792602 |
| *szoM* | 411 | Hydroxymethylglutaryl-CoA synthase protein PyxM [Pyxidicoccus fallax] | 62.1; 411 | β-branching | ASA76639 |

| | | | | | |
|---|---|---|---|---|---|
| *szoN* | 243 | Enoyl-CoA hydratase/isomerase family protein [Streptomyces sp.] | 38.7; 243 | β-branching | WP_100840951 |
| *szoO* | 262 | Thioesterase [Sorangium cellulosum] | 37.0; 262 | β-branching | WP_012232991 |
| *szoP* | 415 | Monooxygenase [Nitrospora sp.] | 29.0; 400 | epoxidation | PLY22143 |

#### 4.3 Analysis of the shifting DH domain on *szoF*

During sandarazol Biosynthesis, we observe the consecutive formation of two β, γ double bonds in module 4 and 5 of the sandarazol biosynthesis. The *in-trans* dehydratase protein SzoD is solely able to form α, β unsaturated precursor molecules as it is the case for CurE and CorN in curacin and corallopyronin biosynthesis respectively for example. [4,5] As these other dehydratase proteins, SzoD has a standard enoyl-CoA dehydratase fold responsible for the formation of α,β unsaturated carbonyls from 3-hydroxycarbonyls. The corresponding β, γ double bonds are subsequently created by the double bond shifting DH domain on the SzoF protein. Double bond shifting DH domains are characterized by the presence of an HxxxGxxxxP motif at the start of the domain and a DxxxQ/H motif at the end of said domain. In shifting DH domains on the other hand, the HxxxGxxxxP is mostly completely present, while the DxxxQ/H is absent.

*Table S 10. Alignment of the double bond shifting sandarazol DH domain to other shifting (marked with an asterisk *) and non-shifting DH domains*

| Domain name | Motif 1 sequence | Motif 2 sequence |
|---|---|---|
| SzoF_DH* | ...HRVHGARVVPG... | ...DGATL... |
| CorL_DH* | ...HEVFGRPLFPT... | ...NGLLM... |
| CorJ_DH* | ...HTVLGQRVLLG... | ...DGVIV... |
| NspC_DH* | ...HTLLGDRVLLG... | ...NSAFL... |
| RhiE_DH* | ...HQFNHRRILLG... | ...NSAFL... |
| BaeR_DH* | ...HQFSGEPVLVG... | ...NSAYL... |
| DifK_DH* | ...HLVFGKPALMG... | ...NSCYM... |
| EryAII_DH | ...HVVGGRTLVPG... | ...DAVAQ... |
| DH_Consensus | ...HXXXGXXXXPG... | ...DXXXQ/H... |

As we can see from this analysis, the SzoF shifting DH domain seems to be an intermediate of the shifting DH domains in corallopyronin on the proteins CorL and CorJ. [4] Therefore, sandarazol biosynthesis seems to contain a dehydratation followed by double bond shifting reaction sequence as proposed in the sandarazol biosynthesis.

### 5 Genetic manipulation protocols applied to MSr10575

#### 5.1 List of primers used in this study

*Table S 11. List of all primers used in this study*

| Primers used for the creation of pBeloBac TransAT | |
|---|---|
| MSr10575 VanR NotI fwd | GAGCGGCCGCCAGTGTGATG |
| MSr10575 VanR rev | CGAATTACTGCGCGCCACAAACATATGCGTTTCCTCGCATCGTGG |
| MSr10575 TransAT fwd | CCACGATGCGAGGAAACGCATATGTTTGTGGCGCGCAGTAATTCG |
| MSr10575 TransAT NotI rev | CGGCGGCCGCGATGTCGGCTCGCCAA |
| Van primer int fwd | TCGCCGCTCTTGATCTCGCC |
| pBeloBac Sa001 int rev | CCACCCGAGCACCTCTTCCG |

#### 5.2 PCR reactions and cycler protocols

Amplification of the sandarazol cluster start as well as the vanillate promotor and repressor cassette as well as the overlap extension PCR step was done with the proofreading Phusion polymerase (Thermo scientific). PCR mix and thermocycler program can be found below.

##### 5.2.1 Thermo scientific Phusion Polymerase

**Phusion Polymerase PCR mix:**

5µL GC Buffer

2.5µL dNTP's (2mM)

1.25µL DMSO

0.5µL Primer fwd and rev (100mM)

0.2 µL Phusion DNA Polymerase

0.5µL gDNA Template (~50 ng/µL)

14.55µL dd$H_2O$

**Phusion Polymerase cycler program:**

*Table S 12: Phusion DNA Polymerase PCR program*

| Step | Time [min:s] | Temperature [°C] |
|---|---|---|
| **Initial denaturation** | 4:00 | 95 |
| | 0:30 | 98 |
| **Cycle, repeat 30x** | 0:15 | 63 |
| | 0:30 per Kbp to amplify | 72 |
| **Final elongation** | 10 | 72 |
| **Store** | forever | 8 |

#### 5.3 Creation of the pBeloBac TransAT plasmid for single crossover integration

The plasmid is based on the commercially available pBeloBac11 (NEB Biolabs). In a first step, the chloramphenicol resistance on the pBeloBac Backbone is replaced by a kanamycin resistance cassette using standard restriction cloning to form pBeloBacKan.

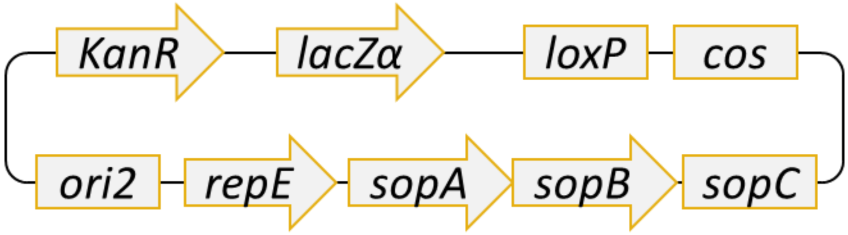

*Figure S 14. Schematic image of pBeloBac11 kan*

The Vanillic acid promotor and repressor cassette are amplified using the above described Phusion polymerase based PCR protocol and the VanR primer pair as described in the primer list from the pFPVan template plasmid described in the Paper describing the discovery of the pyxidicyclines. [6] The sandarazol plasmid cluster fragment necessary for homologous recombination based promotor exchange in front of the sandarazol cluster is amplified using the same PCR protocol from MSr10575 genomic DNA prepared according to the described phenol/chloroform DNA preparation protocol. The two described fragments are fused by overlap extension PCR using the overlapping sequences on the MSr10575 VanR rev and MSr10575 TransAT fwd primers. The fused PCR product is integrated into pBeloBac11 kan using the NotI restriction endonuclease by conventional restriction cloning, thereby replacing the lacZα gene that is not needed. The resulting plasmid is called pBeloBac TransAT. Ligations are performed with T4 DNA ligase (Thermo scientific) according to the manufacturers' instructions.

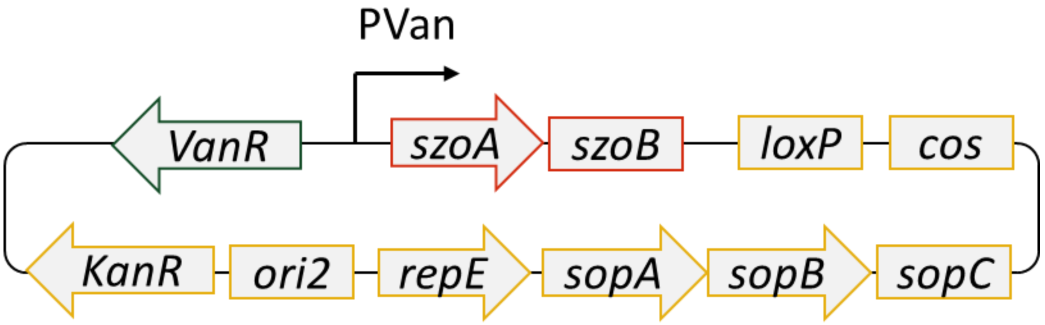

*Figure S 15. Schematic image of pBeloBac TransAT*

The plasmid is confirmed by restriction analysis and prepared in larger amounts for electro transformation into *Sandaracinus sp.* MSr10575.

#### 5.4 Transformation of *Sandaracinus sp.* MSr10575

As transformation of wild type *Sandaracinus* sp. MSr10575 was not yet established we devised an electro transformation protocol using pMycoMar Kan, a mariner element based transposon that contains a kanamycin resistance between its IR elements. Good transformation efficiency was obtained using the following electroporation protocol.

##### 5.4.1 Transformation protocol

1) Centrifuge 2ml of MSr10575 culture in S15 Medium at $OD_{600}$ of approx. 0.8 at 8000 rpm for 2 minutes with an Eppendorf tube table centrifuge.
2) Wash the residual cell pellet 2 times with 1ml autoclaved dd$H_2O$ and discard the supernatant.
3) Resuspend cells in 50µL of dd$H_2O$, add 5 - 10µL of plasmid solution (prepared from E. coli with Thermo Scientific Miniprep Kit) at a conc. of 0.3-0.4 ng/µL and transfer the solution into an electroporation cuvette.
4) Electroporation at 1000V, 400Ω, 25µF and 1mm cuvette settings for optimum electroporation efficiency.
5) Flush out the cells with 1mL fresh S15 medium and transfer the cell suspension into a 2ml Eppendorf tube.

6) Incubate the cells for 5h on a shaker thermostatic to 37°C at 300 rpm.
7) Plate the cells on Kan50 (or Kan75) S15 agar after mixing the cell suspension with 3 ml of Kan50 S15 softagar.
8) Orange, spherical clones appear in the soft agar layer after 9-14 days in the 30°C incubator.

#### 5.5 Integration of pBeloBac TransAT into the native plasmid of MSr10575

The pBeloBac TransAT plasmid is repeatedly transformed into MSr10575 wild type by electroporation according to the devised protocol. The resulting transformants are grown in kanamycin 50 µg/ml supplemented S15 medium and genomic DNA of the transformants is prepared.

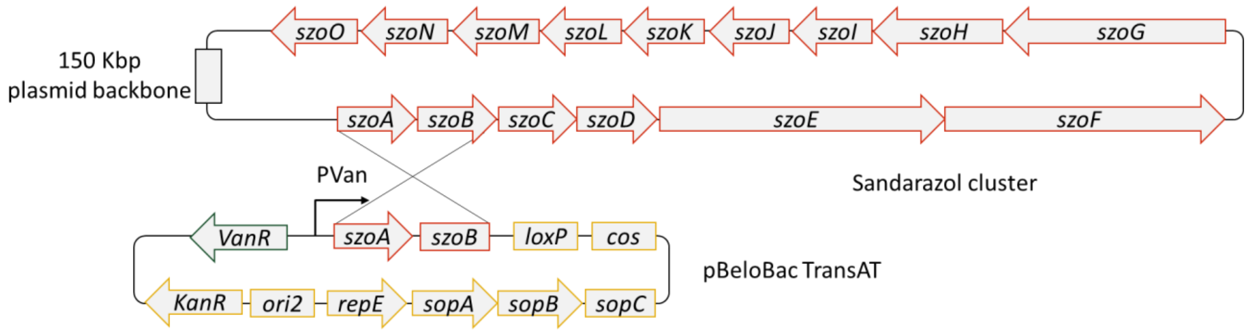

*Figure S 16. Single crossover integration event that integrates pBeloBac TransAT into the MSr10575 wild type plasmid to form pBeloBac Sa001*

Successful integration of pBeloBac TransAT into the MSr10575 wild type plasmid is checked by PCR reaction using the Van primer int fwd and pBeloBac Sa001 int rev primers. After successful homologous recombination of the pBeloBac plasmid with the 209 Kbp pBeloBac Sa001 megaplasmid, MSr10575 clones show a 2359 bp PCR product in this PCR reaction. The strain showing this PCR product is genome sequenced by Illumina to confirm successful homologous recombination on the megaplasmid to form pBeloBac Sa001.

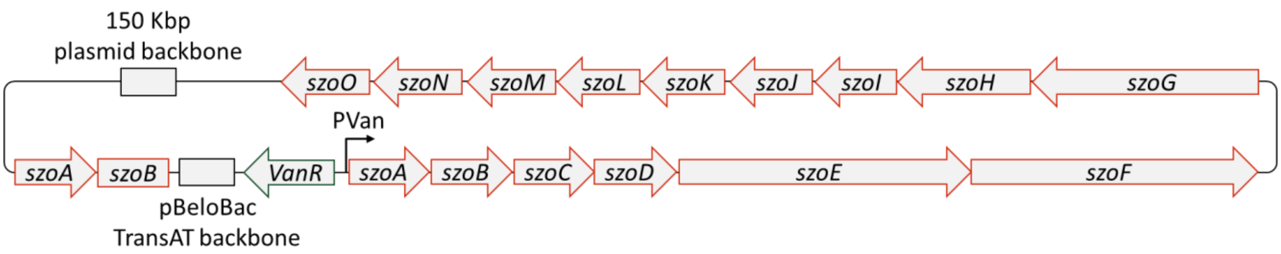

*Figure S 17. Organisation of the szo cluster on pBeloBac Sa001*

### 6 Isolation and structure elucidation of the sandarazols

#### 6.1 Isolation of the sandarazols from MSr10575 :: pBeloBac Sa001

##### 6.1.1 Cultivation and extraction

The *Sandaracinus* strain MSr10575 :: pBeloBac Sa001 is fermented in 50 ml 2SWT medium as a seed culture flasks on an Orbiton shaker at 160 rpm and 30°C. The translucid culture medium becomes orange and opaque after 7 to 11 days of fermentation. This pre-culture is used to inoculate 6 x 2L 2SWYT medium supplemented with 2 % XAD-16 resin suspension in sterilized water in 6 x 5L baffled shake flasks on an Orbiton shaker at 160 rpm and 30°C. Directly after inoculation the culture is induced with 1mM sterile filtrated K-vanillate solution. Fermentation is complete after 16 days. Cells and XAD-16 resin are harvested by centrifugation on a Beckmann Avanti J-26 XP with the JLA 8.1 rotor at 6000 rcf. Combined resin and cells are freeze dried and subsequently extracted using 2x 500 ml of a 2:1 mixture of methanol and chloroform. The combined extracts are concentrated on a rotary evaporator and partitioned between methanol and hexane. The sandarazols remain in the methanol phase. The methanol phase is dried with a rotary evaporator and the residue is partitioned between water and chloroform. The sandarazols are retained in the chloroform phase. The chloroform phase is concentrated and stored in an air-tight glass vial at -20°C until further processing.

##### 6.1.2 Centrifugal partition chromatography (CPC) pre-purification of the compounds

Pre-purification and fractionation of the different sandarazols into four fractions is achieved by CPC. We use a Gilson CPC 100 device (Gilson purification S.A.S.) connected to a Varian ProStar Solvent delivery module and a Varian ProStar 2 Channel UV detector. Fraction collection is done with a Foxy Jr. autosampler (Isco). Used biphasic solvent system is the buffered ARIZONA solvent system consisting of a 1:1:1:1 mixture of 10 mM Tris x HCl buffer pH 8.0 (Sigma), methanol (analytical grade, Fluka), EtOAc (analytical grade, fluka) and hexane (analytical grade, fluka). After equilibration of the system and loading of the sample, the aqueous phase is used as a stationary phase for 40 min at 2 ml/min and a fraction size of 4 ml. Then the organic phase is used as a stationary phase in Dual Mode for 40 min at 2 ml/min and a fraction size of 4 ml. The CPC process is followed up on the UV detector at the wavelengths 220nm and 254nm. After checking the fractions using our standard analytical LC-*hr*MS setup we create a total of 3 fractions of sandarazols, one containing the larger sandarazols of above 700 Da, one containing the chlorinated smaller sandarazols and one containing the smaller non-chlorinated sandarazols below 700 Da.

##### 6.1.3 Purification of sandarazol A and B by HPLC

Purification of sandarazol A and B is carried out using a Dionex Ultimate 3000 SDLC low pressure gradient system on a Waters Acquity CSH C18 250x10mm 5µm column with the eluents $H_2O$ 10mM ammonium formate pH 7 as A and ACN + 10mM ammonium formate pH 7 as B, a flow rate of 5 ml/min and a column thermostatic at 30°C. Sandarazol A and B are detected by UV absorption at 254 nm and purification is done by time dependent fraction collection. Separation is started with a plateau at 60% A for 2 minutes followed by a ramp to 24% A during 24 minutes and a ramp to 0% A during 1 minute. The A content is kept at 0% A for 2 minutes. The A content is ramped back to starting conditions during 30 seconds and the column is re equilibrated for 2 minutes. After evaporation, the sandarazols A and B are obtained as pale yellowish amorphous solids. After purification, LC-*hr*MS analysis shows a single peak with an exact mass of 575.344 and 577.360 $[M+H]^+$ for sandarazol A and B respectively.

##### 6.1.4 Purification of Sandarazol C by HPLC

Purification of sandarazol C is carried out using a Dionex Ultimate 3000 SDLC low pressure gradient system on a Waters Acquity CSH C18 250x10mm 5µm column with the eluents $H_2O$ 10mM ammonium formate pH 7 as A and ACN + 10mM ammonium formate pH 7 as B, a flow rate of 5 ml/min and a column thermostatic at 30°C. Sandarazol C is detected by UV absorption at 254 nm and purification is done by time dependent fraction collection. Separation is started with a plateau at 50% A for 2 minutes followed by a ramp to 20% A during 24 minutes and a ramp to 0% A during 1 minute. The A content is kept at 0% A for 2 minutes, before ramping back to starting conditions during 30 seconds and column re equilibration for 2 minutes. After solvent evaporation, sandarazol C is obtained as dark yellow amorphous solids. After purification, LC-*hr*MS analysis shows a single peak with an exact mass of 583.325 $[M+H]^+$ for sandarazol C.

#### 6.2 NMR based structure elucidation

##### 6.2.1 NMR conditions and spectroscopic data

1D and 2D NMR data used for structure elucidation of the sandarazol derivatives is acquired in Methanol-$d_4$ on a Bruker Ascend 700 spectrometer equipped with a 5 mm TXI cryoprobe ($^1$H at 700 MHz, $^{13}$C at 175 MHz). All observed chemical shift values ($\delta$) are given in in ppm and coupling constant values ($J$) in Hz. Standard pulse programs were used for HMBC, HSQC, and gCOSY experiments. HMBC experiments were optimized for $^{2,3}J_{C\text{-}H}$ = 6 Hz. The spectra were recorded in methanol-$d_4$ and chemical shifts of the solvent signals at $\delta^H$ 3.31 ppm and $\delta^C$ 49.2 ppm were used as reference signals for spectra calibration. To increase sensitivity, the measurements were conducted in a 5 mm Shigemi tube (Shigemi Inc., Allison Park, PA 15101, USA). The NMR signals are grouped in the tables below and correspond to the numbering in the schemes corresponding to every table.

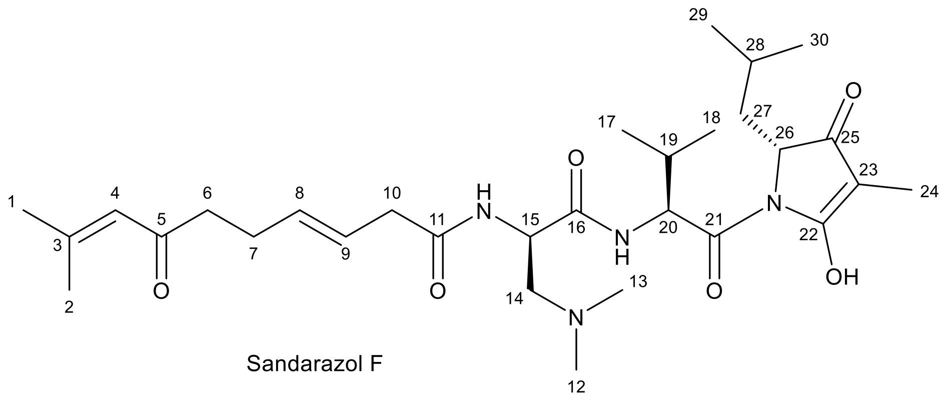

*Table S 13. NMR spectroscopic data for sandarazol F in methanol-*d4.

| # | δ $^{13}$C [ppm] | δ $^1$H [ppm], mult ($J$ [Hz]) | COSY | HMBC |
|---|---|---|---|---|
| **1** | 27.7 | 1.92 (bd, 1.1)$^+$ | 2, 4 | 2, 3, 4 |
| **2** | 20.8 | 2.13 (3H, bd, 1.1) | 1, 4 | 1, 3, 4 |
| **3** | 157.0 | - | - | - |
| **4** | 124.5 | 6.19 (1H, quin, 1.3) | 1, 2 | 1, 2, 5 |
| **5** | 202.3 | - | - | - |
| **6** | 44.1 | 2.56 (2H, bt, 7.3) | 7 | 5, 7, 8 |
| **7** | 28.0 | 2.32 (2H, bdd, 14.0, 7.0) | 6, 8 | 5, 6, 8, 9 |
| **8** | 135.1 | 5.69* (2H, m) | 7, 9 | 10 |
| **9** | 124.1 | 5.62* (2H, m) | 8, 10 | 7 |

| | | | | |
|---|---|---|---|---|
| **10** | 40.5 | 3.02 (2H, q, 6.4) | 9 | 9, 12, |
| **11** | 174.7 | - | - | - |
| **12** | 44.9 | 2.63 (3H, bs) | - | 13, 14 |
| **13** | 44.9 | 2.63 (3H, bs) | - | 12, 14 |
| **14** | 60.4 | 2.93*⁺, 3.11*⁺ | 15 | 12, 13 |
| **15** | 50.6 | 4.78* (1H, bt) | 14 | 16 |
| **16** | 171.1 | - | - | - |
| **17** | 19.8 | 0.98*⁺ | 19 | 18, 19 |
| **18** | 19.8 | 0.98*⁺ | 19 | 17, 19 |
| **19** | 31.8 | 2.19, m⁺ | 17, 18 | 17, 18 |
| **20** | 61.5 | 4.47*⁺ | - | 21 |
| **21** | 171.9 | - | - | - |
| **22** | 176.1 | - | - | - |
| **23** | 96.9 | - | - | - |
| **24** | 5.7 | 1.60 (3H, s) | - | 22, 23, 25 |
| **25** | 188.8 | - | - | - |
| **26** | 61.4 | 4.15 (1H, q, 3.0) | 27 | - |
| **27** | 39.9 | 1.92*⁺, 1.77*⁺ | 26 | 29, 30 |
| **28** | 24.8 | 1.72*⁺ | - | - |
| **29** | 23.6 | 0.97*⁺ | - | 27 |
| **30** | 23.6 | 0.97*⁺ | - | 27 |

* overlapping signals hindering determination of multiplicity and coupling constants

⁺ overlapping signals hindering integration

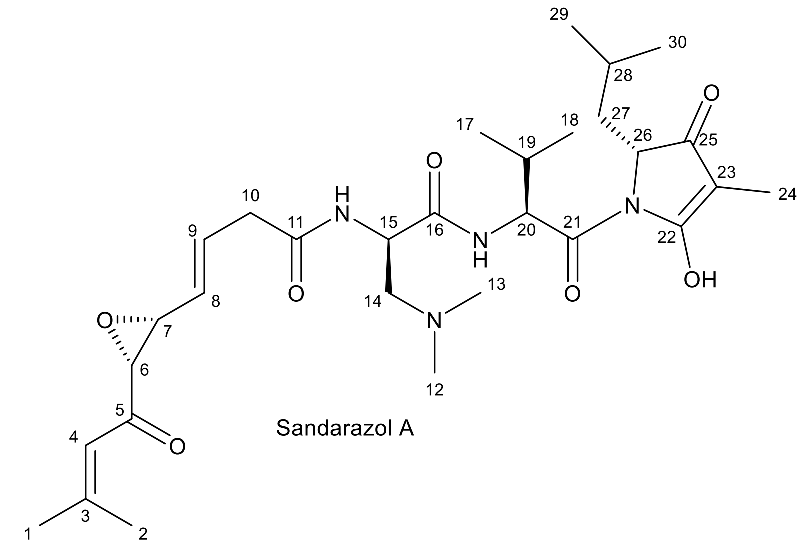

Sandarazol A

*Table S 14 NMR spectroscopic data for sandarazol A in methanol-*d$_4$*.*

| # | δ $^{13}$C [ppm] | δ $^{1}$H [ppm], mult (*J* [Hz]) | COSY | HMBC |
|---|---|---|---|---|
| **1** | 28.3 | 1.97 (3H, d, 1.2) | 2, 4 | 2, 3, 4 |
| **2** | 21.6 | 2.18$^{+}$ (d, 1.1) | 1, 4 | 1, 3, 4 |
| **3** | 162.3 | - | - | - |
| **4** | 119.6 | 6.24 (1H, quin, 1.0) | 1, 2 | 1, 2, 5 |
| **5** | 197.0 | - | - | - |
| **6** | 62.4 | 3.46* (1H) | 7 | 3, 4, 5, 7, 8 |
| **7** | 59.1 | 3.47* (1H) | 6, 8 | 6, 8, 9 |
| **8** | 131.5 | 5.45 (1H, dd, 15.4, 7.7) | 7, 9 | 6, 7, 9, 10, 11 |
| **9** | 131.6 | 6.19 (1H, bdt, 7.2) | 8, 10 | 7, 8, 10, 11 |
| **10** | 40.3 | 3.14* (2H) | 9 | 8, 9, 11 |
| **11** | 173.6 | - | - | - |
| **12** | 45.2 | 2.58 (3H, s) | - | 13, 14 |
| **13** | 45.2 | 2.58 (3H, s) | - | 12, 14 |
| **14** | 60.8 | 2.87*, 3.04* (2H) | 15 | 12, 13, 15 |
| **15** | 51.2 | 4.78 (1H, bt) | 14 | 11, 14, 16 |
| **16** | 171.6 | - | - | - |
| **17** | 20.5 | 0.98 (3H, d, 6.8) | 19 | 18, 19, 21 |
| **18** | 17.4 | 0.86 (3H, d, 6.6) | 19 | 17, 19, 21 |
| **19** | 32.2 | 2.18*$^{+}$ | 17, 18 | 17, 18, 21 |

| | | | | |
|---|---|---|---|---|
| **20** | 61.8 | 4.45*+ | - | 16, 19 |
| **21** | 171.5 | - | - | - |
| **22** | 176.5 | - | - | - |
| **23** | 97.1 | - | - | - |
| **24** | 6.0 | 1.58 (3H, s) | 26 | 22, 23, 25, 26 |
| **25** | 189.2 | - | - | - |
| **26** | 61.8 | 4.13 (1H, dd, 5.7, 3.3) | 24, 27 | 21, 22, 24, 25, 27, 28 |
| **27** | 40.0 | 1.92, 1.75 (2H, m) | 26 | 25, 26, 28, 29, 30 |
| **28** | 25.0 | 1.71 (1H, quin, 6.5) | 27, 29, 30 | 26, 27, 29, 30 |
| **29** | 24.6 | 0.81 (3H, d, 6.5) | 28, 30 | 27, 28, 30 |
| **30** | 24.2 | 0.86+ (3H, d, 6.6) | 28, 29 | 27, 28, 29 |

* overlapping signals hindering determination of multiplicity and coupling constants

+ overlapping signals hindering integration

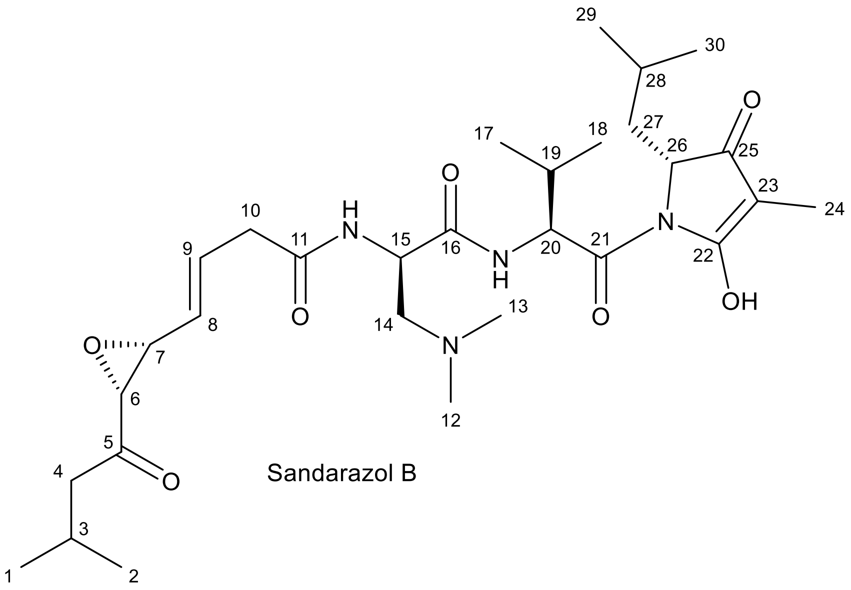

Sandarazol B

*Table S 15 NMR spectroscopic data for sandarazol B in methanol-*$d_4$.

| # | δ $^{13}$C [ppm] | δ $^{1}$H [ppm], mult (*J* [Hz]) | COSY | HMBC |
|---|---|---|---|---|
| **1** | 22.8 | 0.92*+ | 3 | 2, 3, 4 |
| **2** | 22.8 | 0.92*+ | 3 | 1, 3, 4 |
| **3** | 25.1 | 2.12*+ | 1, 2, 4 | 1, 2, 4 |
| **4** | 47.3 | 2.29*+, 2.39*+ | 3 | 1, 2, 3, 5 |
| **5** | 208.1 | - | - | - |

| | | | | |
|---|---|---|---|---|
| **6** | 61.5 | 3.47 (1H, d, 1.7) | 7 | 5, 7, 8 |
| **7** | 58.4 | 3.53* (1H, bdd, 7.8, 1.6) | 6, 8 | 5, 6, 8, 9 |
| **8** | 130.9 | 5.44 (1H, dd, 15.4, 7.7) | 7, 9 | 6, 7, 10 |
| **9** | 131.3 | 6.19* (1H, m) | 8, 10 | 7, 10 |
| **10** | 40.0 | 3.15*+ | 9 | 8, 9 |
| **11** | 173.2 | - | - | - |
| **12** | 44.9 | 2.59 (3H, s) | - | 13, 14 |
| **13** | 44.9 | 2.59 (3H, s) | - | 12, 14 |
| **14** | 60.5 | 2.87*, 3.03* (2H) | 15 | 12, 13, 15 |
| **15** | 50.8 | 4.75* (1H) | - | 11, 14, 16 |
| **16** | 171.4 | - | - | - |
| **17** | 20.1 | 0.98*+ | 18, 19 | 18, 19 |
| **18** | 17.2 | 0.86*+ | 17, 19 | 17, 19 |
| **19** | 32.2 | 2.18*+ | 17, 18 | 17, 18, 21 |
| **20** | 61.8 | 4.45*+ | - | 16, 19 |
| **21** | n.d. | - | - | - |
| **22** | 176.3 | - | - | - |
| **23** | 97.0 | - | - | - |
| **24** | 5.7 | 1.59 (3H, bs) | - | 22, 23, 25 |
| **25** | 188.8 | - | - | - |
| **26** | 61.4 | 4.15 (1H, dd, 5.6, 3.2) | 27 | 25, 27, 28 |
| **27** | 39.7 | 1.91+, 1.76+ (m) | 26 | 25, 26, 28, 29, 30 |
| **28** | 24.8 | 1.71*+ | 27, 29 | 27, 29, 30 |
| **29** | 24.4 | 0.81 (3H, d, 6.5) | 28, 30 | 27, 28, 30 |
| **30** | 24.0 | 0.86*+ | 28, 29 | 27, 28, 29 |

* overlapping signals hindering determination of multiplicity and coupling constants

+ overlapping signals hindering integration

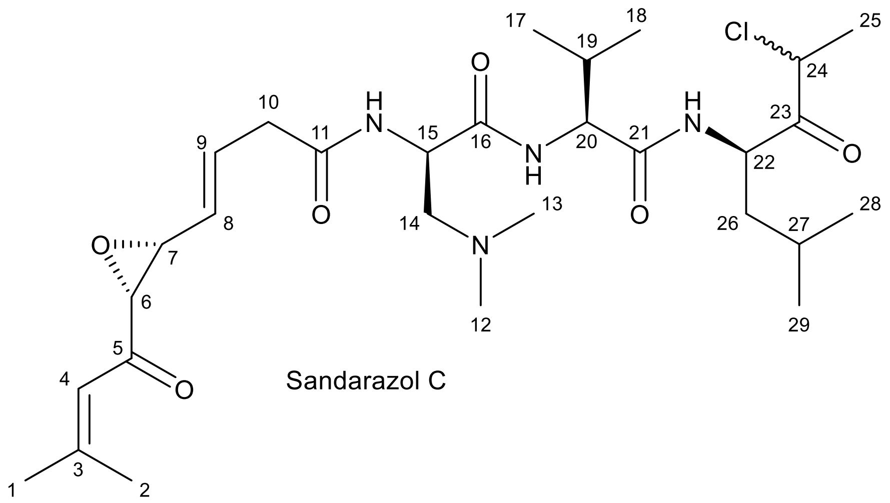

Sandarazol C

*Table S 16 NMR spectroscopic data for sandarazol C in methanol*-d$_4$.

| # | δ $^{13}$C [ppm] | δ $^{1}$H [ppm], mult (*J* [Hz]) | COSY | HMBC |
|---|---|---|---|---|
| **1** | 21.5 | 2.19 (3H, bs) | 2, 4 | 2, 3, 4, 5 |
| **2** | 28.2 | 1.97 (3H, d, 1.1) | 1, 4 | 1, 3, 4 |
| **3** | 162.3 | - | - | - |
| **4** | 119.6 | 6.24 (1H, dquin, 5.1, 1.3) | 1, 2 | 1, 2, 5 |
| **5** | 197.2 | - | - | - |
| **6** | 62.4 | 3.44* (1H) | 7 | 4, 5, 7, 8, 9 |
| **7** | 59.2 | 3.44* (1H) | 6, 8 | 5, 6, 8, 9 |
| **8** | 131.1 | 5.40 (1H, ddd, 15.4, 7.9) | 7, 9 | 6, 7, 9, 10 |
| **9** | 131.9 | 6.12 (1H, dt, 15.4, 7.1) | 8, 10 | 7, 8, 11, 10 |
| **10** | 40.0 | 3.09 (2H, m) | 9 | 8, 9, 11 |
| **11** | 173.5 | - | - | - |
| **12** | 45.8 | 2.32 (3H, s) | - | 13, 14 |
| **13** | 45.8 | 2.32 (3H, s) | - | 12, 14 |
| **14** | 61.3 | 2.59 (2H, m) | 15 | 12, 13, 15, 16 |
| **15** | 53.1 | 4.50 (1H, q, 7.3) | - | 11, 14, 16 |
| **16** | 173.9 | - | - | - |
| **17** | 19.9 | 0.99*+ | 18, 19 | 18, 19 |
| **18** | 18.2 | 0.94*+ | 17, 19 | 17, 19 |
| **19** | 31.5 | 2.21*+ | 17, 18 | 17, 18, 21 |
| **20** | 60.1 | 4.23*+ | - | 16, 19 |

| | | | | |
|---|---|---|---|---|
| **21** | 180.7 | - | - | - |
| **22** | 55.6 | 4.74 (1H, m) | 25, 26 | 23, 26, 27 |
| **23** | 204.6 | - | - | - |
| **24** | 56.3 | 4.79 (1H, m) | 25 | 25 |
| **25** | 20.7 | 1.53 (3H; dd, 8.8, 6.8) | 24 | 24, 26, 22 |
| **26** | 39.8 | 1.64 (2H, m) | 22, 27 | 22, 23, 27, 28, 29 |
| **27** | 26.2 | 1.72 (1H, m) | 26, 28, 29 | 22, 26, 28, 29 |
| **28** | 23.8 | 0.96*+ | 27, 29 | 26, 27, 29 |
| **29** | 21.7 | 0.93*+ | 27, 28 | 26, 27, 28 |

* overlapping signals hindering determination of multiplicity and coupling constants

+ overlapping signals hindering integration

##### 6.2.2 Structure elucidation based on relevant chemical shifts and correlations

The $^{1}$H spectrum of sandarazol A showed nine methyl groups at $\delta^{1}$H = 0.81 (3H, d, 6.5), 0.86 (6H, d, 6.6), 0.98 (3H, d, 6.8), 1.58 ppm (3H, s), 1.97 (3H, d, 1.2), 2.18 (d, 1.1) and 2.58 (6H, bs). Furthermore, the spectrum revealed three methylene groups at $\delta^{1}$H = 1.75/1.92 (2H, m), 2.87/3.04 (2H) and 3.14 (2H) ppm. Two methine groups were found at $\delta^{1}$H = 4.78 (1H, bt) and 4.45 ppm, whose characteristic chemical shift revealed them as α-protons. COSY correlations of the methine group at $\delta^{1}$H = 4.45 ppm to another methine group at 2.19 ppm and HMBC correlations to two of the nine methyl groups at 0.98 and 0.86 ppm, as well as HMBC correlations of those groups to a quaternary carbon at $\delta^{13}$C = 171.5 ppm revealed the first amino acid to be valine. The second α-proton at $\delta^{1}$H = 4.78 ppm showed COSY correlations to a diastereotopic methylene group at $\delta^{1}$H = 2.87/3.04, which shows HMBC correlations to two methyl groups at $\delta^{1}$H = 2.58 ppm and one quaternary carbon at $\delta^{13}$C = 171.6 ppm. The $^{13}$C downfield chemical shift of the methylene group at $\delta^{13}$C = 60.8 ppm, as well as the downfield shift of the two methyl groups at $\delta^{13}$C = 45.2 ppm, besides the correlations described beforehand, revealed the second amino acid to be 2-Amino-3-(N,N-dimethyl amino)-propanoic acid. HMBC correlations of the valine methine group at $\delta^{1}$H = 4.45 ppm to the acid function of the 2-Amino-3-(N,N-dimethyl amino)-propanoic acid suggested their amide connection at valine the α-proton.

HMBC correlations of the 3*N*,3*N*-dimethyl-2,3-diaminopropionic acid α-proton to a quaternary carbon at $\delta^{13}$C = 173.6 ppm suggested its participation in another amide function. HMBC and COSY correlations of two downfield methine groups at $\delta^{1}$H = 6.19 and 5.45 ppm, as well as one methylene group at $\delta^{1}$H = 3.14 ppm revealed unsaturation in β position of the PKS part of the molecule. The aliphatic double bond proton at $\delta^1$H = 5.45 ppm showed COSY correlations to a methine group at $\delta^1$H = 3.47 ppm which could, along with its neighboring methine group at $\delta^1$H = 3.46 ppm, be determined as epoxide function due to their characteristic chemical shifts. The downfield chemical shift of the quaternary carbon at $\delta^{13}$C = 197.0 ppm which shows HMBC correlations to the two epoxide methines revealed the next functional group as a ketone. This ketone shows further HMBC correlations to one aliphatic double bond proton at $\delta^1$H = 6.24 ppm, as well as two methyl groups at $\delta^1$H = 2.18 and 1.97 ppm. Only one additional quaternary carbon at $\delta^{13}$C = 162.3 ppm showed further HMBC correlations to the two methyl groups and the aliphatic double bond proton, revealing the N-terminal end of sandarazol A here.

The carboxylic acid of valine shows correlations to a methine group at $\delta^1$H = 4.13 ppm, suggesting elongation through an amide bond in this part of the molecule as well. The methine group exhibits COSY correlations to a diastereotopic methylene group at $\delta^1$H = 1.92/1.75 ppm and HMBC correlations to one methine and two methyl groups at $\delta^1$H = 1.71, 0.86 and 0.81 ppm. Their characteristic chemical shifts as well as further correlations to a quaternary carbon at $\delta^{13}$C = 189.2 ppm revealed this part of the molecule to derive from leucine. The downfield chemical shift of the quaternary carbon at $\delta^{13}$C = 189.2 ppm and further HMBC correlations of the methine group at $\delta^1$H = 4.13 ppm to two quaternary carbons at $\delta^{13}$C = 97.1 and 176.5 as well as a methyl group at $\delta^1$H = 1.58 ppm suggested further elongation of the molecule in C-terminal direction of the leucine. The characteristic chemical shift of the methyl group at $\delta^1$H = 1.58 ppm and $\delta^{13}$C = 6.0 ppm, alongside with the sum formula of sandarazol A revealed, that the elongated leucine forms a five-membered heterocycle at the nitrogen atom of the amide bond to valine.

Sandarazol B shows high similarity to sandarazol A, as it only differs in the terminal part of the PKS part of the molecule. Instead of the quaternary carbon at $\delta^{13}$C = 162.3 ppm and the aliphatic double bond proton at $\delta^1$H = 6.24 ppm, the HSQC spectrum of sandarazol B reveals one diastereotopic methylene and one methine group at $\delta^1$H = 2.29/, 2.39 and 2.12 ppm, indicating saturation of the terminal double bond.

In sandarazol C, the PKS part is the same as in sandarazol A. It is fused to 3*N*,3*N*-dimethyl-2,3-diaminopropionic acid and valine as well. The leucine derived heterocycle however is missing in sandarazol C. The methine group showing HMBC correlations to the carboxylic acid of valine is downfield shifted to $\delta^1$H = 4.74 ppm, but the subsequent leucine part shows similar chemical shifts and correlations like in sandarazol A.. The methyl group at $\delta^{13}$C = 6.0 in sandarazol A however, is shifted to $\delta^{13}$C = 20.7 ppm in sandarazol C, revealing its terminal position of an aliphatic chain now. It exhibits correlations to a methine group at $\delta^1$H = 4.79 ppm and a quaternary carbon at $\delta^{13}$C = 204.6 ppm, a ketone deriving from the leucine carboxylic acid. The sum formula of sandarazol C in line with the chemical shifts in this part of the molecule revealed chlorination at the methine group at $\delta^1$H = 4.79 ppm.

Sandarazol F also shows high similarity to sandarazol A, as it only differs in the middle part of the PKS part of the molecule. The two epoxide bearing methines are replaced by two methylene groups at $\delta^1$H = 2.56 and 2.32 ppm in sandarazol F, revealing missing epoxidation of the molecule.

#### 6.3 Elucidation of the absolute stereochemistry

##### 6.3.1 Marfey's analysis protocol

To determine absolute configurations of amino acids the derivatization method Marfey's analysis is employed. Approximately 50 µg of the peptide to analyze is dried in a glass vial at 110°C. 100 µL 6N HCl are added, the vial is filled with $N_2$ gas and incubated at 110°C for 45 minutes to 20 hours, depending on peptide stability, for peptide hydrolysis. The vial is subsequently opened and containing fluid is dried at 110 °C. The residue is taken up in 100 µL milliQ $H_2O$ and split into two 2 ml Eppendorf tubes. To each tube 20 µL of 1N $NaHCO_3$ is added as well as 20 µL of D- respective L-FDLA as a 1% solution in Acetone. The mixture is incubated at 40°C and centrifuged at 700 rpm. 10 µL 2N HCL solution is added to quench the reaction and 300 µL ACN is added to obtain a total volume of 400 µL. The Eppendorf tube is centrifuged at 15000 rpm in a table centrifuge and transferred into a conical HPLC vial.

Separation is done on Waters Acquity BEH C-18 1.7µ, 100mm x 2.1mm at the maxis 4G coupled UHPLC-*hr*MS system. Eluents are water dd. + 0.1% formic acid as A and acetonitrile dest. + 0.1% formic acid as B. Column is thermostatic at 45°C and volume flow rate is set to 600 µL/min. The gradient is a linear gradient from 5 to 10% B over 1 min, followed by a linear gradient to 35% B over 14 min, followed by a linear gradient over 6 min to 50% B and linear gradient to 80% B. 80% B are held for one minute and the column is reconditioned with a linear gradient of 1 min to 5% B. Detection of the Marfey's derivatives is done by mass spectrometry and UV detection at 340 nm. Identification of the correct stereochemistry of the amino acid is done via comparison of retention times to FDLA derivatized standards. [7]

##### 6.3.2 Marfey's analysis of sandarazol A

Sandarazol A poses some significant challenges to Marfey's analysis as on the one hand, Leucine has to be released from the compound following a retro aldol decomposition of the terminal heterocyclic system as seen in Scheme S 1.

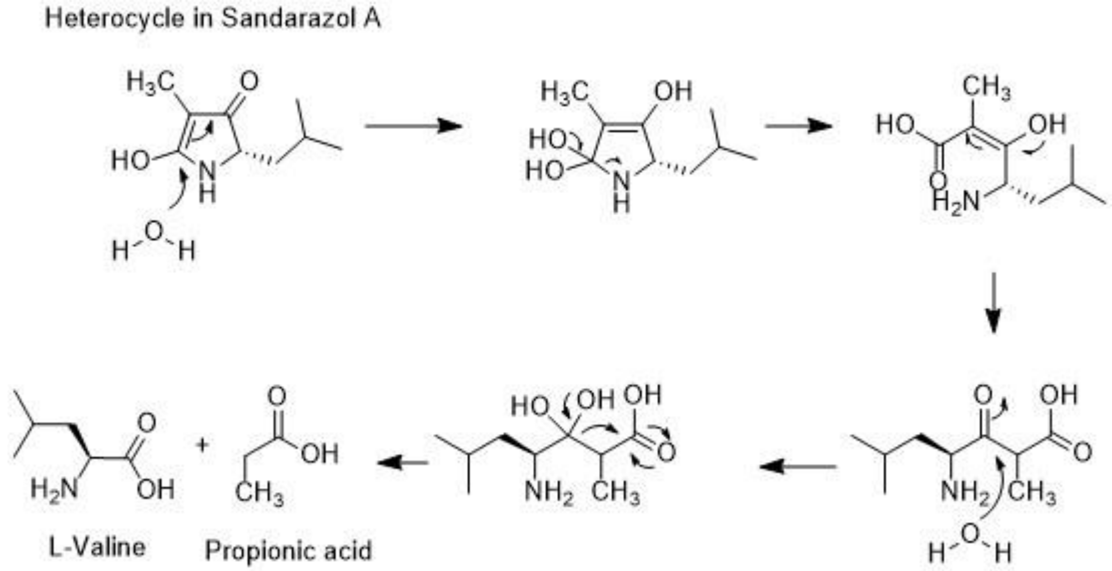

*Scheme S 1. Acid catalyzed retro aldol reaction that releases L-Valine from the terminal heterocycle in sandarazol A*

This reaction requires rather harsh hydrolysis conditions. On the other hand configuration assignment of the 2-Amino-3-(N,N-dimethyl amino)-propanoic acid (L-DimeDap) has to be done following very mild hydrolysis conditions as this amino acid is very prone to racemization even under acidic conditions. Thus, one hydrolysis reaction is set up according to the protocol above using 6N HCl for 15 min at 110°C to avoid DimeDap racemization and one Marfey's reaction is set up using conc. (37%, 12N) HCl for 12 h at 110°C to get sufficient turnover in the retro aldol reaction. Used standard substances are L-Valin and L-Leucin (Sigma Aldrich) as well as Boc-L-DimeDap (Boc-(2S)-2-Amino-3-(N,N-dimethyl amino)-propanoic acid, Thermo Fischer) that are all commercially available. It is worth noticing that the Boc protection group will be removed in the acid treatment of the reference compound and therefore the reference amino acid in the Marfey's reaction is L-DimeDap. The retention times for all Marfey's derivatized amino acids are tabulated below.

*Table S 17. Retention times and exact masses of the Marfeys derivatives.*

| Sample/Standard | FDLA derivative | Exact mass [M+H]$^+$ | Retention time | Detected species |
|---|---|---|---|---|
| L-Leu | D-FDLA | 420.204 Da | 21.04 min | L-Leu-D-FDLA |
| L-Leu | L-FDLA | 420.204 Da | 17.67 min | L-Leu-L-FDLA |
| L-Val | D-FDLA | 412.189 Da | 19.57 min | L-Val-D-FDLA |
| L-Val | L-FDLA | 412.189 Da | 16.00 min | L-Val-L-FDLA |
| Boc-L-DimeDap | D-FDLA | 427.201 Da | 8.96 min | L-DimeDap-D-FDLA |
| Boc-L-DimeDap | L-FDLA | 427.201 Da | 9.23 min | L-DimeDap-L-FDLA |
| Sandarazol A | D-FDLA | 420.204 Da | 21.03 min | L-Leu-D-FDLA |
| Sandarazol A | D-FDLA | 412.189 Da | 19.56 min | L-Val-D-FDLA |
| Sandarazol A | D-FDLA | 427.201 Da | 9.23 min | D-DimeDap-D-FDLA |
| Sandarazol A | L-FDLA | 420.204 Da | 17.67 min | L-Leu-L-FDLA |
| Sandarazol A | L-FDLA | 412.189 Da | 16.02 min | L-Val-L-FDLA |
| Sandarazol A | L-FDLA | 427.201 Da | 8.94 min | D-DimeDap-L-FDLA |

From the measured retention times as well as from the fact that enantiomers cannot differ in retention time on an achiral HPLC column as the one we used, we can conclude sandarazol A to contain D-DimeDap, L-Val and L-Leu. This is also in accordance with the modules arrangement in the sandarazol biosynthetic gene cluster as the only module containing an epimerization domain is the one introducing DimeDap into the sandarazols.

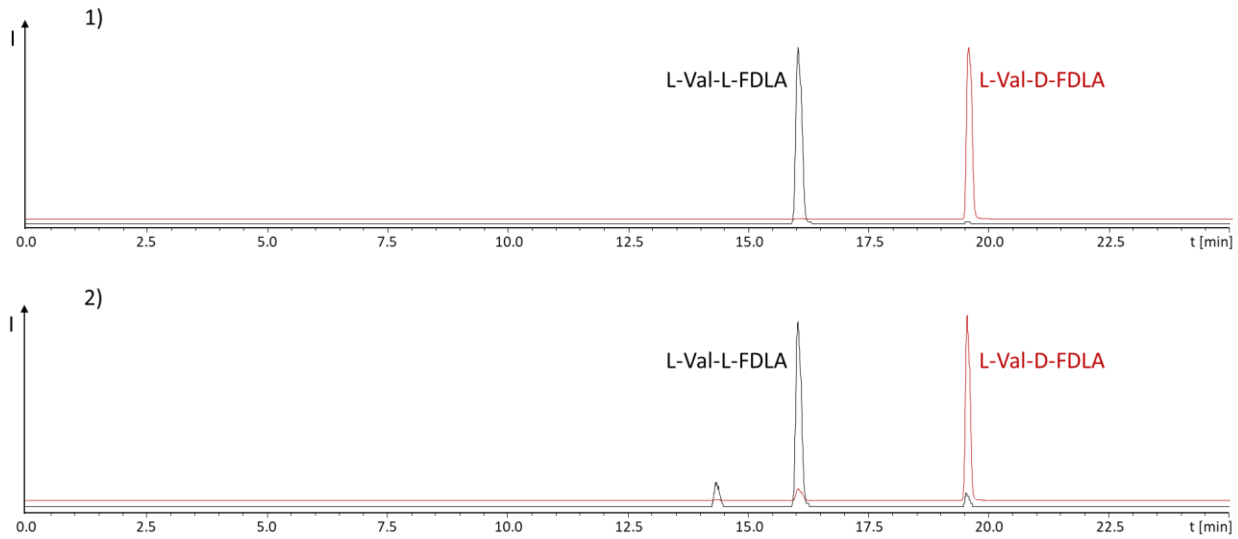

*Figure S 18. 1) Derivatization of L-Val standard with L-FDLA (black trace) and D-FDLA (red trace). 2) Derivatization of hydrolyzed sandarazol with L-FDLA (black trace) and D-FDLA (red trace). The traces represent EIC chromatograms at 412.189 Da. Peak retention time comparison reveals the Valine present in sandarazol to be L- configured.*

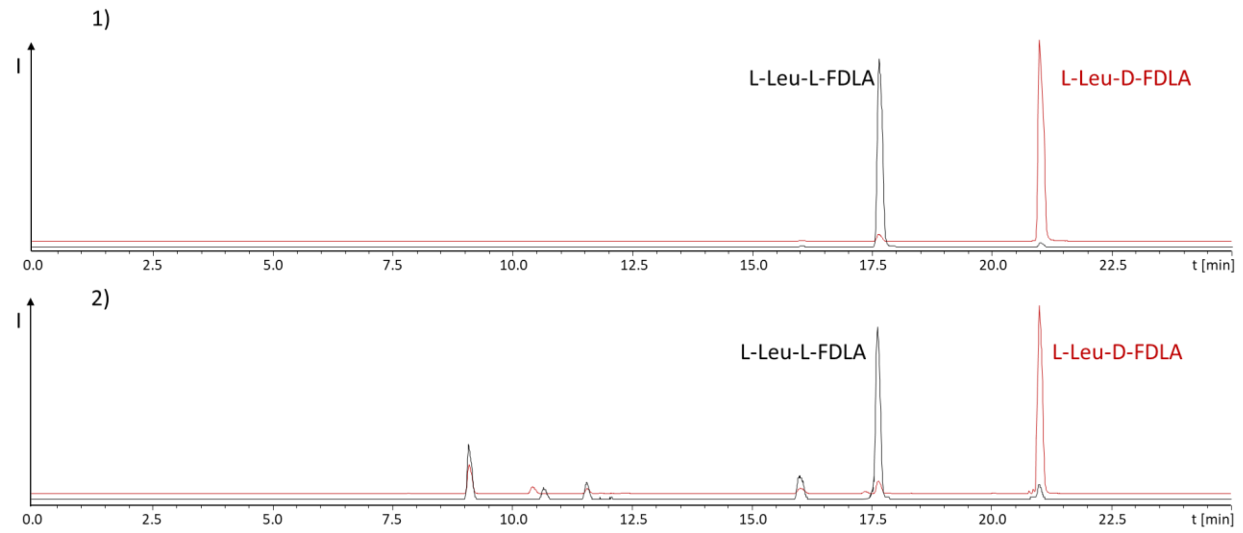

*Figure S 19. 1) Derivatization of L-Leu standard with L-FDLA (black trace) and D-FDLA (red trace). 2) Derivatization of hydrolyzed sandarazol with L-FDLA (black trace) and D-FDLA (red trace). The traces represent EIC chromatograms at 420.204 Da. Peak retention time comparison reveals the Leucine present in sandarazol to be L- configured.*

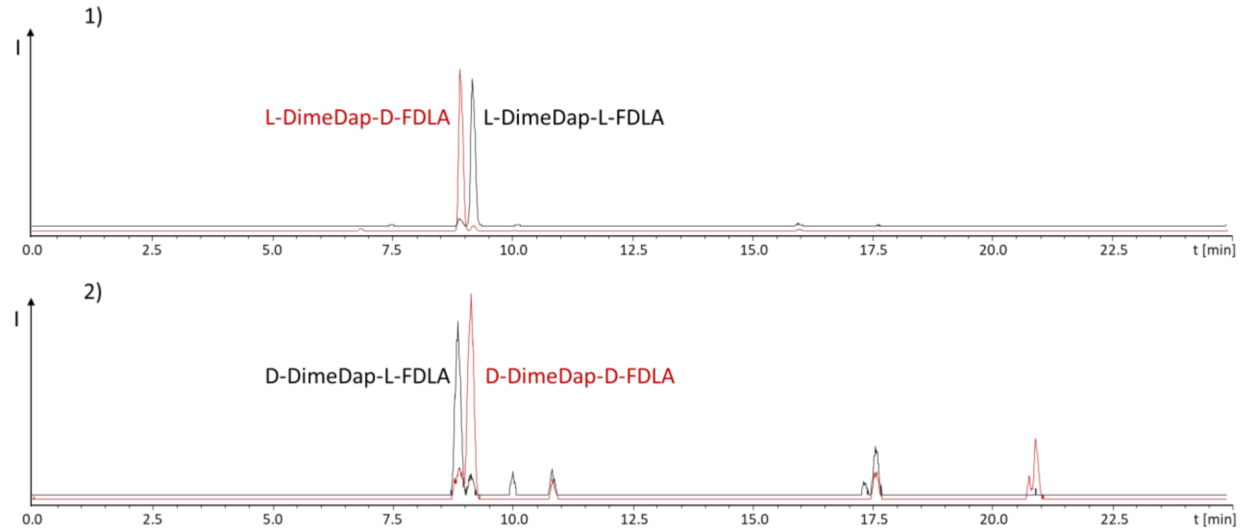

*Figure S 20. 1) Derivatization of L-DimeDap standard with L-FDLA (black trace) and D-FDLA (red trace). 2) Derivatization of hydrolyzed sandarazol with L-FDLA (black trace) and D-FDLA (red trace). The traces represent EIC chromatograms at 412.189 Da. Peak retention time comparison reveals the DimeDap present in sandarazol to be D-configured as it shows inverse identity to the peaks in the L-DimeDap standards and enantiomers elute at the same retention time.*

##### 6.3.3 The stereo center on the chlorinated carbon

The chlorinated sandarazols have one additional stereo center at the chlorinated carbon in the molecule. Unfortunately, we were not able to determine its original stereochemistry as this carbon racemizes fast, even in the culture broth. Sandarazol C and D always show two distinct LC-MS peaks of a constant peak area ratio that represent the R- and the S-configuration at this carbon atom. After isolation of both of these peaks by LC we again get both peaks with the same peak area ratio meaning the isomerization happens fast at room temperature. Since the non-chlorinated sandarazols do not show this behavior, this splitting in the LC must occur at the stereo center at the chlorinated carbon that racemizes and creates diastereomers in that process.

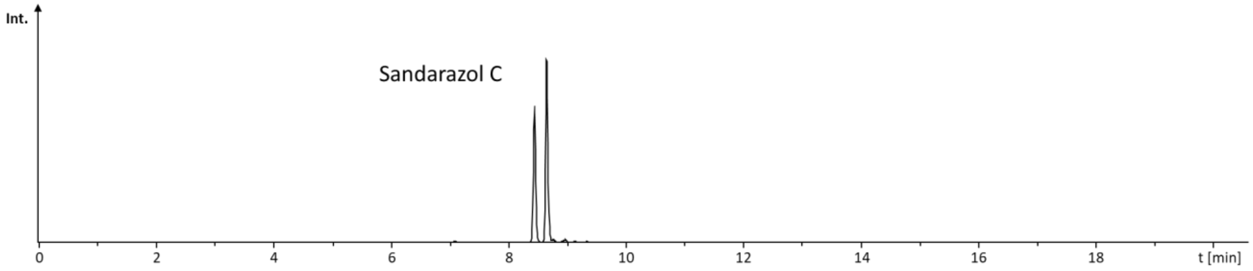

*Figure S 21. Chromatogram displaying the equilibrium distribution of sandarazol C under HPLC conditions, the chromatogram displays an extracted ion chromatogram at 583.325 Da to visualize sandarazol C with R- and S- config at the Cl bound carbon atom.*

For that reason we concluded that natural sandarazol C and D are mixtures of both occurring epimers. The rapid racemization process did not allow us to assign the earlier or the latter peak to either the S- or R- form as racemization occurs faster than compound isolation.

##### 6.3.4 Stereo chemical elucidation of the epoxide stereo center

Most sandarazols except for sandarazols E and F feature an epoxy ketone in their chemical structure. The ketone is introduced by an asymmetric FAD monooxigenase type epoxidation putatively catalyzed by the flavin containing protein SzoP and its reductase SzoA. [8] As there are no stereochemistry prediction tools available to predict the stereo chemical outcome of such a reaction based on the protein sequence, the stereochemistry of the epoxide has to be elucidated by chemical methods. While there are methods available to assign the stereochemistry of a secondary alcohol to its respective stereochemistry, the tool for direct stereo chemical elucidation of epoxides is limited to X-ray crystal structure analysis. Still, as nucleophilic ring opening of epoxide structures is always *anti* based on its mechanism, an epoxide transfers its stereochemistry to the corresponding alcohol in this type of an epoxide ring opening reaction. The resulting alcohol can then be structurally elucidated using Mosher's esterification method. [9] To use this, sandarazol A is transformed into methoxy-sandarazol A in a Lewis catalyzed epoxide ring opening reaction. The resulting methoxy-sandarazol A is purified by HPLC on a Phenomenex C18 biphenyl column under nitrogen gas using water + 10mM Ammonium formate (AmFo) and methanol + 10mM AmFo as eluents A and B. Separation is started with a plateau at 50% A for 2 minutes followed by a ramp to 32% A during 24 minutes and a ramp to 0% A during 1 minute. The A content is kept at 0% A for 2 minutes. The A content is ramped back to starting conditions during 30 seconds and the column is re equilibrated for 2 minutes. After evaporation, the methoxy-sandarazol A is obtained as pale yellowish amorphous solid.

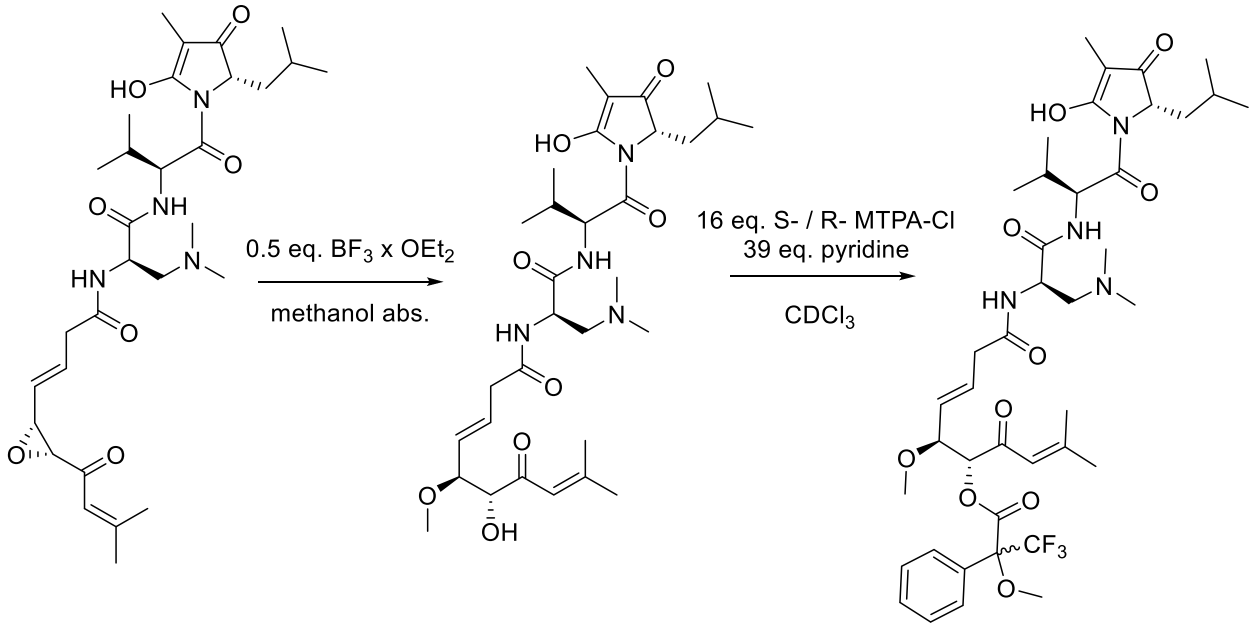

*Figure S 22. Epoxide ring opening reaction and Mosher's esterification for determination of the sandarazoles' epoxide stereochemistry.*

The resulting methoxy-sandarazol A features only one alcohol function that can be used in Mosher's method that inherited its stereochemistry from the epoxide. [9,10] Methoxy-sandarazol A is transferred into the corresponding S- and R- Mosher's ester by incubation with 16 eq. of the corresponding MTPA chloride and 39 eq. pyridine in chloroform. Configuration of the hydroxyl-group and therefore configuration of the epoxide is determined by the proton shifts of the S- and R- Moshers esters to be R.

*Table S 18. Chemical shifts of S- and R-ester of methoxy-sandarazol A and resulting assignment of R1 and R2.*

| Functional group | No. | *δ* S-ester [ppm] | *δ* R-ester [ppm] | Δ ppm | Hz (700 MHz) | |
|---|---|---|---|---|---|---|
| **$CH_3$** | 1 | 2.1 | 2.14 | -0.04 | -28 | R1 |
| **$CH_3$** | 2 | 1.85 | 1.96 | -0.11 | -77 | -259 |
| **CH** | 3 | 6.09 | 6.31 | -0.22 | -154 | |
| **CH** | 4 | 5.43 | 5.39 | 0.04 | 28 | R2 |
| **CH** | 5 | 4.13 | 4.05 | 0.08 | 56 | +553 |
| **$CH_3$** | 6 | 3.32 | 3.26 | 0.06 | 42 | |
| **CH** | 7 | 5.55 | 5.46 | 0.09 | 63 | |
| **CH** | 8 | 5.89 | 5.82 | 0.07 | 49 | |
| **$CH_2$** | 9 | 3.11 | 3.04 | 0.07 | 49 | |
| **CH** | 10 | 5.01 | 5.01 | 0 | 0 | |
| **$CH_2$** | 11a | 3.39 | 3.38 | 0.01 | 7 | |
| **$CH_2$** | 11b | 3.55 | 3.55 | 0 | 0 | |
| **$CH_3$** | 12 | 2.95 | 2.96 | -0.01 | -7 | |
| **$CH_3$** | 13 | 2.95 | 2.96 | -0.01 | -7 | |
| **CH** | 14 | 5.64 | 5.64 | 0 | 0 | |
| **CH** | 15 | 2.19 | 2.19 | 0 | 0 | |
| **$CH_3$** | 16 | 1.01 | 1.01 | 0 | 0 | |
| **$CH_3$** | 17 | 0.91 | 0.9 | 0.01 | 7 | |
| **$CH_3$** | 18 | 1.69 | 1.77 | -0.08 | -56 | |
| **CH** | 19 | 4.93 | 4.94 | -0.01 | -7 | |
| **$CH_2$** | 20a | 1.6 | 1.45 | 0.15 | 105 | |
| **$CH_2$** | 20b | 1.87 | 1.86 | 0.01 | 7 | |
| **CH** | 21 | 1.45 | 1.32 | 0.13 | 91 | |
| **$CH_3$** | 22 | 0.74 | 0.65 | 0.09 | 63 | |
| **$CH_3$** | 23 | 0.71 | 0.58 | 0.13 | 91 | |

### 7 NMR spectra employed in sandarazol structure elucidation

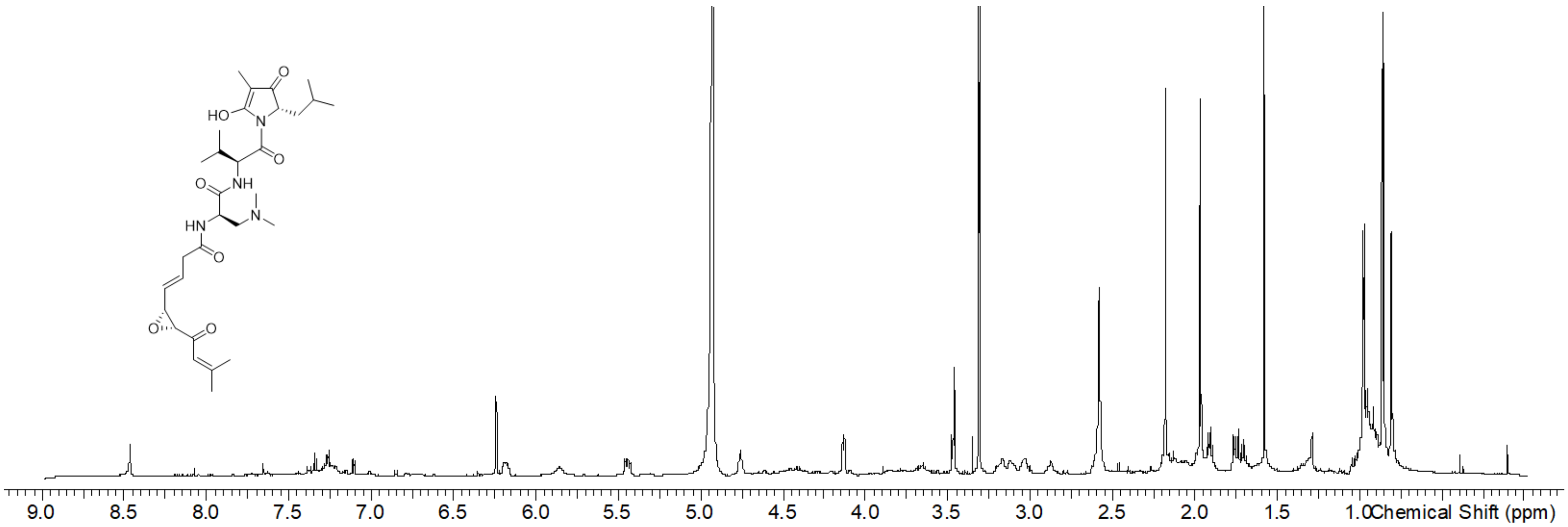

*Figure S 23. $^{1}$H spectrum of sandarazol A in* MeOD-d4.

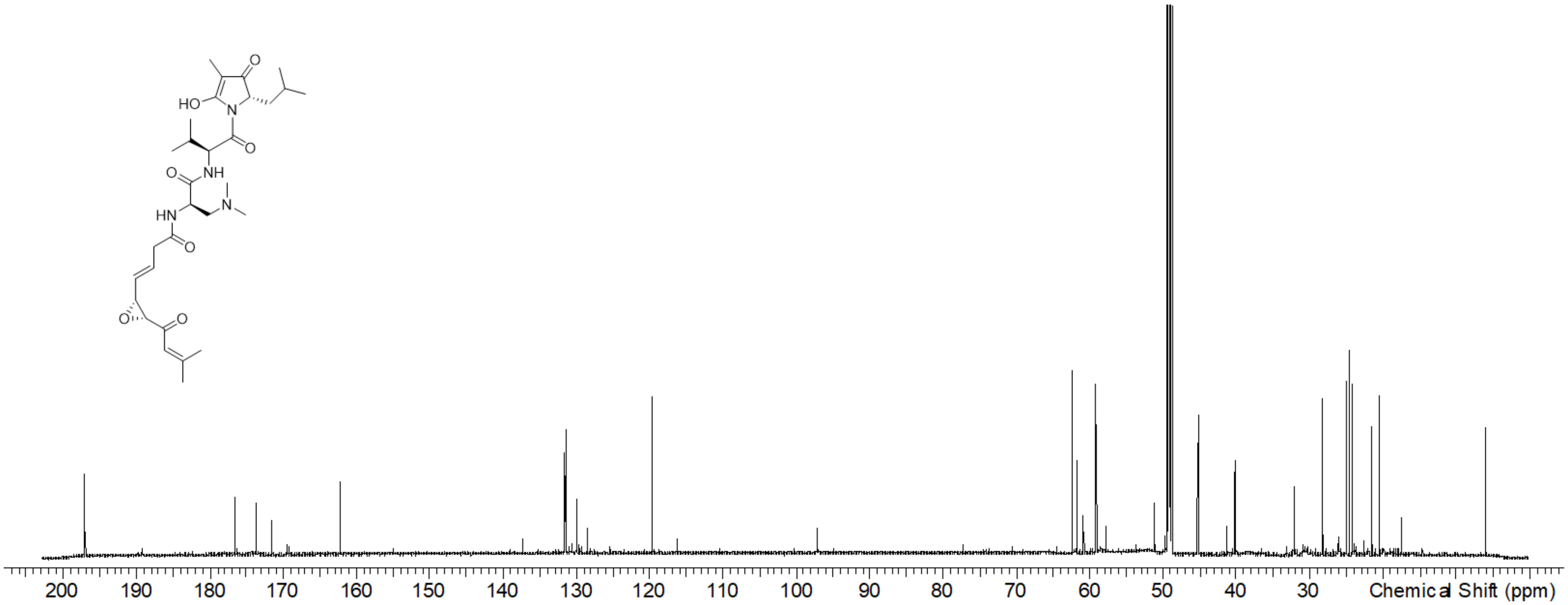

*Figure S 24. [13]C spectrum of sandarazol A in* MeOD-d4.

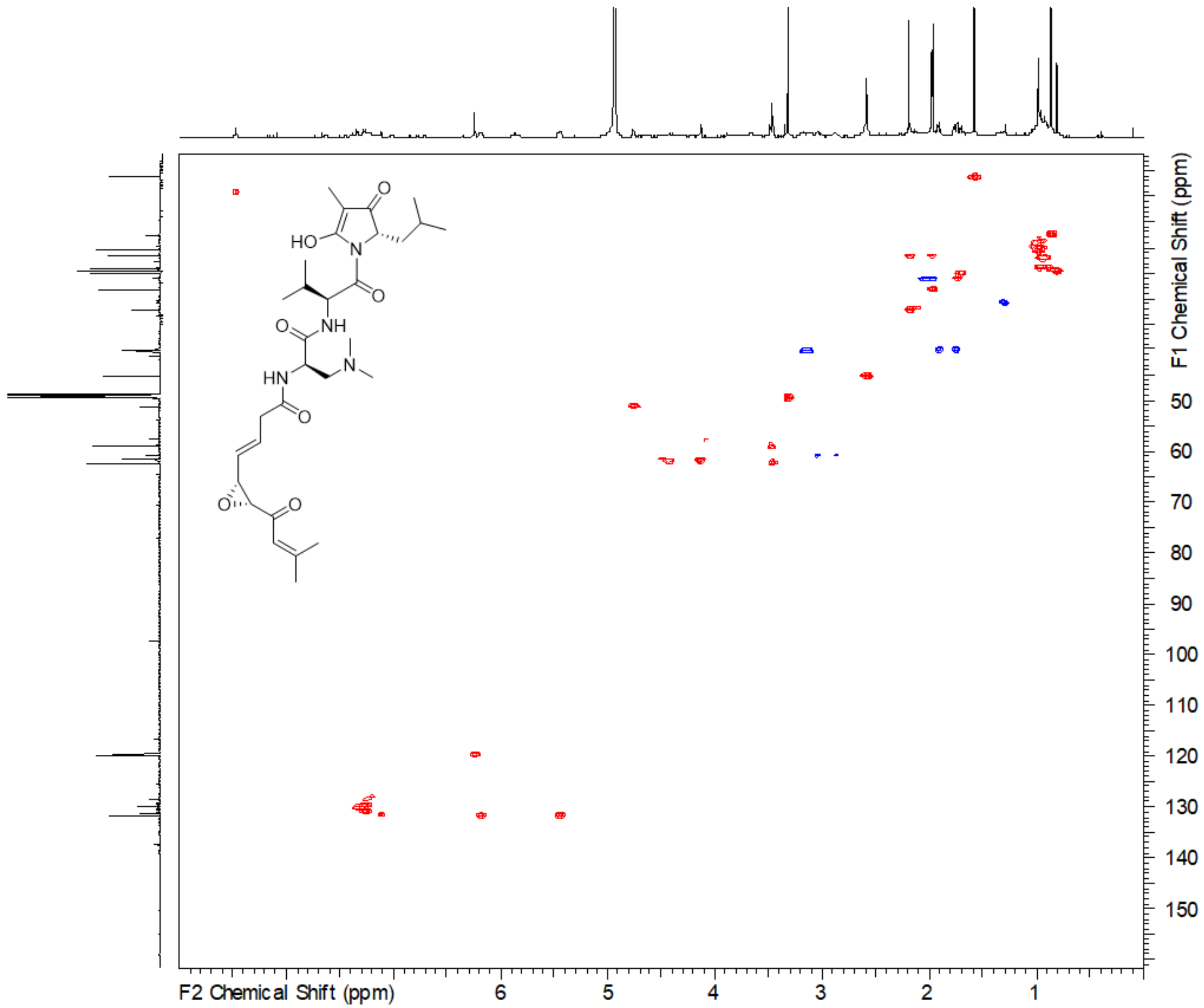

*Figure S 25. HSQC spectrum of sandarazol A in* MeOD-d4.

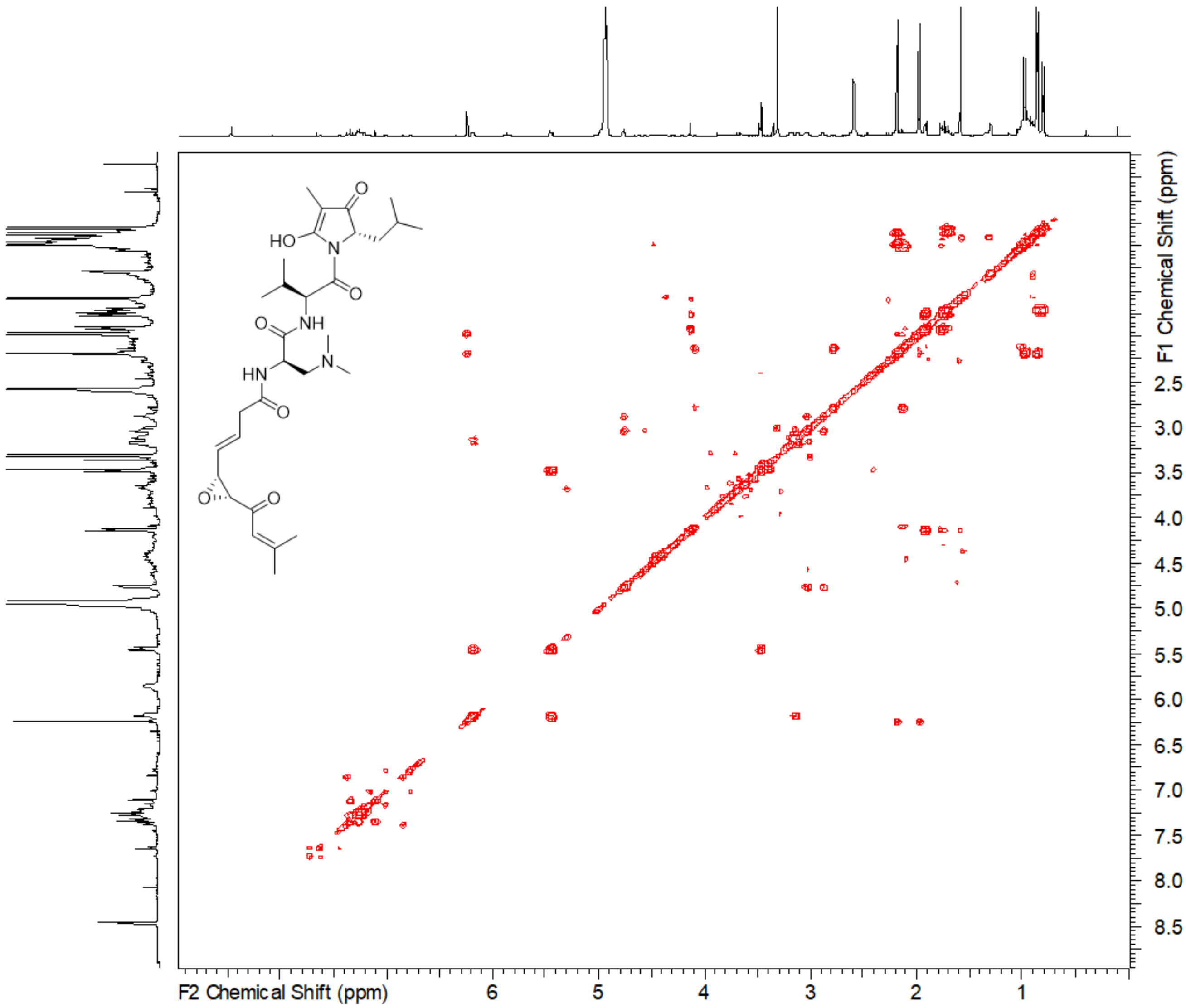

*Figure S 26. COSY spectrum of sandarazol A in* MeOD-d4.

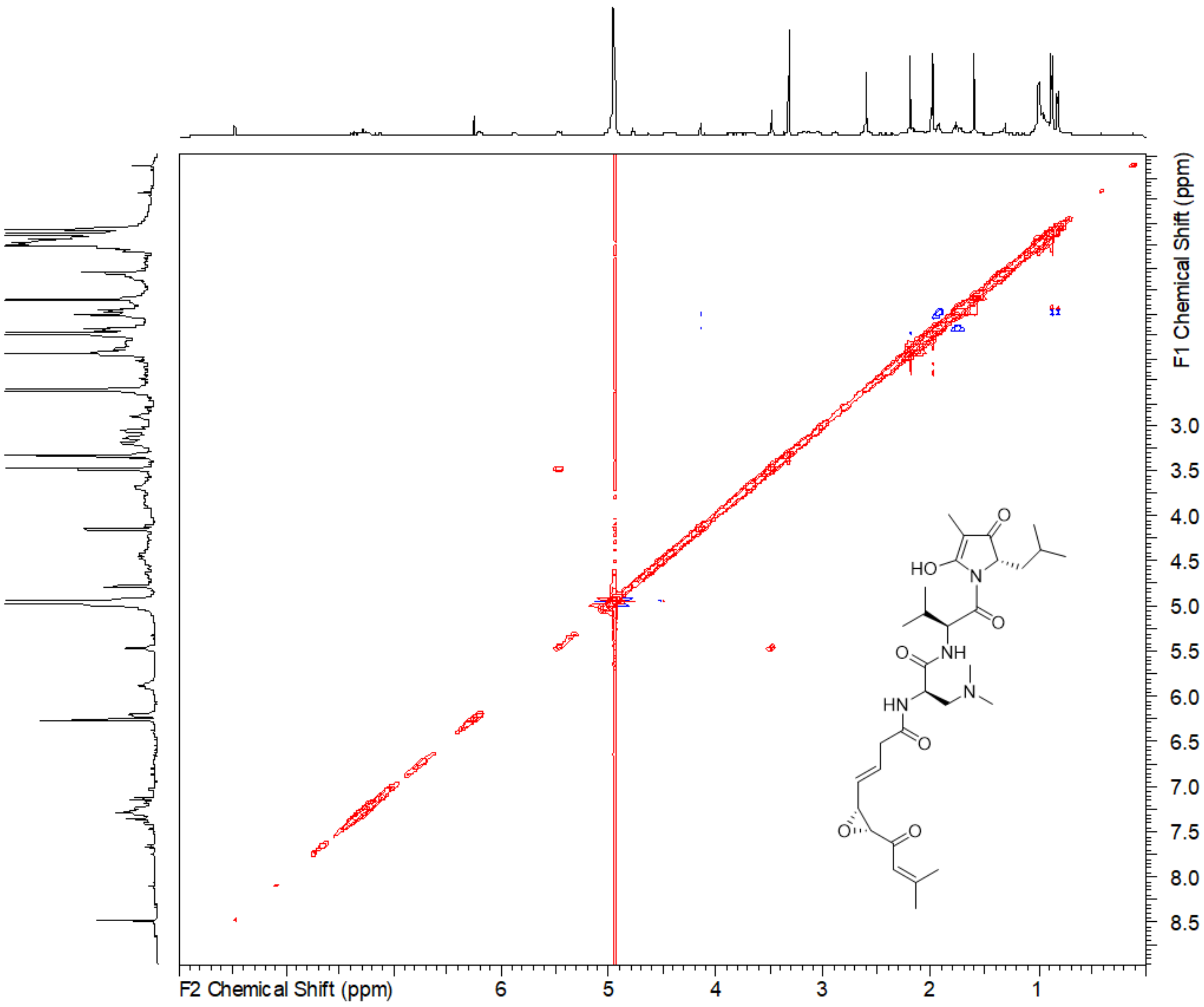

*Figure S 27. ROESY spectrum of sandarazol A in* MeOD-$d_4$.

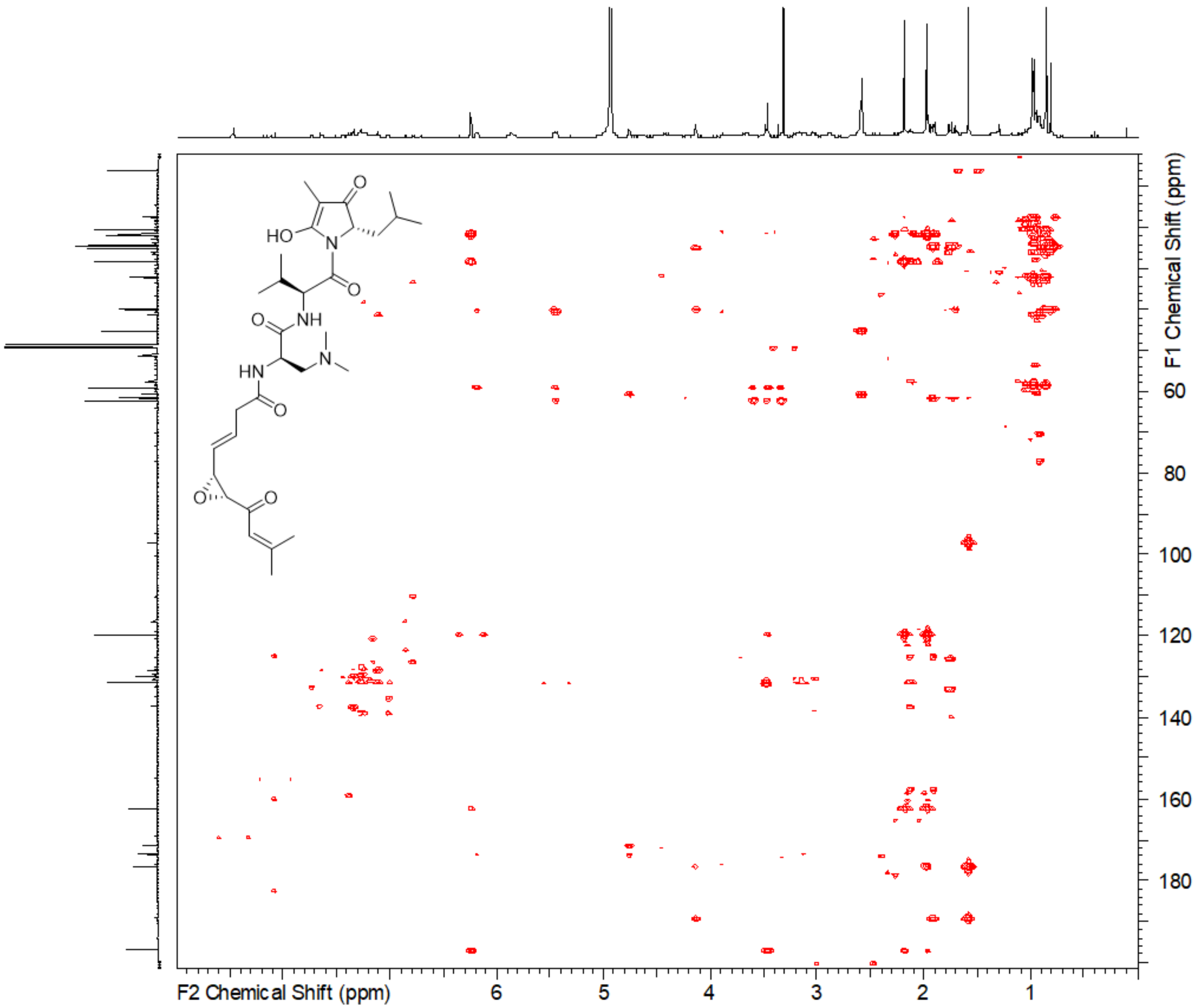

*Figure S 28. HMBC spectrum of sandarazol A in* MeOD-d4.

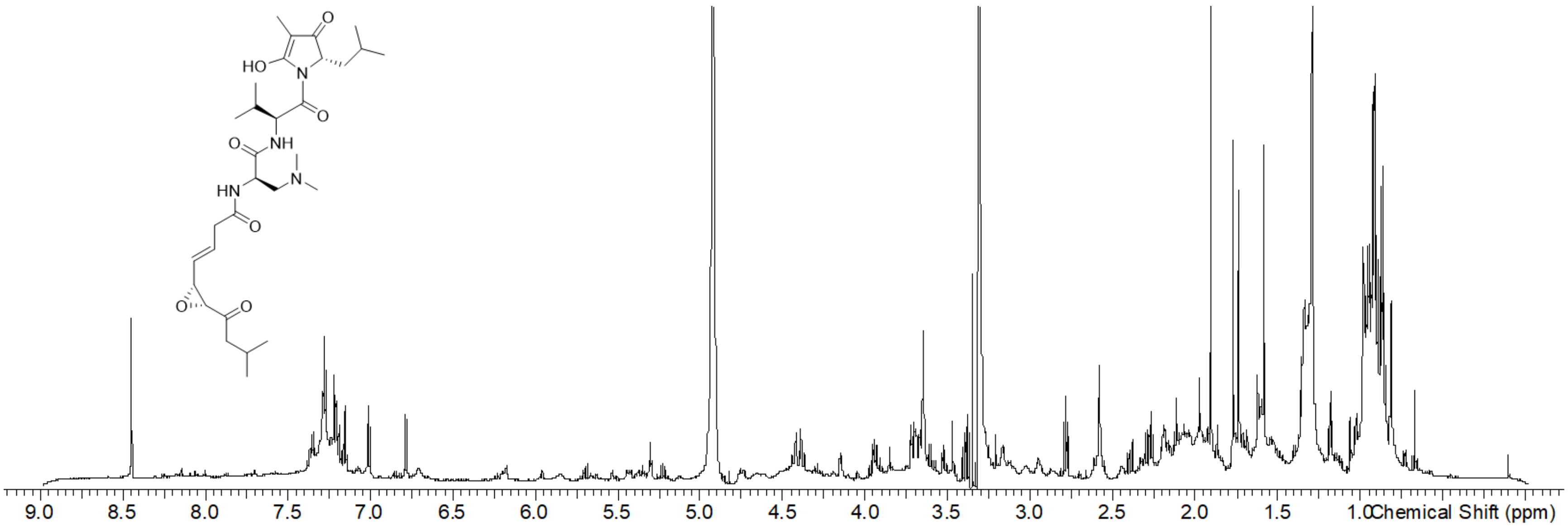

*Figure S 29. [1]H spectrum of sandarazol B in* MeOD-$d_4$.

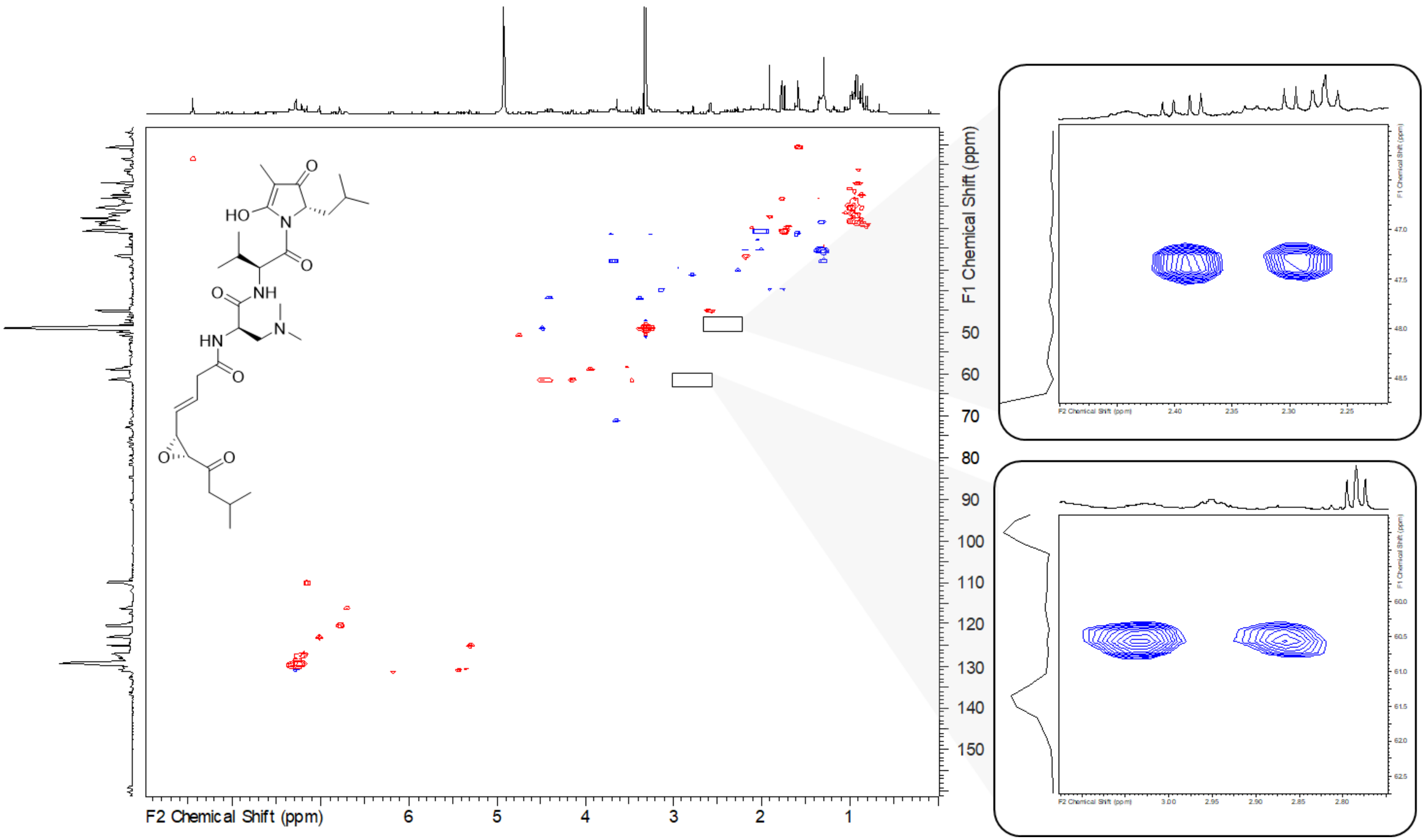

*Figure S 30. HSQC spectrum of sandarazol B in* MeOD-$d_4$. *Right side: Zoom into two methyl groups not visible at the zoom factor of the complete spectrum.*

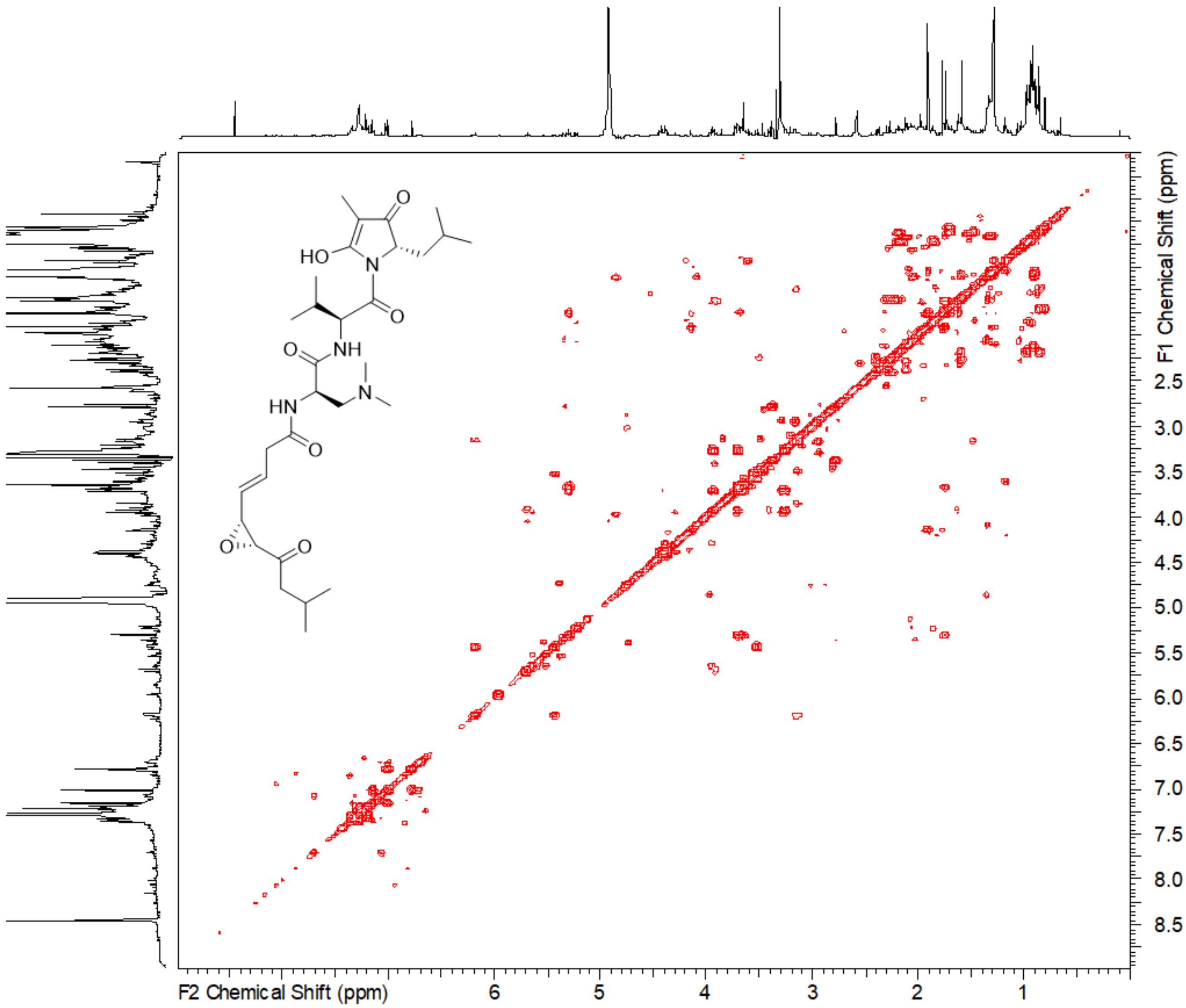

*Figure S 31. COSY spectrum of sandarazol B in* MeOD-$d_4$.

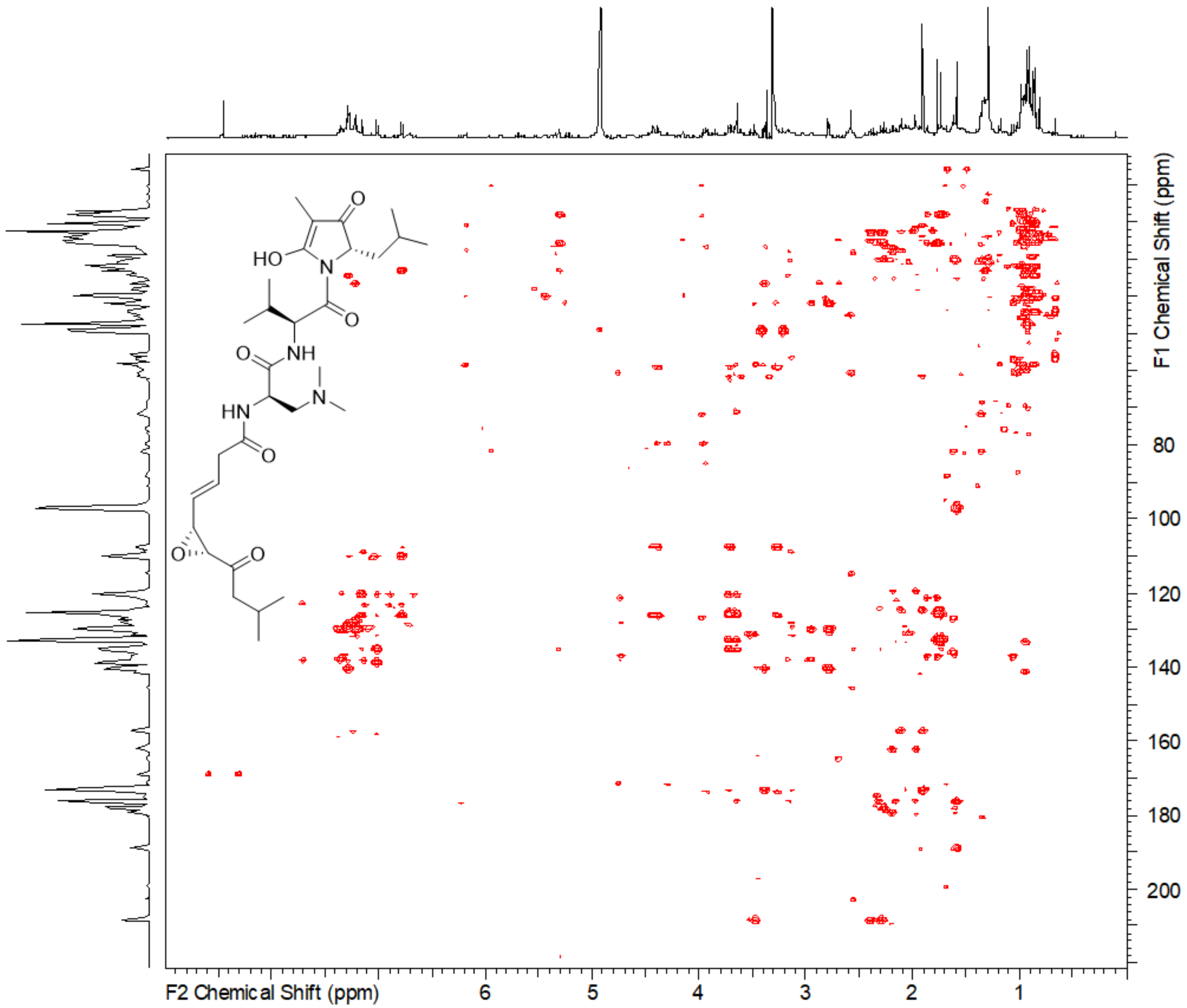

*Figure S 32. HMBC spectrum of sandarazol B in* MeOD-$d_4$.

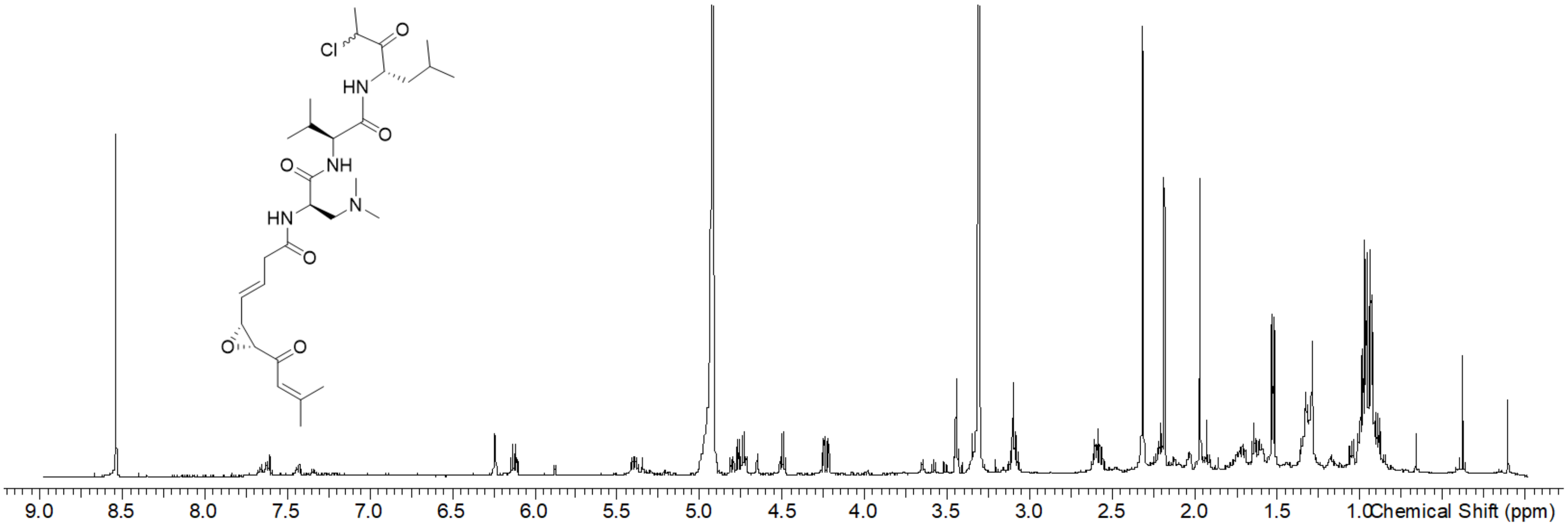

*Figure S 33. [1]H spectrum of sandarazol C in* MeOD-d4.

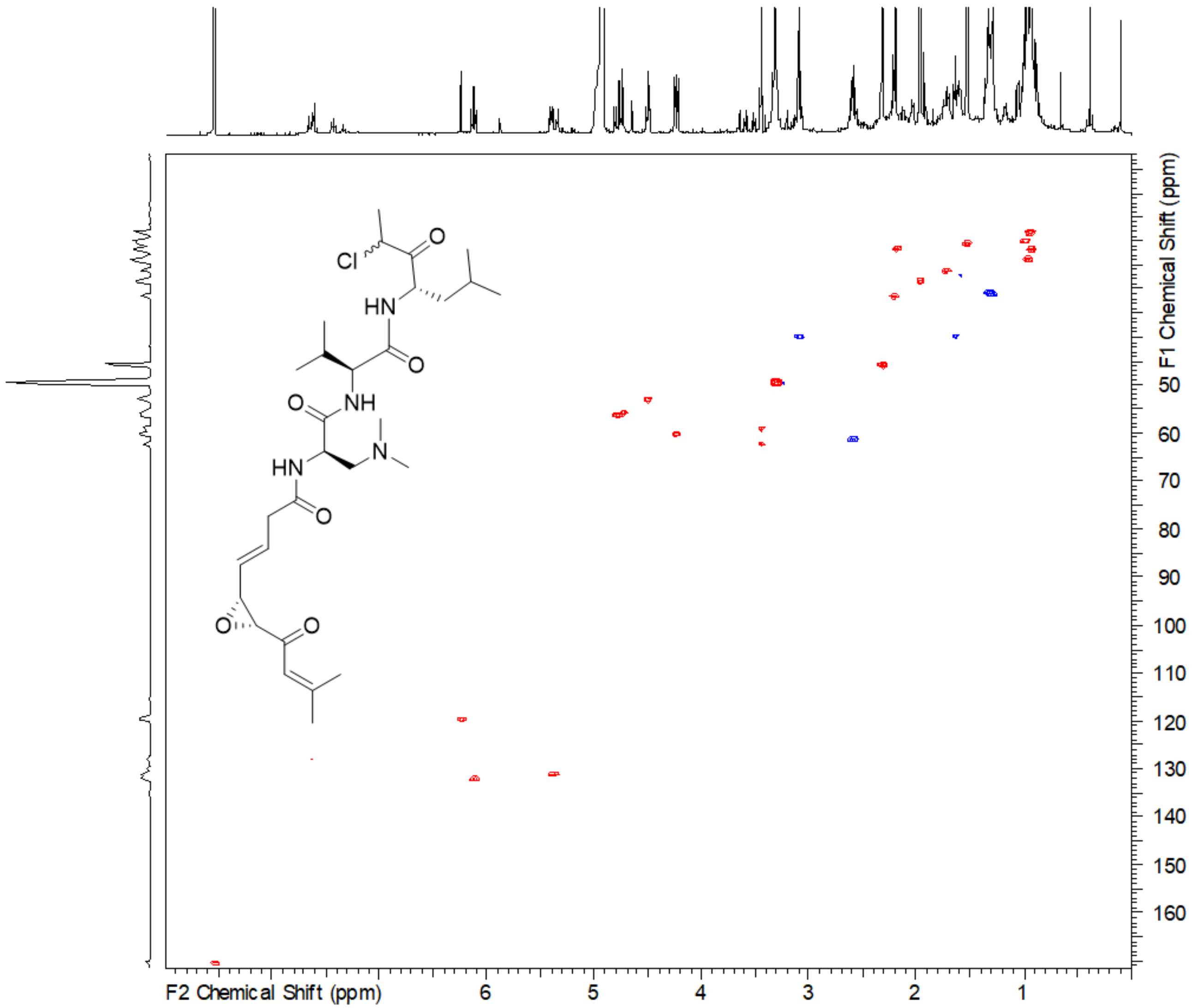

*Figure S 34. HSQC spectrum of sandarazol C in* MeOD-$d_4$.

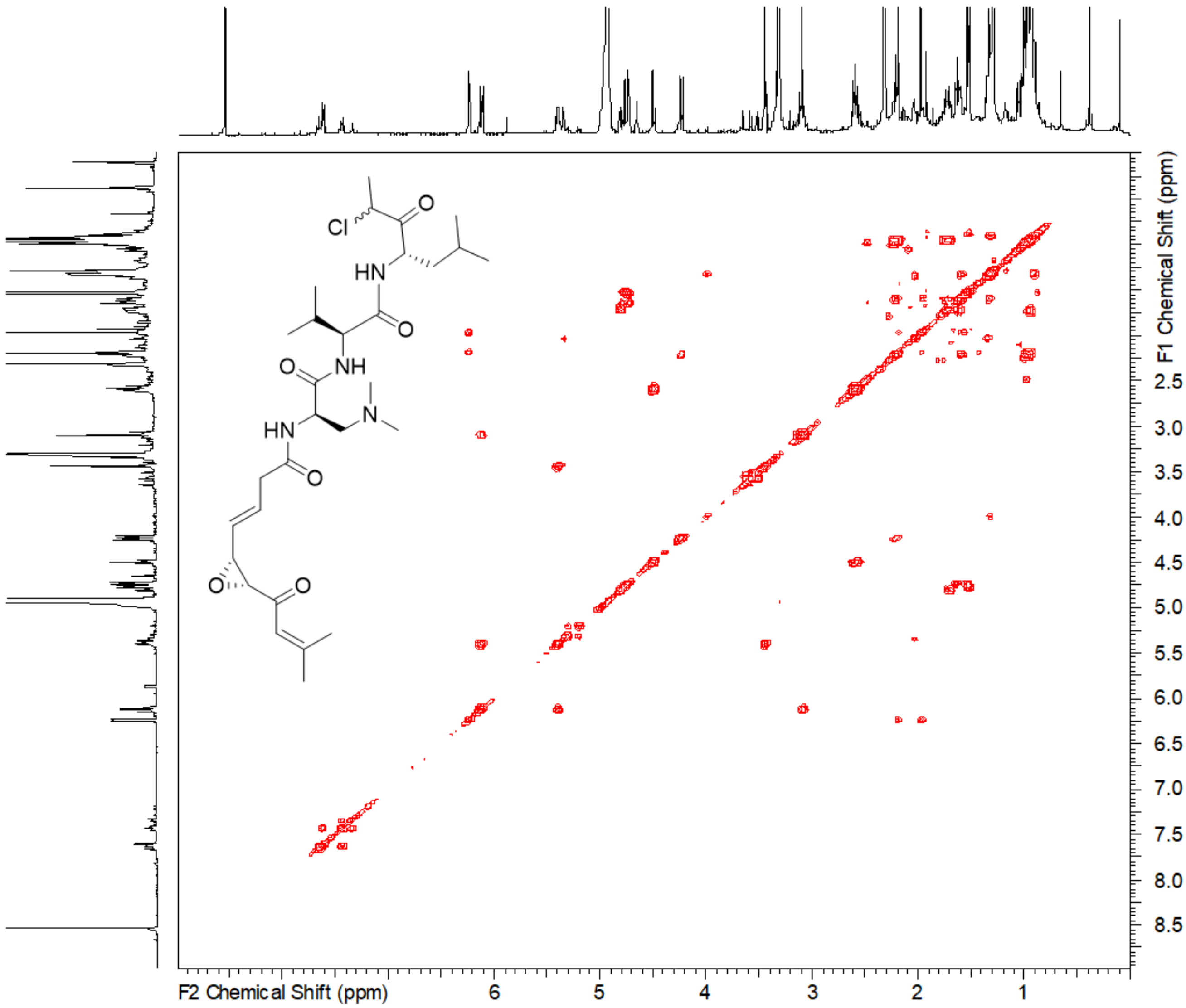

*Figure S 35. COSY spectrum of sandarazol C in* MeOD-d4.

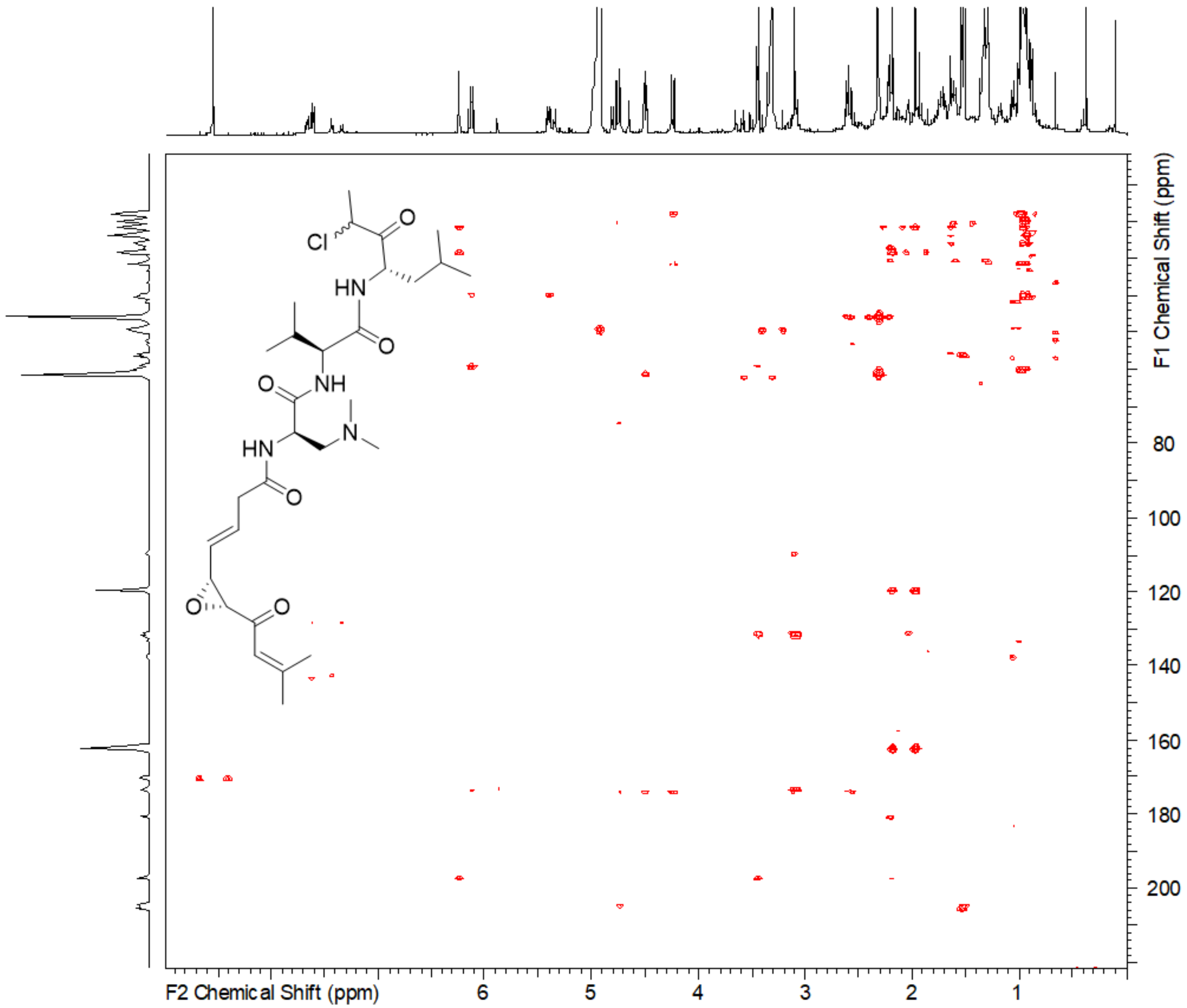

*Figure S 36. HMBC spectrum of sandarazol C in* MeOD-d4.

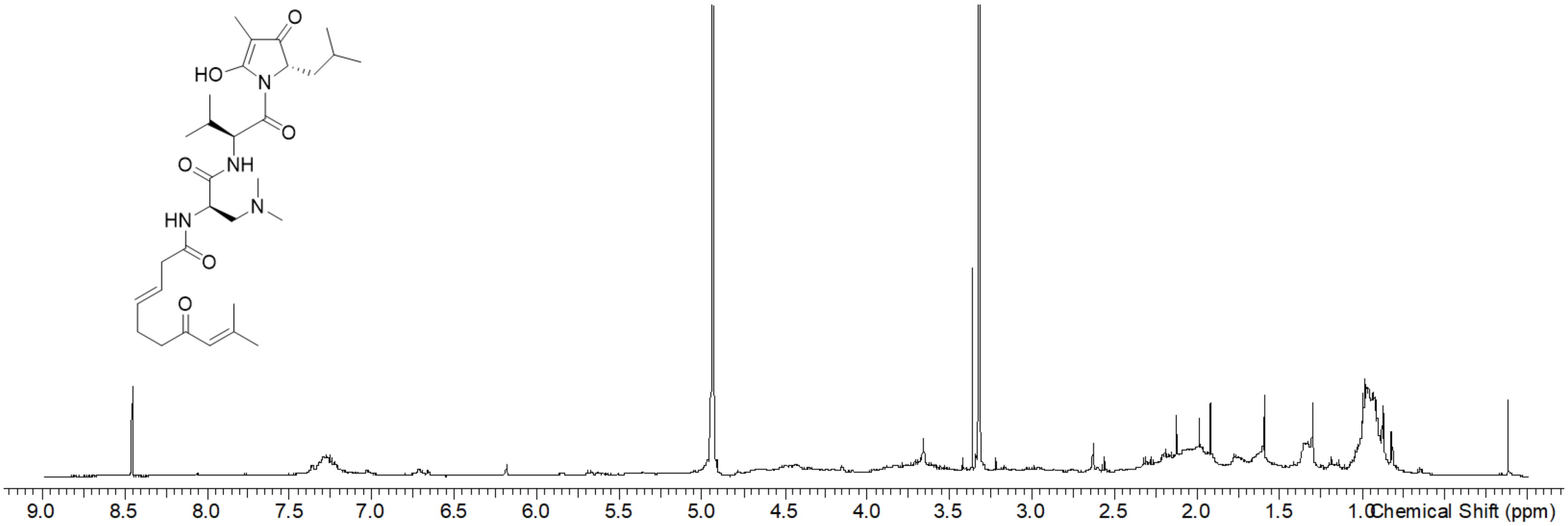

*Figure S 37. [1]H spectrum of sandarazol F in* MeOD-d4.

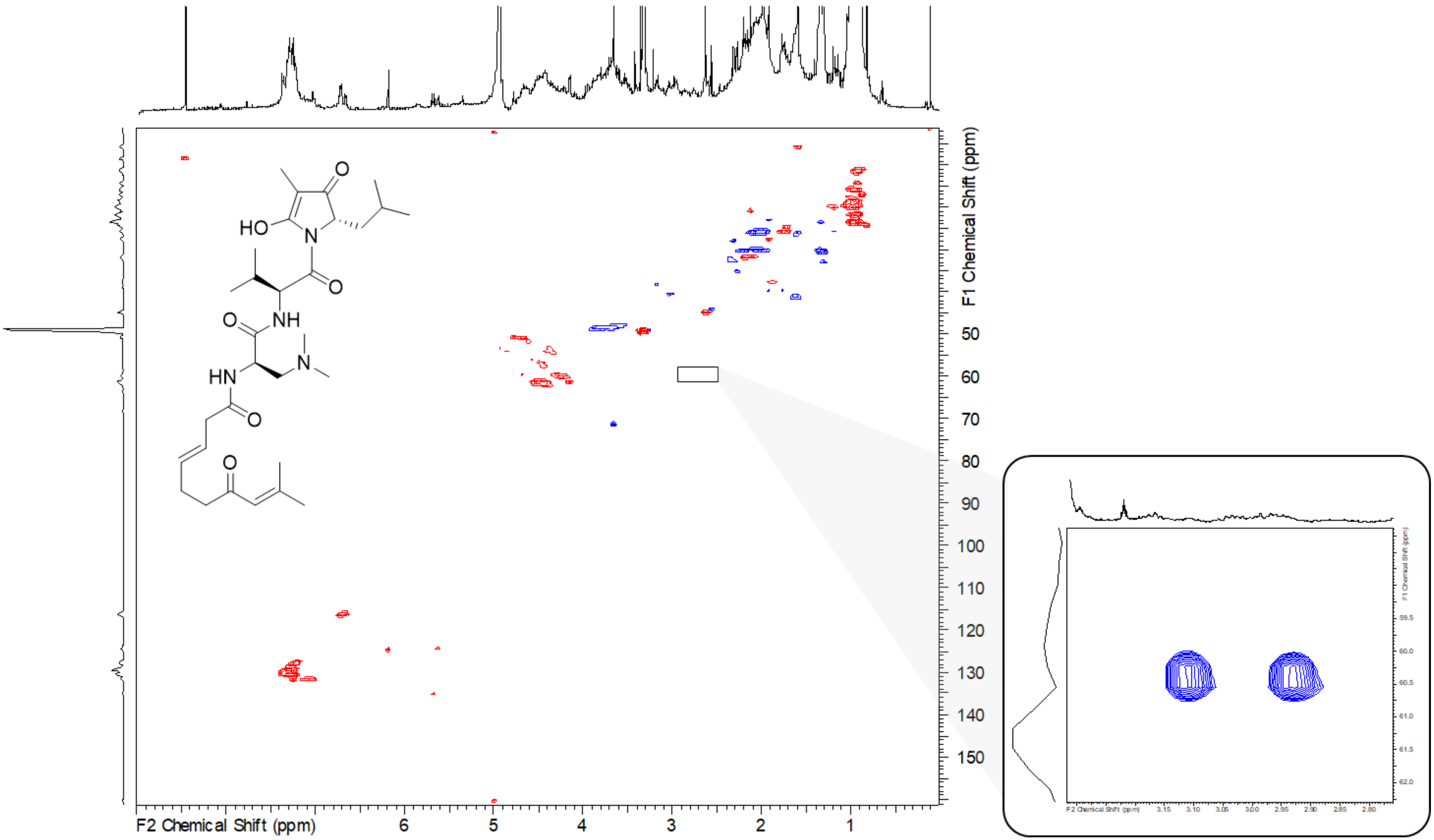

*Figure S 38. HSQC spectrum of sandarazol F in* MeOD-d4. *Right side: Zoom into methyl group not visible at the zoom factor of the complete spectrum.*

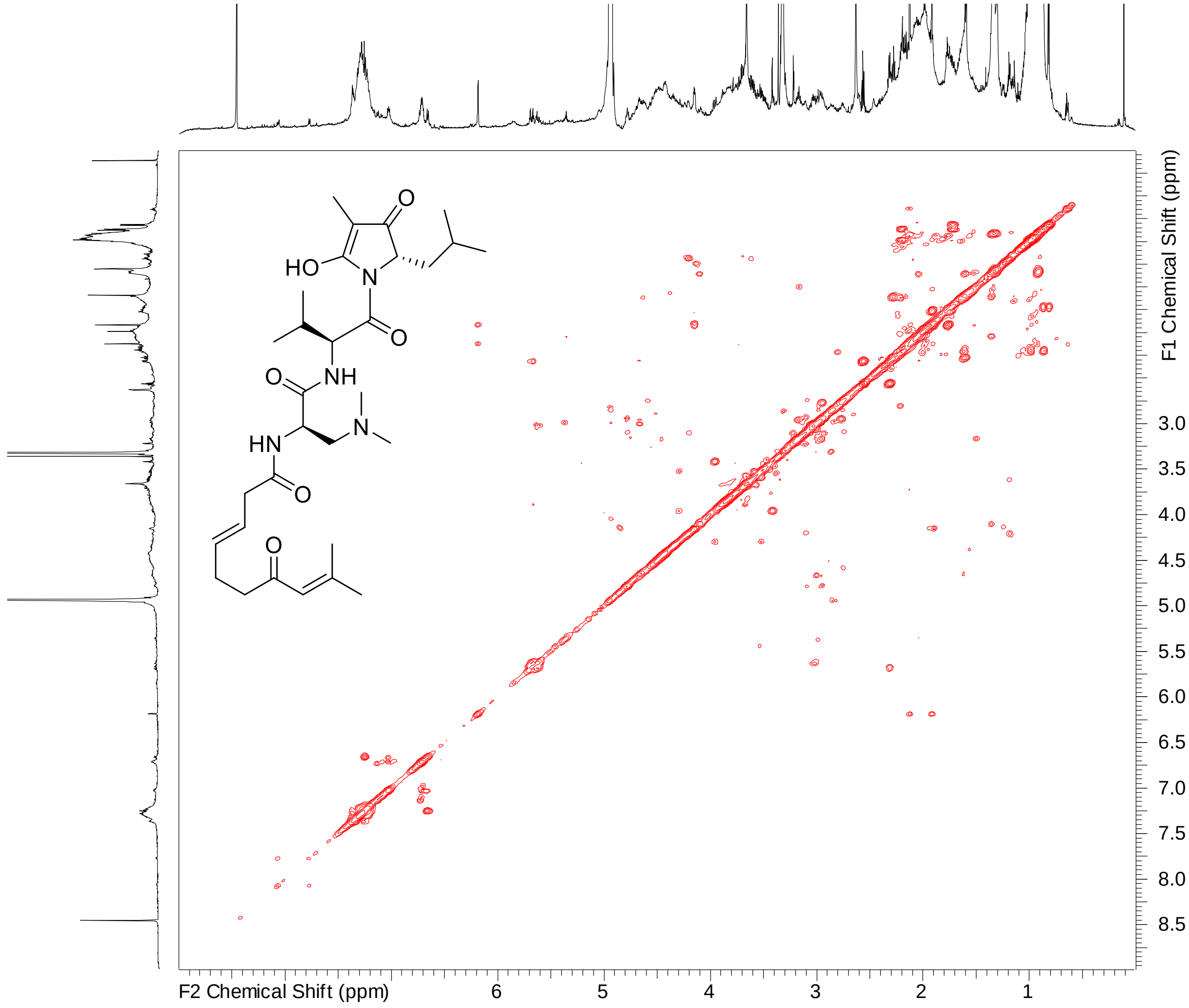

*Figure S 39. . COSY spectrum of sandarazol F in* MeOD-d4.

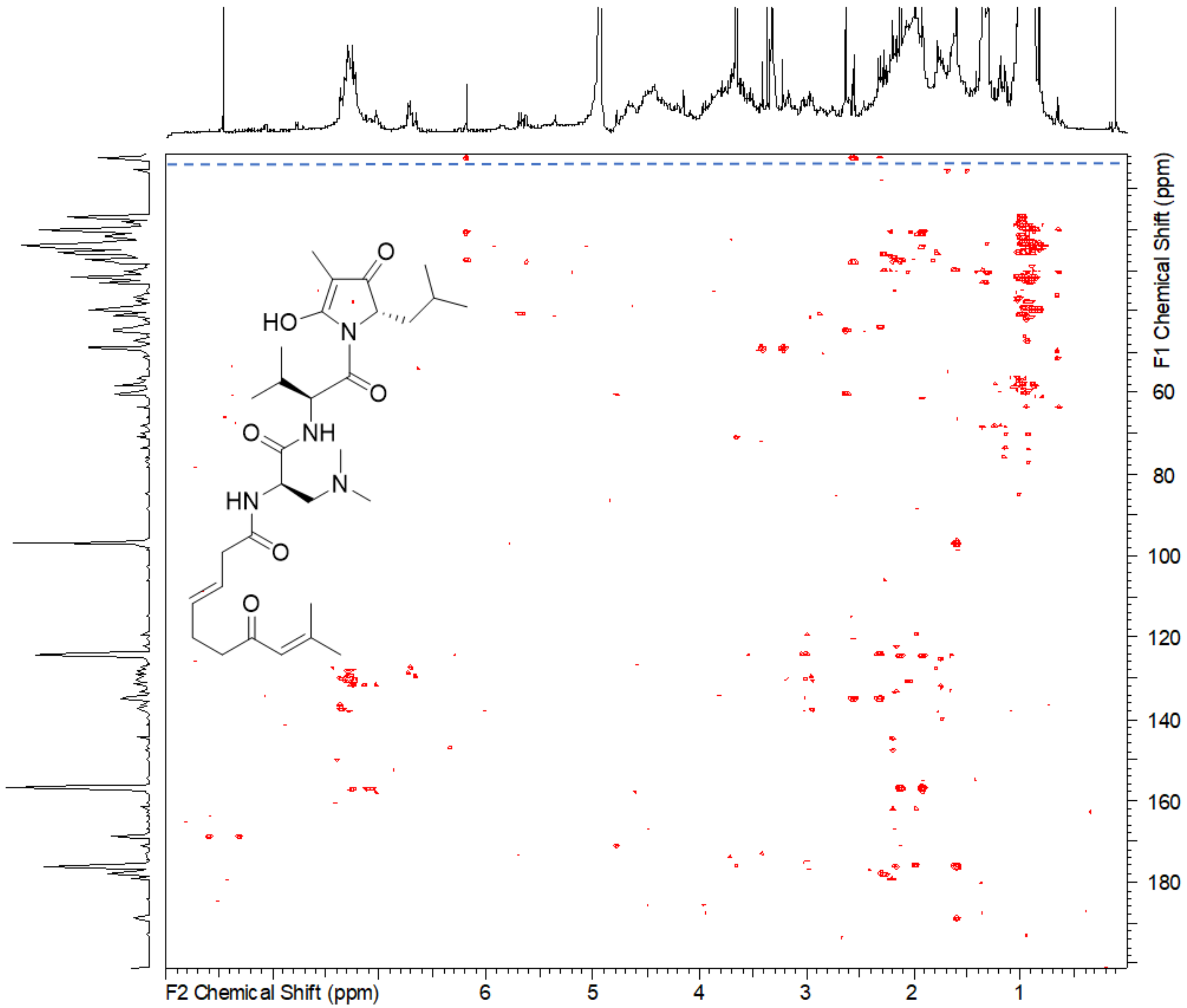

*Figure S 40. HMBC spectrum of sandarazol F in* MeOD-$d_4$. Blue dashed line: Signals exceeding $^{13}C$ 200 ppm, displayed highfield due to limited width of 200 ppm.

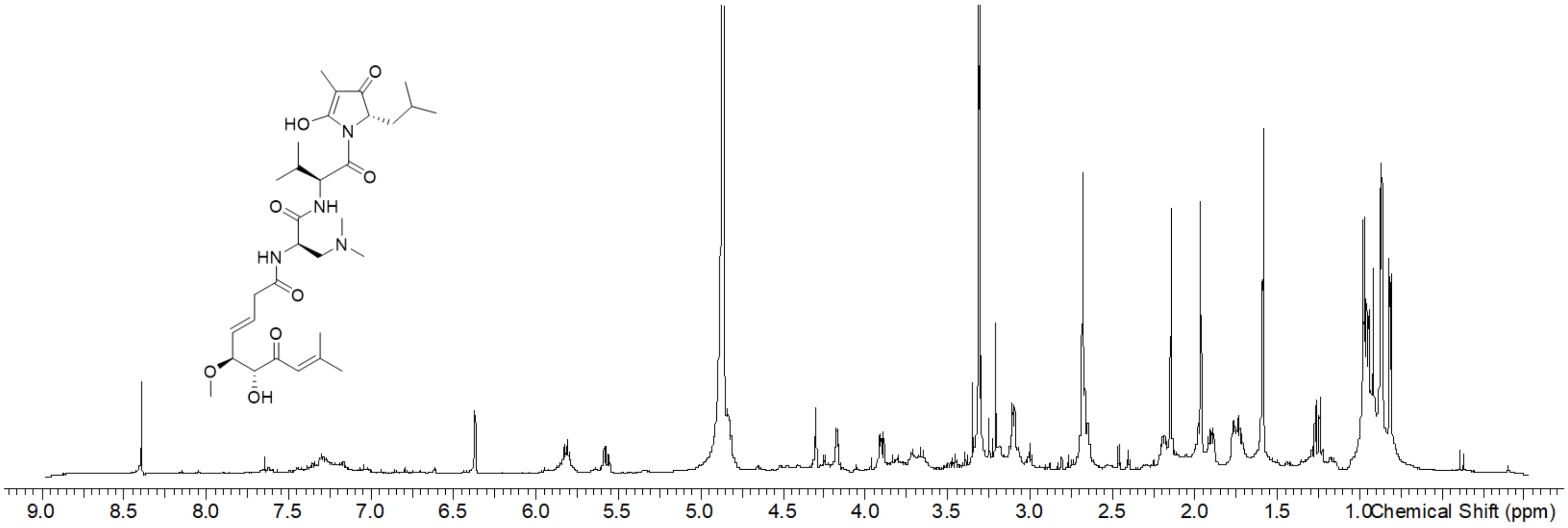

*Figure S 41. $^{1}$H spectrum of methoxy sandarazol A in* MeOD-d$_4$.

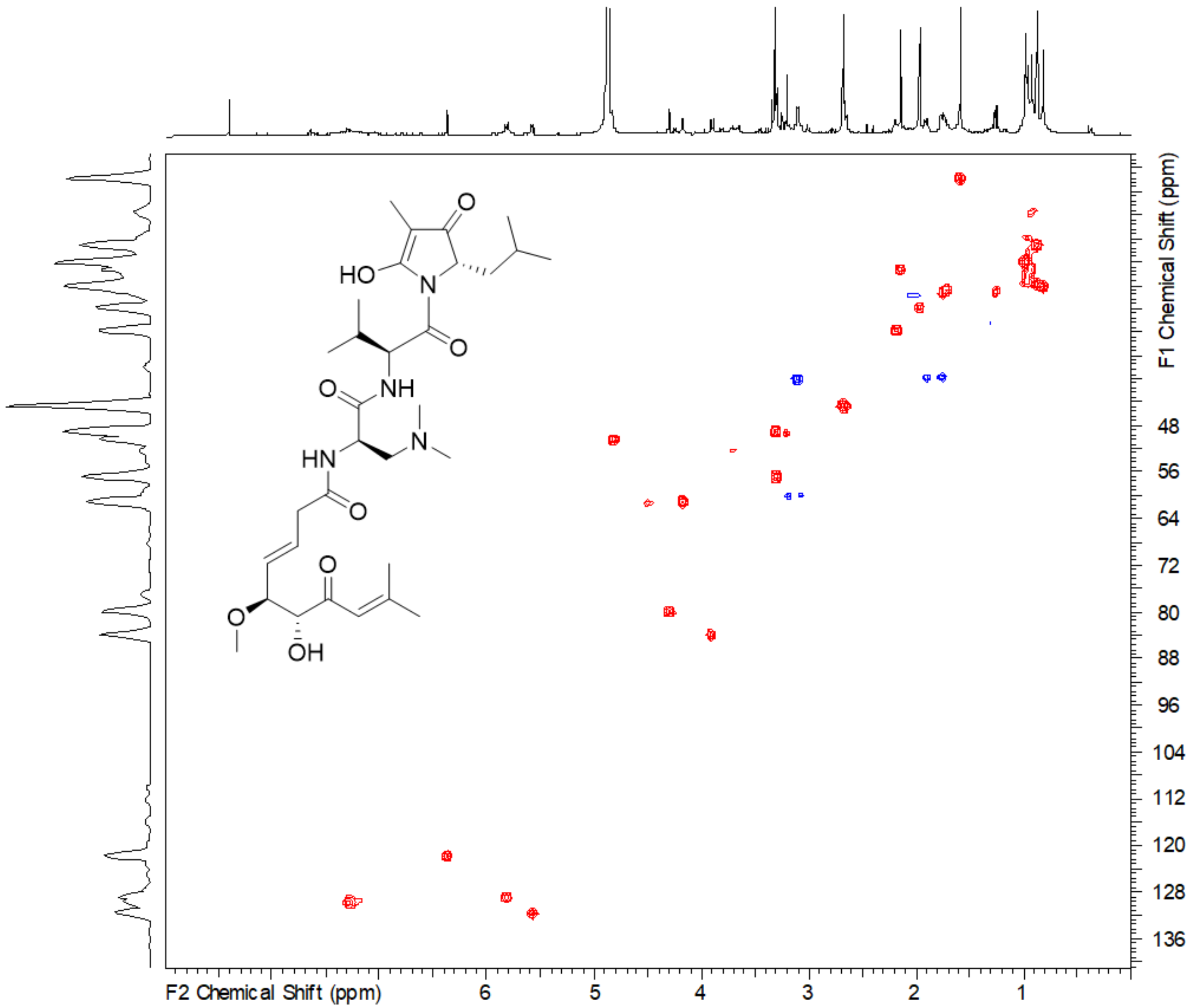

*Figure S 42. HSQC spectrum of methoxy sandarazol A in* MeOD-d4.

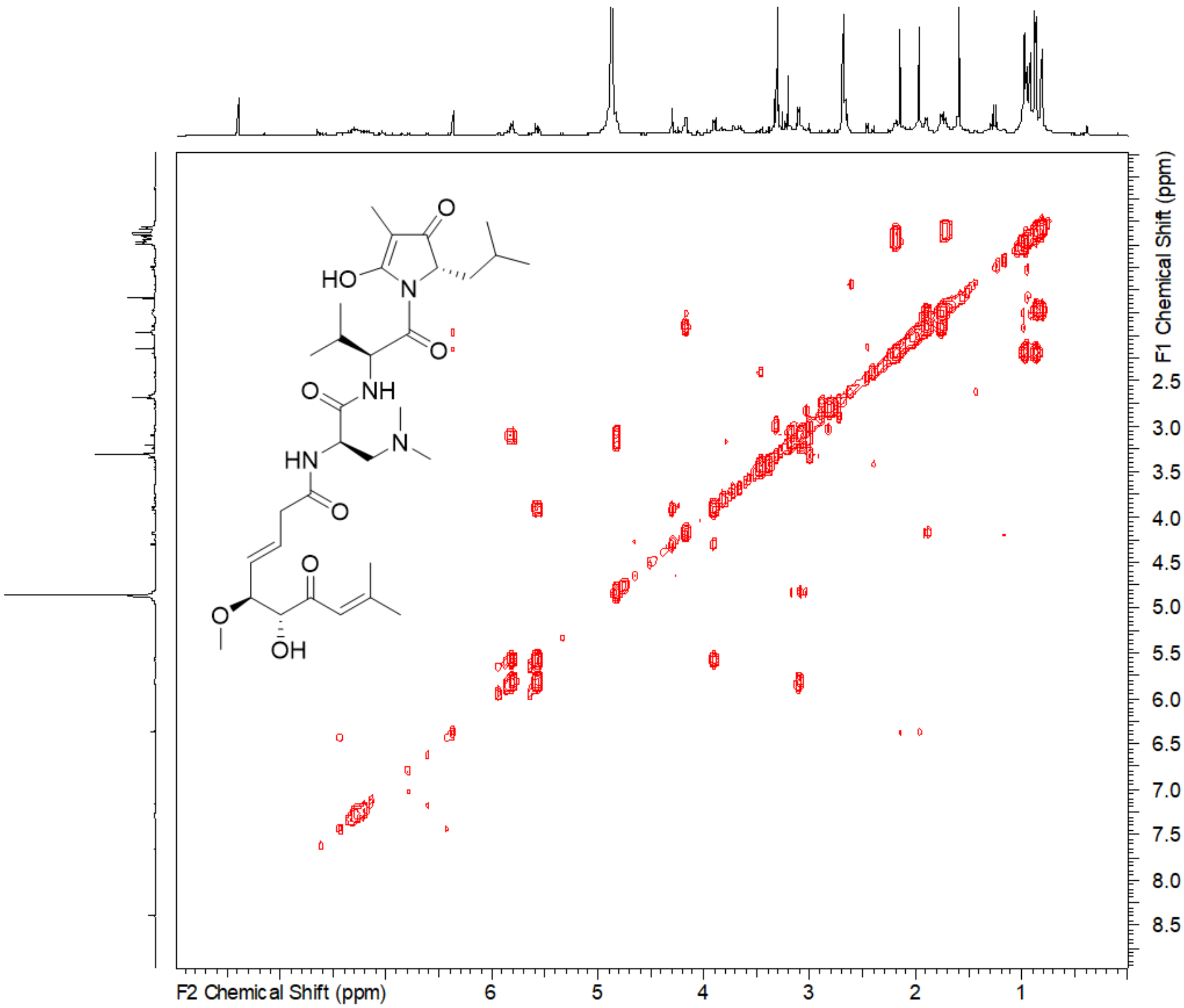

*Figure S 43. COSY spectrum of methoxy sandarazol A in* MeOD-$d_4$.

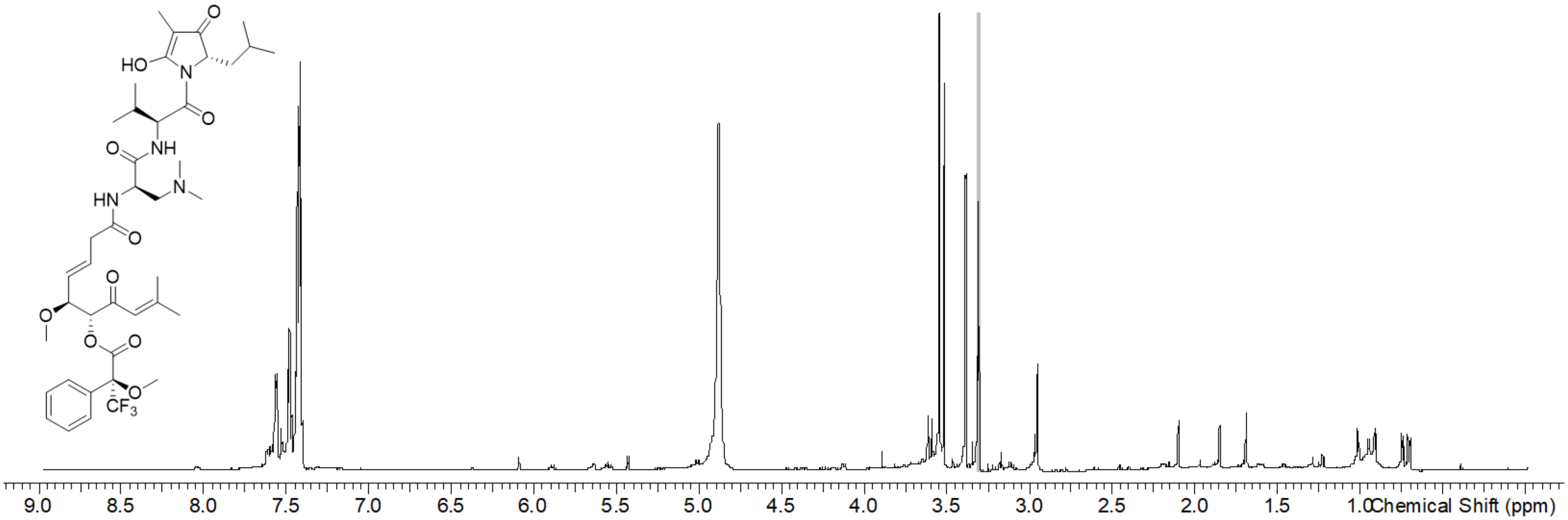

*Figure S 44. [1]H spectrum of methoxy sandarazol A(S)-Mosher ester in* MeOD-d4.

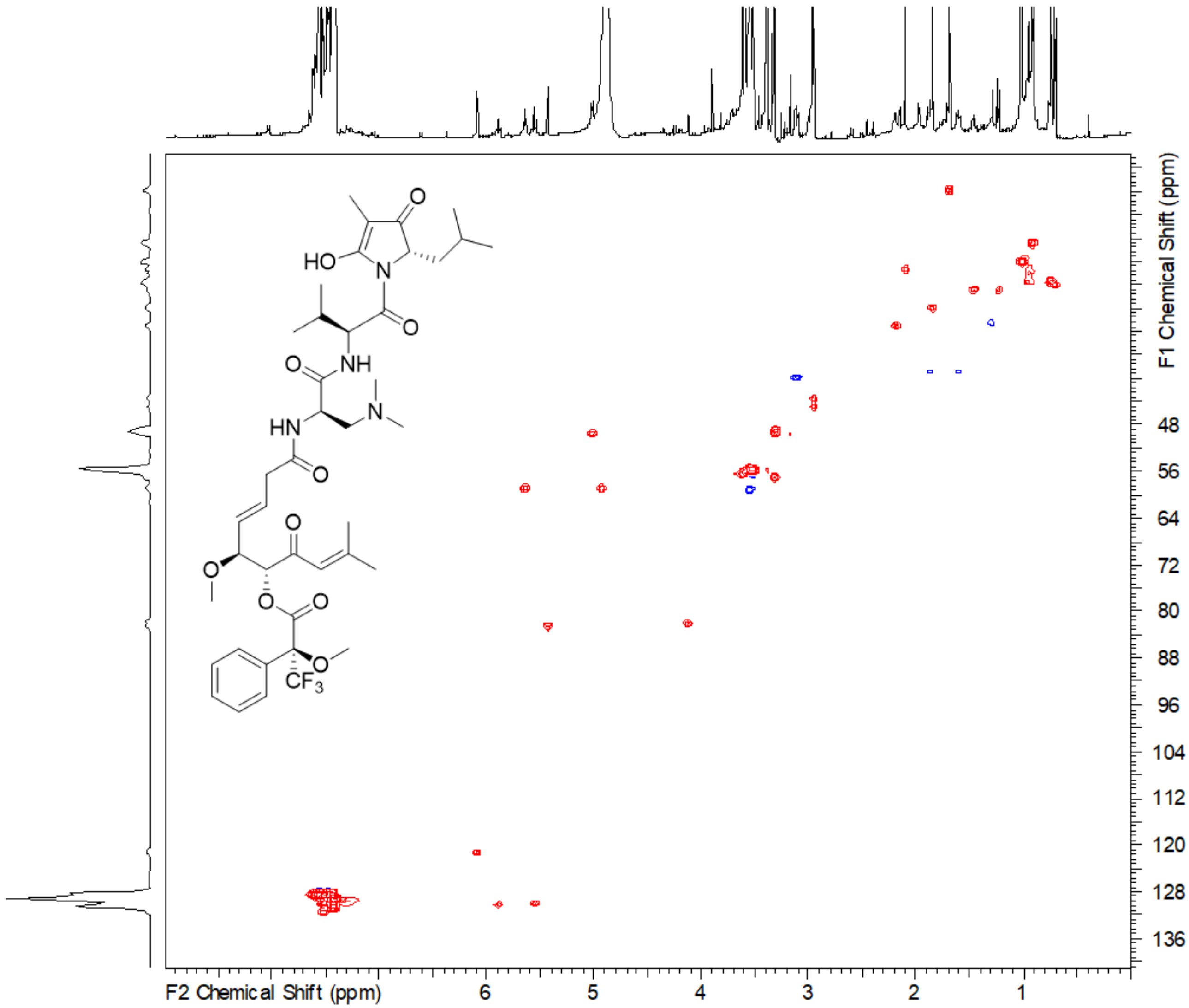

*Figure S 45.HSQC spectrum of methoxy sandarazol A(S)-Mosher ester in* MeOD-d4.

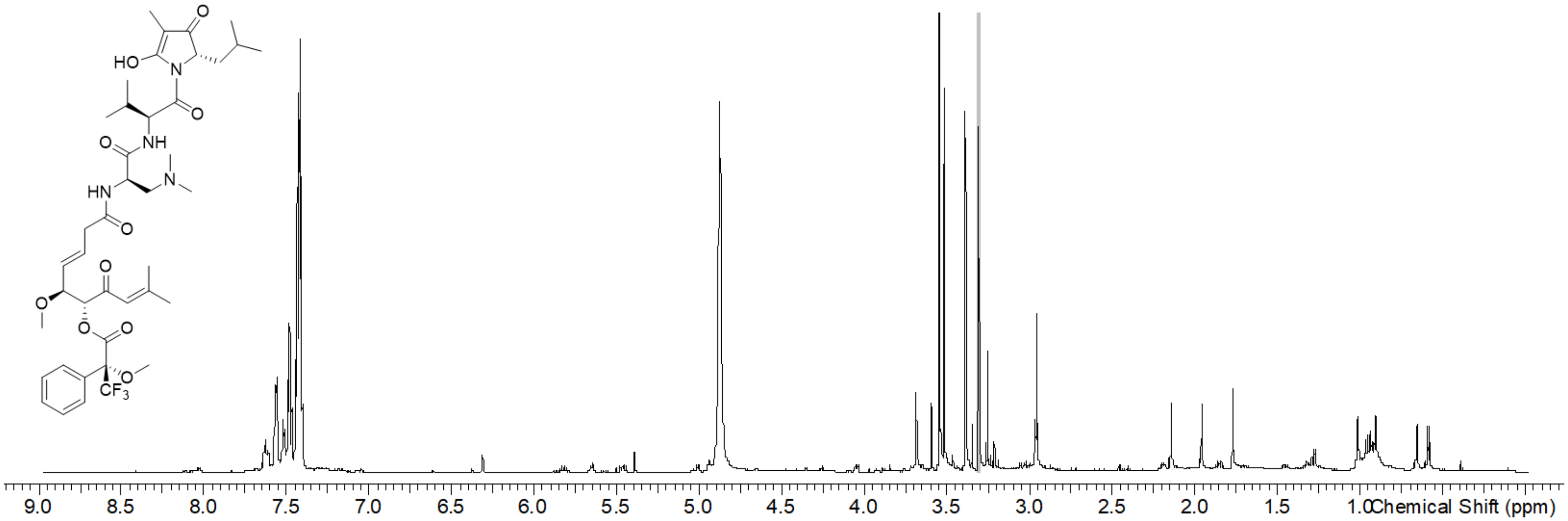

*Figure S 46. [1]H spectrum of methoxy sandarazol A(R)-Mosher ester in* MeOD-d4.

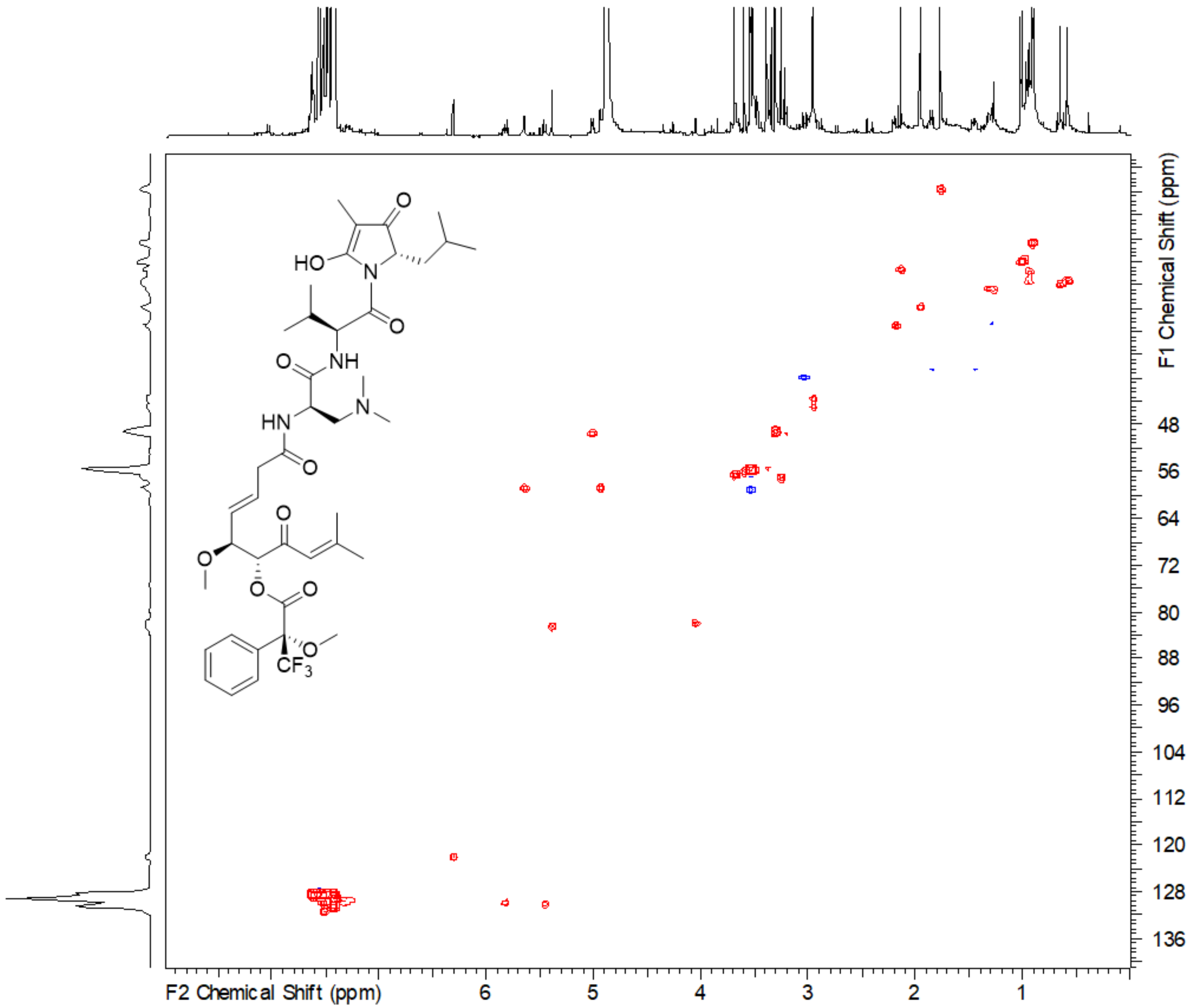

*Figure S 47. HSQC spectrum of methoxy sandarazol A(R)-Mosher ester in* MeOD-d4.

### 8 Biological assay conditions

Human HCT-116 colon carcinoma cells (ACC-581) were received from the German Collection of Microorganisms and Cell Cultures (Deutsche Sammlung für Mikroorganismen und Zellkulturen, DSMZ) and were cultured under the conditions recommended by the depositor. To determine the cytotoxic activities of the sandarazols, cells from actively growing cultures were harvested and seeded at 5 x 104 cells per well in a 96 CELLBind® surface well plate in 120 µL 90% modified McCoy's 5A medium with 10% h.i. fetal bovine serum (FBS). After 2 h of equilibration, the cells were treated with the compounds in a serial dilution. After 5 days of incubation at 37 °C, 20 µL of 5 mg/mL thiazolyl blue tetrazolium bromide (MTT) in PBS was added. After discarding the medium, 100 µL of a 2-propanol 10 N HCl mixture (250:1) was added to dissolve formazan granules. A microplate reader (EL808, Bio-Tek Instruments Inc.) was used to determine the absorbance at 570 nm.

All microorganisms used for the biological assays were obtained from the German Collection of Microorganisms and Cell Cultures (Deutsche Sammlung für Mikroorganismen und Zellkulturen, DSMZ) or were part of our in-house strain collection and were cultured under the conditions recommended by the depositor. Bacterial cultures were prepared in MHB (2.9 g/L beef infusion solids, 17.5 g/L casein hydrolysate, 1.5 g/L starch at pH 7.4), M7H9 (0.5 g/L ammonium sulfate, 2.5 g/L disodium phosphate, 1.0 g/L monopotassium phosphate, 0.1 g/L sodium citrate, 0.05 g/L magnesium sulfate, 0.0005 g/L calcium chloride, 0.001 g/L zinc sulfate, 0.001 g/L copper sulfate, 0.04 g/L ferric ammonium citrate, 0.50 g/L L-Glutamic acid, 0.001 g/L pyridoxine, 0.0005 g/L biotin at pH 6.6) or Myc 2.0 medium inoculated from the strain grown on agar plate. The compounds were diluted serially in sterile 96 well-plates before adding the bacterial cell suspension. The bacteria were grown for 24 h at RT, 30 °C or 37 °C. Growth inhibition was inspected visually. MIC50 values were determined relative to the respective control samples by sigmoidal curve fitting.

(2) Wang, M.; Carver, J. J.; Phelan, V. V.; Sanchez, L. M.; Garg, N.; Peng, Y.; Nguyen, D. D.; Watrous, J.; Kapono, C. A.; Luzzatto-Knaan, T.; Porto, C.; Bouslimani, A.; Melnik, A. V.; Meehan, M. J.; Liu, W.-T.; Crusemann, M.; Boudreau, P. D.; Esquenazi, E.; Sandoval-Calderon, M.; Kersten, R. D.; Pace, L. A.; Quinn, R. A.; Duncan, K. R.; Hsu, C.-C.; Floros, D. J.; Gavilan, R. G.; Kleigrewe, K.; Northen, T.; Dutton, R. J.; Parrot, D.; Carlson, E. E.; Aigle, B.; Michelsen, C. F.; Jelsbak, L.; Sohlenkamp, C.; Pevzner, P.; Edlund, A.; McLean, J.; Piel, J.; Murphy, B. T.; Gerwick, L.; Liaw, C.-C.; Yang, Y.-L.; Humpf, H.-U.; Maansson, M.; Keyzers, R. A.; Sims, A. C.; Johnson, A. R.; Sidebottom, A. M.; Sedio, B. E.; Klitgaard, A.; Larson, C. B.; Boya P, C. A.; Torres-Mendoza, D.; Gonzalez, D. J.; Silva, D. B.; Marques, L. M.; Demarque, D. P.; Pociute, E.; O'Neill, E. C.; Briand, E.; Helfrich, E. J. N.; Granatosky, E. A.; Glukhov, E.; Ryffel, F.; Houson, H.; Mohimani, H.; Kharbush, J. J.; Zeng, Y.; Vorholt, J. A.; Kurita, K. L.; Charusanti, P.; McPhail, K. L.; Nielsen, K. F.; Vuong, L.; Elfeki, M.; Traxler, M. F.; Engene, N.; Koyama, N.; Vining, O. B.; Baric, R.; Silva, R. R.; Mascuch, S. J.; Tomasi, S.; Jenkins, S.; Macherla, V.; Hoffman, T.; Agarwal, V.; Williams, P. G.; Dai, J.; Neupane, R.; Gurr, J.; Rodriguez, A. M. C.; Lamsa, A.; Zhang, C.; Dorrestein, K.; Duggan, B. M.; Almaliti, J.; Allard, P.-M.; Phapale, P.; Nothias, L.-F.; Alexandrov, T.; Litaudon, M.; Wolfender, J.-L.; Kyle, J. E.; Metz, T. O.; Peryea, T.; Nguyen, D.-T.; VanLeer, D.; Shinn, P.; Jadhav, A.; Muller, R.; Waters, K. M.; Shi, W.; Liu, X.; Zhang, L.; Knight, R.; Jensen, P. R.; Palsson, B. O.; Pogliano, K.; Linington, R. G.; Gutierrez, M.; Lopes, N. P.; Gerwick, W. H.; Moore, B. S.; Dorrestein, P. C.; Bandeira, N. Sharing and community curation of mass spectrometry data with Global Natural Products Social Molecular Networking. *Nat. Biotechnol.* **2016**, *34* (8), 828–837. DOI: 10.1038/nbt.3597.

(3) O'Leary, N. A.; Wright, M. W.; Brister, J. R.; Ciufo, S.; Haddad, D.; McVeigh, R.; Rajput, B.; Robbertse, B.; Smith-White, B.; Ako-Adjei, D.; Astashyn, A.; Badretdin, A.; Bao, Y.; Blinkova, O.; Brover, V.; Chetvernin, V.; Choi, J.; Cox, E.; Ermolaeva, O.; Farrell, C. M.; Goldfarb, T.; Gupta, T.; Haft, D.; Hatcher, E.; Hlavina, W.; Joardar, V. S.; Kodali, V. K.; Li, W.; Maglott, D.; Masterson, P.; McGarvey, K. M.; Murphy, M. R.; O'Neill, K.; Pujar, S.; Rangwala, S. H.; Rausch, D.; Riddick, L. D.; Schoch, C.; Shkeda, A.; Storz, S. S.; Sun, H.; Thibaud-Nissen, F.; Tolstoy, I.; Tully, R. E.; Vatsan, A. R.; Wallin, C.; Webb, D.; Wu, W.; Landrum, M. J.; Kimchi, A.; Tatusova, T.; DiCuccio, M.; Kitts, P.; Murphy, T. D.; Pruitt, K. D. Reference sequence (RefSeq)

database at NCBI: current status, taxonomic expansion, and functional annotation. *Nucleic acids research* **2016**, *44* (D1), D733-45. DOI: 10.1093/nar/gkv1189.

(4) Pogorevc, D.; Panter, F.; Schillinger, C.; Jansen, R.; Wenzel, S. C.; Müller, R. Production optimization and biosynthesis revision of corallopyronin A, a potent anti-filarial antibiotic. *Metab. Eng.* **2019**, *55*, 201–211. DOI: 10.1016/j.ymben.2019.07.010.

(5) Chang, Z.; Sitachitta, N.; Rossi, J. V.; Roberts, M. A.; Flatt, P.; Jia, J.; Sherman, D. H.; Gerwick, W. H. Biosynthetic pathway and gene cluster analysis of Curacin A, an antitubulin natural product from the tropical marine cyanobacterium *Lyngbya majuscula*. *J. Nat. Prod.* **2004**, *67*, 1356–1367. DOI: 10.1021/np0499261.

(6) Panter, F.; Krug, D.; Baumann, S.; Müller, R. Self-resistance guided genome mining uncovers new topoisomerase inhibitors from myxobacteria. *Chem. Sci.* **2018**, *9* (21), 4898–4908. DOI: 10.1039/C8SC01325J.

(7) Marfey's reagent: Past, present, and future uses of 1-fluoro-2,4-dinitrophenyl-5-L-alanine amide.

(8) Morrison, E.; Kantz, A.; Gassner, G. T.; Sazinsky, M. H. Structure and mechanism of styrene monooxygenase reductase: new insight into the FAD-transfer reaction. *Biochemistry* **2013**, *52* (35), 6063–6075. DOI: 10.1021/bi400763h.

(9) Sullivan, G. R.; Dale, J. A.; Mosher, H. S. Correlation of configuration and fluorine-19 chemical shifts of .alpha.-methoxy-.alpha.-trifluoromethylphenyl acetate derivatives. *J. Org. Chem.* **1973**, *38* (12), 2143–2147. DOI: 10.1021/jo00952a006.

(10) Hoye, T. R.; Jeffrey, C. S.; Shao, F. Mosher ester analysis for the determination of absolute configuration of stereogenic (chiral) carbinol carbons. *Nat. Protoc.* **2007**, *2* (10), 2451–2458. DOI: 10.1038/nprot.2007.354.
