## Supplementary figures and images for "The sandarazols are cryptic and structurally unique plasmid encoded toxins from a rare myxobacterium"

### thumb.png

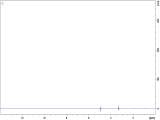

### thumb.png

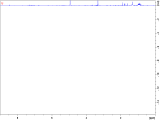

### thumb.png

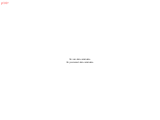

### thumb.png

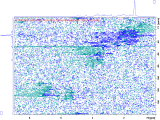

### thumb.png

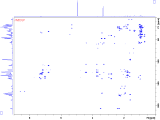

### thumb.png

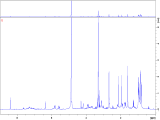

### thumb.png

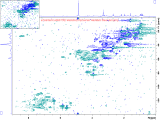
